# Transcriptional Atlas of Ileal-Anal Pouch Immune Cells from Ulcerative Colitis Patients

**DOI:** 10.1101/2020.07.31.231308

**Authors:** Joseph C. Devlin, Jordan Axelrad, Ashley M. Hine, Shannon Chang, Suparna Sarkar, Jian-Da Lin, Kelly V. Ruggles, David Hudesman, Ken Cadwell, P’ng Loke

## Abstract

How the human intestinal immune system is distinctly organized to respond to inflammation is still poorly understood. Here, we used single-cell RNA-sequencing to examine lamina propria CD45+ hematopoietic cells from patients with inflammatory bowel disease that have undergone ileal pouch-anal anastomosis, or the colon mucosa of ulcerative colitis patients. We identified a population of *IL1B*+ antimicrobial macrophages and *FOXP3*/+*BATF*+ T cells that are associated and expanded in inflamed tissues, which we further validated in other scRNA-seq datasets from IBD patients. CD8+ T cells were unexpectedly more abundant in the pouch than colon. Cell type specific markers obtained from single-cell RNA-sequencing was used to infer representation from bulk RNA sequencing datasets, which further implicated antimicrobial macrophages expressing *IL1B* with *S100A8/A9* calprotectin as being associated with inflammation, as well as *Bacteroides* and *Escherichia* bacterial species. Finally, we find that non-responsiveness to anti-integrin biologic therapies in UC patients is associated with the signature of this antimicrobial macrophage population in a subset of patients. This study identified conserved and distinct features of intestinal inflammation between parts of the small and large intestine undergoing similar inflammation conditions.

## INTRODUCTION

The intestinal immune system is organized distinctly between anatomically defined segments that have different physiological functions. Whereas immune cells in the small intestine protect the epithelium from infection to enable nutrient absorption while maintaining tolerance to dietary antigens, the large intestine must maintain a détente with the large number of commensal bacteria without triggering overt inflammation. This fine balance breaks down in the context of inflammatory bowel diseases (IBD) such as ulcerative colitis (UC) and Crohn’s disease (CD), whereby inflammatory damage to the epithelium results in mucosal ulceration causing diarrhea, bleeding, and abdominal pain. Restorative proctocolectomy with ileal pouch-anal anastomosis (IPAA) is a common surgical procedure in UC patients with medically refractory disease^1^. The ileal pouch reservoir, or J-pouch, is a novel organ created from small intestine formed into a J-shaped pouch that restores intestinal continuity and replaces the function of the large intestine. Unfortunately, nearly 50% of UC patients who undergo this surgery will develop *de novo* intestinal inflammation in the pouch, or pouchitis^2^. Symptoms of pouchitis mimic IBD, including urgency, diarrhea, bleeding, and abdominal pain. The etiology of pouchitis is not known and there are no FDA approved therapies. Treatment often entails chronic antibiotics and restarting immunosuppressive IBD therapies. A better understanding of the inflammatory response for this condition, as well as the anatomically distinct features of intestinal inflammation in the large intestine compared to the pouch, may provide new insights for the management of this disease.

Single-cell RNA sequencing (scRNA-seq) has enabled us to characterize tissue states at a high resolution to better understand human diseases. Surveys of the intestinal tissues have identified unknown subtypes of intestinal epithelial cells and cellular inflammation modules that predict treatment responsiveness^3–5^. In the field of IBD, this approach is providing new insights into these complex diseases and may provide opportunities to define new drug targets, therapeutic strategies and more personalized treatment options^6^. In support of region-specific properties of the intestinal epithelium, small intestinal organoids but not colonic organoids derived from individuals homozygous for the Crohn’s disease variant of *ATG16L1* are susceptible to TNF-α and respond to drugs targeting JAK/STAT and necroptosis signaling^7,8^. Although these findings highlight the importance of epithelial-intrinsic factors in determining the anatomical site affected in IBD subtypes, identifying differences in immune cells between the large and small intestine may also be necessary for explaining disease presentations specific to these regions.

Detailed analysis of lymphocyte differentiation in mice have shown that peripheral regulatory T cells (pTregs) and Th17 cell numbers are differentially regulated in these two anatomical sites in response to region-specific exposure or sampling of food antigens, metabolites, and microbes^9–12^. Homing signals, structural features inherent to the organ, and compartmentalized draining by lymph nodes can also contribute to the localization and function of immune cell subsets^13,14^. It is unclear whether the inflamed pouch harbors immune infiltrates similar to the inflamed colon, the organ it functionally replaces, or if it retains immune characteristics of the small intestinal origin for this new organ. A previous study using bulk RNA-Seq showed that, in addition to acquiring markers of the colon, the pouch is enriched for transcripts related to IL-17 signaling and dendritic cell maturation when compared with the small intestine^15^. Therefore, a detailed survey of immune cells may provide novel insight into mechanisms of inflammation and reveal unique therapeutic interventions.

Here, we utilized scRNA-seq to examine the immune cell types and their activation states from intestinal biopsies collected from the J-pouch as well as the colon of UC patients. Our goal was to identify shared and distinctive features of inflammation in the colon and J-pouch. By focusing on CD45+ leukocytes, we can obtain greater resolution of the immune cell landscape and utilize this data to generate cell type specific transcriptional signatures to deconvolute bulk RNA datasets. This enabled us to implicate an IL-1B signature in macrophages as being associated with inflammation and unresponsiveness to anti-integrin biologic therapies in UC patients.

## RESULTS

### Immune cell transcriptional profiling landscape of intestinal biopsies from the J-pouch and colon of ulcerative colitis (UC) patients

To minimize batch effects and streamline processing, we first established a cryopreservation pipeline (See Methods) to store and analyze leukocytes from intestinal biopsy samples of UC patients (Supplementary Figure 1). Optimization of the freezing medium in particular enabled a robust yield of live immune cells that could be sorted on the flow cytometer for high quality transcriptome analysis (See Methods). Cells from all patient samples were sorted for CD45+ surface expression and sequenced by scRNA-seq on the 10X platform. Nearly 56,000 CD45+ cells were obtained from scRNA-seq of 26 frozen biopsies from inflamed ulcerative colitis (18,375 cells, *n*=11 samples), inflamed pouchitis (20,678 cells, *n*=10 samples) and uninflamed pouch (16,678 cells, *n*=5 samples). Two samples from inflamed UC patients were removed due to low quality of the sequencing reaction. Single cell transcriptomes from all samples were normalized and merged with Seurat version 3^16^ (Supplementary Figure 2A). Over half of the profiled cells, 30,863 cells, were from T cell subsets, 21,324 cells from B cell subsets and the remaining 3,747 cells were myeloid subsets (Fig. 1A). From these 3 lineages we determined 6 major populations of cells including T cells, germinal center/follicular cells, plasma cells, cycling B cells, mast cells and monocyte/macrophage cells. These populations were defined by specific immune cell markers such as *CD3D, CD8A* and *CD4* for T cells, *BANK1, CD19* and *VPREB3* for GC/Follicular cells, *STMN1, MKI67* and *HMGB2* for cycling B cells, *MZB1, XBP1* and *DERL3* for plasma cells, *IL1B, LYZ* and *IL8* for Monocytes/Macrophages and *KIT, CPA3* and *CD9* for mast cells (Fig. 1B,D). The relative percentages of these 6 major populations per patient sample did not separate inflamed or uninflamed samples or pouch and UC samples by principle component analysis (PCA) (Supplementary Figure 2B-C), indicating that there are not major differences in the immune cell landscape between UC colon and pouch samples. When we examined inter-individual variation in relative percentages of the 6 major populations, there was a significantly greater (*p*<0.01) proportion of monocyte/macrophages in pouchitis and UC patient samples compared to the uninflamed pouch (Fig. 1C). This indicated that the inflammatory condition was most consistently driven by differences in the myeloid cell compartment, especially in the monocyte/macrophage population.

**Figure 1.**
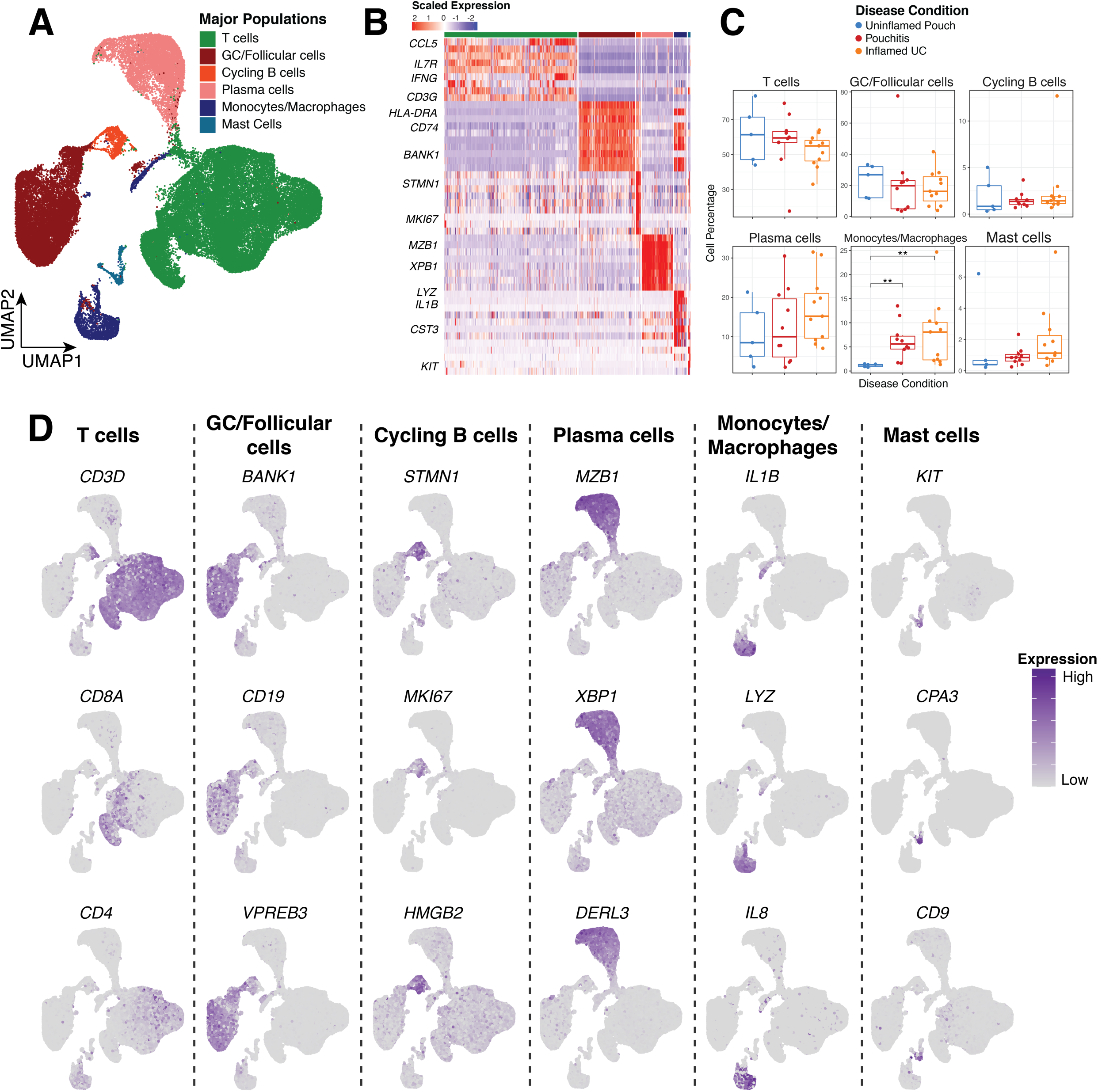
Immune cell landscape of frozen biopsy specimens obtained from the ileal-anal pouch and colon of ulcerative colitis patients. (A) Census of the major immune cell clusters and visualization by UMAP. (B) Heatmap of scaled expression profiles of the major immune cell clusters (C) Boxplots showing cell cluster frequency of the major populations as a percentage of total cells of each patient shown as individual datapoints, compared between patient groups. (D) Feature plots showing representative UMAP visualizations of normalized expression for marker genes enriched in the major cell types. Asterisks indicate significance testing for Wilcoxon ranked test, * = *p* < 0.05, ** = *p* < 0.01, *** = *p* < 0.001.

### Three monocyte/macrophage populations that are increased in inflamed patient samples

Further analysis of the myeloid cell clusters indicated at least 5 minor clusters of cells including three distinct types of monocyte/macrophages, mast cells and a small population of plasmacytoid dendritic cells (Fig. 2A). Differential analysis identified specific immune cell markers for each of these populations including *SOX4, MAFA* and *IERL5* (*SOX4*+/*MAFA*+), *IL1B, LYZ* and *S100A9* (*IL1B*+/*LYZ*+), *APOE, C1QA* and *DNASEIL3* (*APOE*+/*C1QC*+), *CCDC50, IRF4* and *PLAC8* (pDCs) and *KIT, CD69* and *CLU* (Mast Cells) (Fig. 2B). These markers led us to refer to the monocyte/macrophage populations as *SOX4*+/*MAFA*+, *IL1B*+/*LYZ*+ and *APOE*+/*C1QC*+ from here on. *SOX4*+/*MAFA*+ and *IL1B*+/*LYZ*+ Monocyte/macrophage populations were more abundant in patients with both inflamed UC and pouches compared to uninflamed pouches (p<0.05) (Fig. 2C). *APOE*+/*C1QC*+ Monocyte/macrophages and Mast cells had no significant differences (Fig. 2C). pDCs were not detected in uninflamed pouches and are small in number (Fig. 2C).

**Figure 2.**
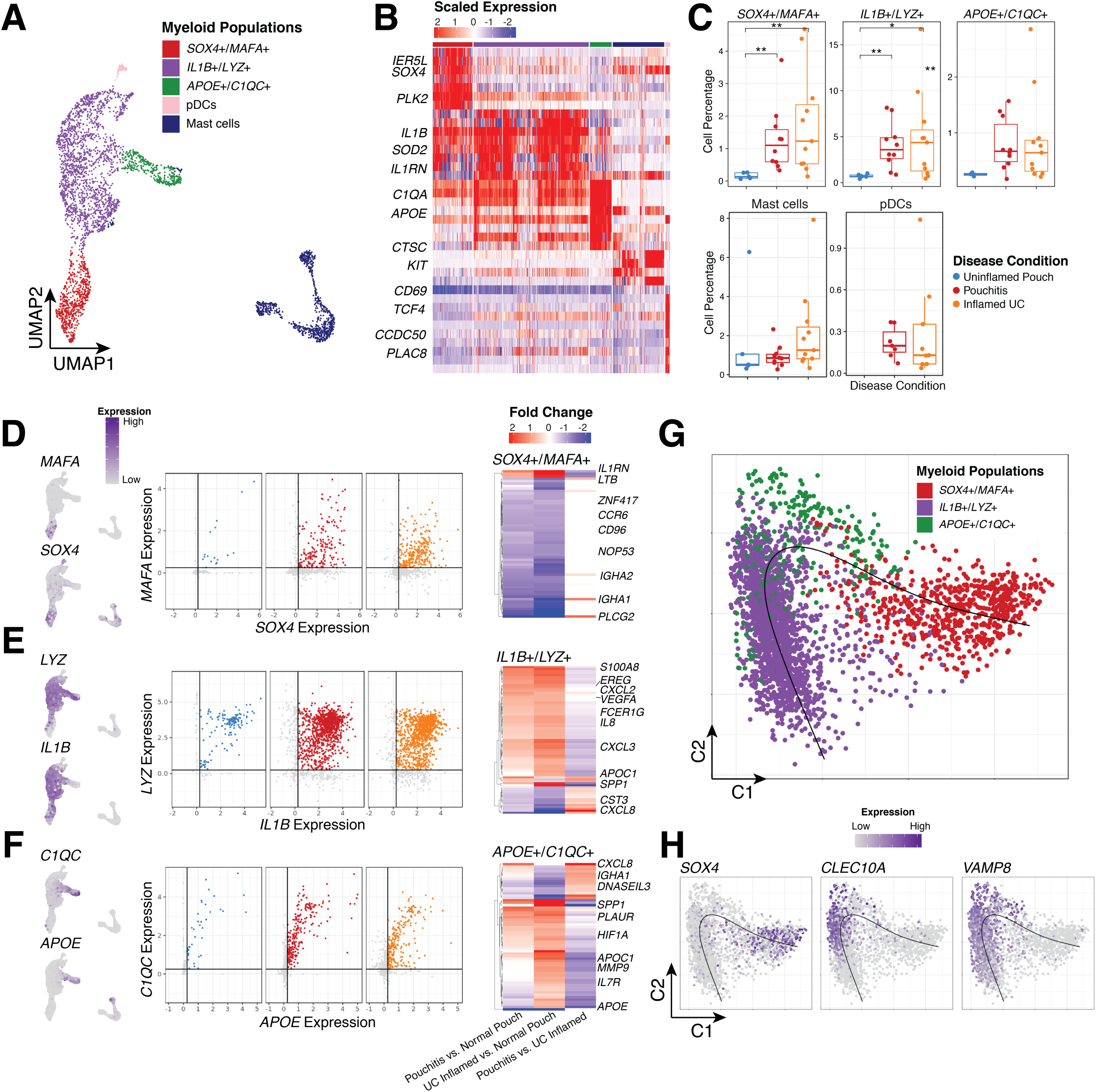
Increased accumulation of myeloid cell populations in inflamed pouchitis and ulcerative colitis patients. (A) Census of myeloid cell types and visualization by UMAP. (B) Heatmap of scaled expression profiles of myeloid clusters, with selected genes of interests shown on left. (C) Boxplots showing cell cluster frequency of the major populations as a percentage of total cells of each patient shown as individual datapoints, compared between patient groups. (D-F) Feature plots showing representative UMAPs of marker gene pairs (left), gene by gene expression plots (center) and differential expression heatmaps of log2 fold change between Uninflamed Pouch, Pouchitis and UC inflamed samples (right) in *SOX4*+/*MAFA*+ (D), *IL1B*+/*LYZ*+ (E) and *APOE*+/*C1QC*+ Monocyte/Macrophage populations. Significantly expressed genes are determined by Log2 fold change greater than 0.75 and adjusted p-value less than 0.05. (G) Diffusion map of *IL1B+/LYZ+, SOX4+/MAFA+* and *APOE+/C1QC+* monocyte/macrophages (top) with a psuedotime projection (black line). (H) *SOX4, CLEC10A* and *VAMP8* normalized expression shown on diffusion map as feature plots (bottom). Asterisks indicate significance testing for Wilcoxon ranked test, * = *p* < 0.05, ** = *p* < 0.01, *** = *p* < 0.001.

Gene expression profiles marking each of the monocyte/macrophage cell clusters indicated enrichment for unique gene pairs that varied within each cluster. In *SOX4*+/*MAFA*+ monocyte/macrophages, *SOX4* and *MAFA* were used as specific markers and used for *in silico* gating to identify cells with enriched expression for both of these genes (Fig. 2D-F). When examined in individual samples, the relative percentage of *SOX4*+/*MAFA*+ cells are significantly increased (*p* < 0.01) in inflamed pouch and UC samples compared to uninflamed pouches. Differential expression between pouch and UC patients in these cells indicated an increase in *IL1RN* and *LTB* in inflamed samples compared to uninflamed pouches (Fig. 2D). Increased relative percentage in inflamed samples was also true for *LYZ+*/*IL1B+* cells in IL1B+/LYZ+ monocyte/macrophages and differentially expressed genes between pouch and UC included significant increases in *EREG, S100A8, CXCL2, VEGFA, IL8* and *APOC1* in inflamed samples (Fig. 2E). Additionally, *SPP1* an important macrophage marker for colorectal cancer^17^, was found to be differentially expressed in inflamed UC samples compared to uninflamed pouches. It was not, however, significant between inflamed and uninflamed pouches. *APOE+*/*C1QC+* cells (Fig. 2F) were also significantly increased in inflamed samples (*p* < 0.01) and *HIF1A, PLAUR, APOC1, MMP9* and *IL7* all increased in inflamed samples. Interestingly, *CXCL8* and *DNASEIL3* increased specifically in pouchitis compared to UC inflamed samples. To further delineate transcriptional relationships between the monocyte/macrophage clusters we performed a pseudotime analysis restricted to the three monocyte/macrophage cell clusters (Fig. 2G). In contrast to the UMAP representation (Fig. 2A) a diffusion map projection with pseudotime maintains the difference between *SOX4*+/*MAFA*+ and *IL1B*+/*LYZ*+ monocyte/macrophages but suggests *APOE*+/*C1QC*+ cells as an intermediary population (Fig. 2E, black line). Specifically, *SOX4* is strongly associated with pseudotime as a marker of *SOX4*+/*MAFA*+ cells (Fig. 2H). *CLEC10A* and *VAMP8* are also associated with pseudotime but were not significantly associated with specific macrophage populations (Fig. 2H).

The *IL1B+/LYZ+* monocyte/macrophages appear similar to butyrate-induced anti- microbial macrophages^18^, which also has a signature characterized by the expression of calprotectin *S100A8/A9*. The expression of *APOE* and *C1QC* indicate those monocyte/macrophages could have a more phagocytic/efferocytotic phenotype^19,20^. In addition to *SOX4* and *MAFA, SOX4*+/*MAFA*+ monocyte/macrophages also express TREM1 and CXCL10 (IP-10) (Supplementary Figure 3) but otherwise express transcripts that are not associated with known macrophage functions. Hence, we identify here 3 types of related monocyte/macrophage cell states that are all increased in the inflamed samples regardless of large or small intestine origin.

### Specific T cell subsets differ according to patient inflammation status and disease

Considering T cells comprised the largest cluster of cells in 25 of the 26 samples profiled we performed further clustering and differential expression analysis to characterize T cell subsets (Supplementary Figure 4A). Specifically, 12 clusters of T cells were identified and characterized by expression of *IFNG, CD8, FOXP3, IL2, ATF3, IL7, TNF, CCL5, GZMB, TOX2* (Fig. 3A and Supplementary Figure 4A-B). From these subsets we find a cluster of *FOXP3*+ T cells (Tregs) increased in inflamed UC and pouch samples compared to uninflamed pouches (Fig. 3B, Supplementary Figure 4C). Naïve and activated CD8+ cells were increased in relative percentage in pouch samples compared to inflamed UC samples while *CD8*+ *NR4A2*+ cells were generally higher in uninflamed pouches compared to inflamed pouch and UC samples (Fig. 3B). For the other T cell subsets that were identified the relative percentages were not significantly different between the patient groups (Supplementary Figure 4D).

**Figure 3.**
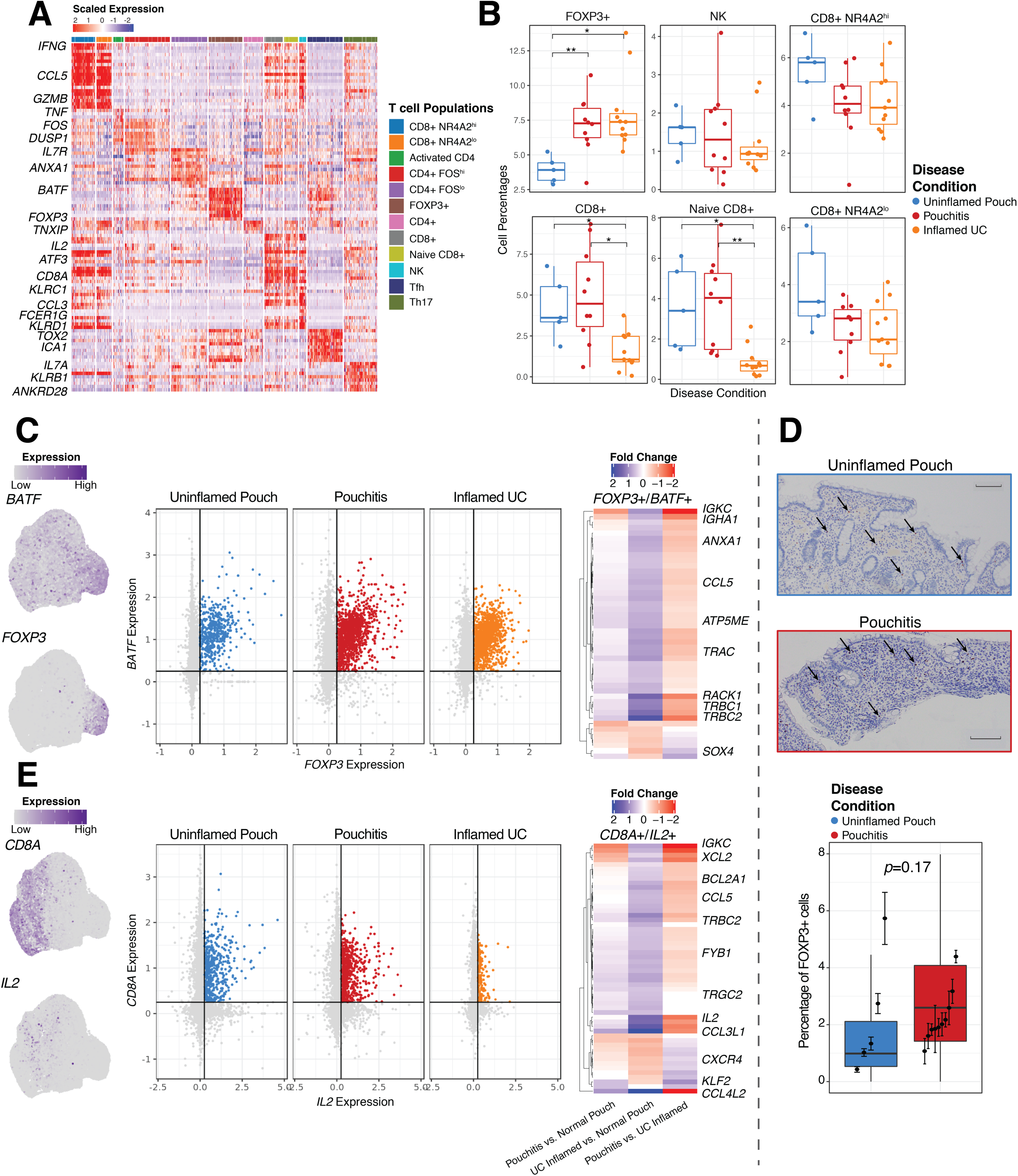
Dysregulation of the T cell compartment in inflamed pouchitis and ulcerative colitis patients. (A) Heatmap of scaled expression profiles of the 12 major T cell clusters, with selected genes of interests shown on left. (B) Boxplots showing cell cluster frequency of selected T cell populations as a percentage of total cells of each patient shown as individual datapoints, compared between patient groups. (C) Feature plots showing representative UMAP visualizations of *FOXP3* and *BATF* expression in T cells (left), *FOXP3* by BATF expression plots (center) and differential expression heatmaps of genes with log2 fold change between Uninflamed Pouch, Pouchitis and UC inflamed samples (right) in *FOXP3*+/*BATF*+ T cells. (D) Representative *FOXP3* stained pouch tissue sections (top) and quantification (bottom), where dark brown colored cells indicate *FOXP3* nuclear expression and quantification of percent *FOXP3*+ cells in 5 uninflamed patient samples and 10 pouchitis patient sample. Significance was determined by nested ANOVA correcting for multiple data points per patient. (E) Feature plots showing representative UMAP visualizations of *CD8A* and *IL2* expression in T cells (left), *CD8A* by IL2 expression plots (center) and differential expression heatmaps of log2 fold change between Uninflamed Pouch, Pouchitis and UC inflamed samples (right) in *CD8A*+/*IL2*+ T cells. Selected genes of interests shown on right. Significantly differentially expressed genes are determined by Log2 fold change greater than 0.5 and adjusted p-value less than 0.05. Asterisks indicate significance testing for Wilcoxon ranked test, * = *p* < 0.05, ** = *p* < 0.01, *** = *p* < 0.001. Scale bars indicate 200μm imaged at 10X magnification.

The marked expansion of *FOXP3*+ T cells in inflamed samples was consistent with a previous report whereby *FOXP3*+ regulatory T cells also expressed TNF^3^. Here, we find that expression of *BATF* is also an important feature of these cells (Fig. 3C) and *in silico* gating of *FOXP3*+/*BATF*+ expression in T cells clearly shows an increase in inflamed pouch and UC samples compared to uninflamed pouches (Fig. 3C). Differential expression analysis of the *FOXP3*+/*BATF*+ cells indicated a number of transcriptional changes between uninflamed pouches, pouchitis and inflamed UC including an increase in *TRBC1, TRBC2, RACK1, ANXA1* and *CCL5* in pouchitis samples compared to UC. Given the importance of *FOXP3* expression in these cells we confirmed their presence by staining for FOXp53 nuclear expression (see methods) in tissue sections from uninflamed pouch (*n*=5) and pouchitis (*n*=10) patients (Fig. 3D). The relative percentage of *FOXP3*+ cells in pouchitis patients again was greater than in the uninflamed pouches (*p*=0.17). In contrast, both *CD8*+ and naïve *CD8*+ T cells were less represented in the colon samples from inflamed UC patients compared to the pouch samples, regardless of inflammation status (Fig. 3B). Within these CD8+ populations *in silico* gating of *CD8*+/*IL2*+ expression in T cells identified differences between inflamed UC and pouch patients regardless of inflammation (Fig. 3E). *IL2* expression was identified as a marker for *CD8*+ and naive *CD8*+ T cell subsets (Fig. 3A). Additionally, differential expression analysis of the *CD8*+/*IL2*+ cells indicated *IL2, CCL4L2, CCL5, TRBC2, XCL2* and *IGKC* are all increased in both pouchitis and uninflamed pouch compared to inflamed UC (Fig. 3E). One possibility is that this difference between pouch and colon may reflect the increased proportion of CD8+ T cells observed in the ileum compared with colon^21,22^. Hence, we identified here *FOXP3*+*BATF*+ T cells as being the most increased population in inflamed samples, whereas CD8+ cells in general are more abundant in the pouch than in colon samples regardless of inflammation state.

### B cell subsets are similar across disease types

Recently, intestinal inflammation in UC was reported to be associated with increased anti-commensal IgG, which could induce IL-1B by engaging Fcγ receptors on macrophages^23^ indicating a role for B cells in pathogenesis. Despite their high frequency in many of the patient samples profiled, B cell subsets clustered into only 5 clusters across 26 patient samples (Supplementary Figure 5A). These subsets include cycling B cells, follicular cells, germinal center cells and two types of plasma cells which differ in *NFKBIA* expression (Supplementary Figure 5A). Traditional B cell markers and differentially expressed transcripts were used to identify these subtypes including *STMN1, MKI67, BANK1, LMO2, LY9, MZB1 and XPB1* (Supplementary Figure 5B). While there is a slight increase in plasma cell percentages in inflamed UC samples compared to inflamed and uninflamed pouches, there were no other significant differences in B cell states between the patient groups (Supplementary Figure 5C).

### Validation of inflammation associated immune cell populations in public datasets of ulcerative colitis and Crohn’s disease patient samples

*FOXP3*+ Tregs may be induced or maintained by intestinal antigen presenting cells such as macrophages and dendritic cells^24^. To investigate this relationship further, we compared the relative percentage of *FOPX3+/BATF+* T cells and each of the specific monocyte/macrophages subsets as defined by their respective markers *SOX4, MAFA, IL1B, LYZ* and *APOE, C1QC* by linear regression (Fig. 4A). While, all three populations were significantly associated, the *SOX4*+/*MAFA*+ cells were most strongly associated (R^2^=0.388). *TREM1* is expressed by *SOX4*+/*MAFA*+ monocyte/macrophage cells and can be induced by retinoic acid^25^ which also induces *FOXP3*+ Treg differentiation. However, apart from *CXCL10* expression, these cells do not exhibit many other features of immune activation (Supplementary Figure 3A).

**Figure 4.**
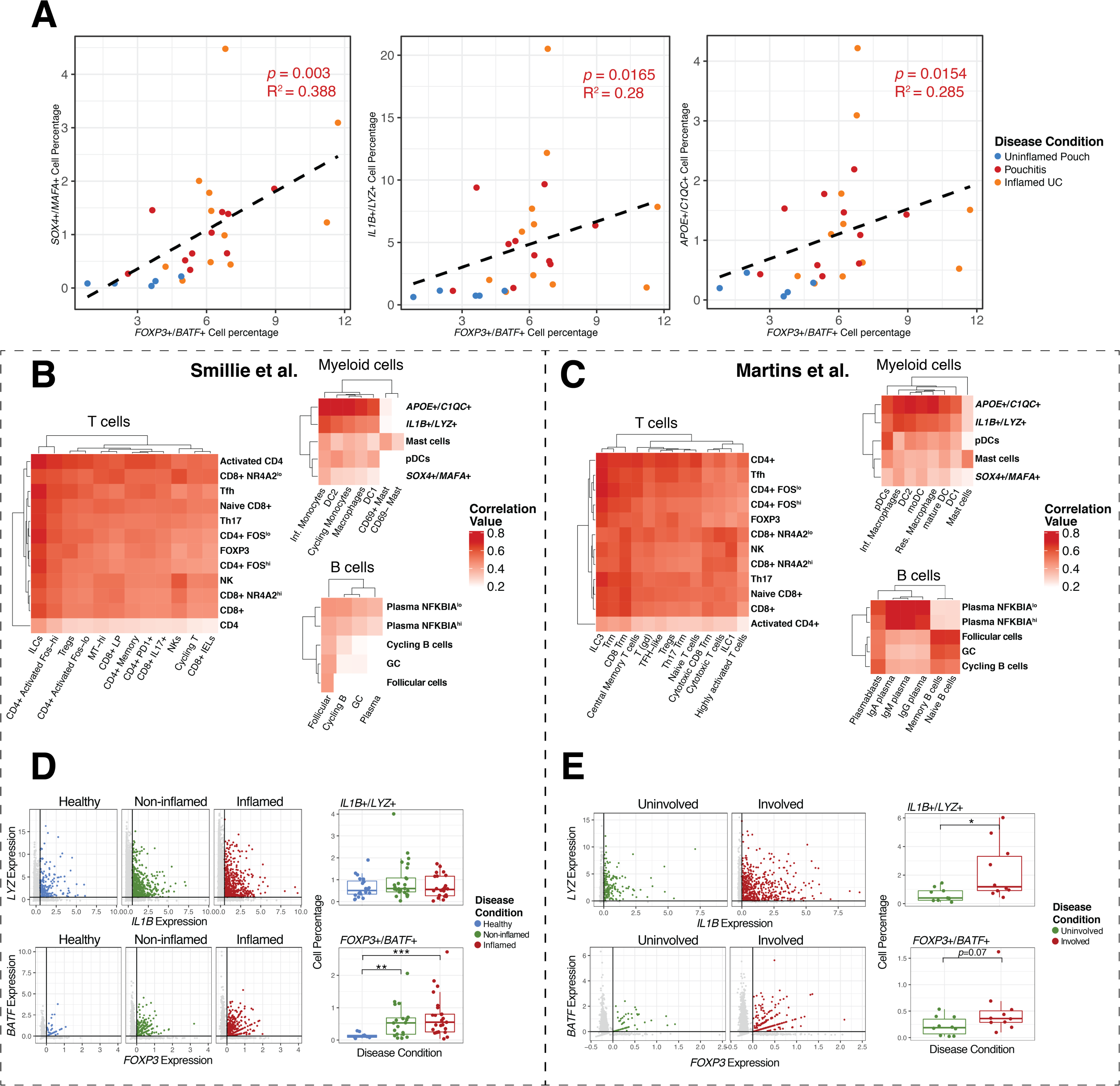
Defining cell states related in active inflammation in J-pouch, ulcerative colitis and Crohn’s disease. (A) Linear model of the associations between the percentage of *FOXP3*+*BATF*+ cells and *SOX4*+*MAFA*+ (left), *IL1B*+*LYZ*+ (middle) and *APOE*+*C1QC*+ cells (right). R^2^ and *p* value was determined by linear regression. Individual samples are shown and color coded based on disease condition. (B, C) Heatmaps of correlated cell types from scRNA-seq datasets of Smillie et al. (B) and Martins et al. (C), comparing T cell (left), myeloid subsets (top) and B cell subsets (bottom). (D) Single cell expression of *IL1B* and *LYZ* (top) and relative percentage of LYZ+IL1B+ cell populations from healthy, inflamed and non-inflamed ulcerative colitis patient samples from Smillie et al. Single cell expression of *FOXP3* and *BATF* (bottom) and relative percentage of *FOXP3*+*BATF*+ cell populations from healthy, inflamed and non-inflamed ulcerative colitis patients samples from Smillie et al. (E) Single cell expression of *IL1B* and *LYZ* (top) and relative percentage of LYZ+IL1B+ cell populations from matched involved and uninvolved Crohns disease patient samples from Martins et al. Single cell expression of *FOXP3* and *BATF* (bottom) and relative percentage of *FOXP3*+*BATF*+ cell populations from involved and uninvolved Crohns disease patient samples from Martins et al. Asterisks indicate significance testing for Wilcoxon ranked test, * = *p* < 0.05, ** = *p* < 0.01, *** = *p* < 0.001.

Recently, studies from Smillie^3^ et al. and Martin^26^ et al. have provided additional datasets for single cell analysis of IBD subsets in UC and Crohn’s Disease. We compared the 22 immune cell populations identified in this study with those public datasets. Based on the expression profiles of T cell subsets, many T cell populations were highly correlated with at least one other labeled T cell subset from Martin^26^ or Smillie^3^ et al., r > 0.6, with the exception of activated CD4 cells which did not correlate in Martins et al., and CD4+ cells in Smillie et al. (Fig. 4B,C). *IL1B*+/*LYZ*+, *APOE*+/*C1QC*+ monocyte/macrophages, mast cells and pDCs were also highly correlated with named populations “Inflammatory Monocytes” and “Resident Macrophages” from both public datasets (Fig. 4B, C). However, *SOX4*+/*MAFA*+ monocyte/macrophages were not strongly correlated in either of the two datasets, r < 0.4. It is possible that CD45+ enrichment of immune cells enabled us to identify this monocyte/macrophage population that was not previously detected. In B cells many of the subsets were correlated with named populations from both Martin^26^ and Smillie^3^ et al. including plasma cells, germinal centers, cycling B cells and follicular cells, r > 0.6 (Fig. 4B, C).

After confirming the identify of *FOXP3*+ regulatory T cells and *IL1B*+/*LYZ*+ monocyte/macrophages populations in public datasets, we wanted to validate the relationship of these cell subsets in inflammation. In both CD and UC patient samples, we found that *FOXP3*+/*BATF*+ regulatory T cells were significantly increased in inflamed/involved compared to non-inflamed and healthy patient samples respectively (Fig. 4D,E). We did not find major differences in the percentage of *IL1B*+/*LYZ*+ monocyte/macrophage cells in the data from Smillie^3^ et al., which profiled inflamed UC and paired non-inflamed UC samples (*n*=18) and healthy samples (*n*=12) (Fig. 4D). However, CD patients from Martin^26^ et al. with active inflammation (*n*=11) did exhibit significant increases in *IL1B*+/*LYZ*+ cell subsets compared to matched uninvolved tissue (Fig. 4E). No major differences between inflamed and uninflamed samples were found for *APOE*+/*C1QC*+ monocyte/macrophages (Supplementary Figure 6). We could not detect *MAFA* in either dataset so we used *MAFB* which was also a marker. However, even *SOX4*+/*MAFB*+ cell populations were scarce in either datasets and did not allow us to determine if these cells were associated with inflammation (Supplementary Figure 6). Hence, we find that increased abundance of *FOXP3*+/*BATF*+ T cells is a consistent feature of inflammation in both UC and CD patients, whereas enrichment of specific monocyte/macrophage subsets could be more context-dependent.

### Prioritizing inflammatory markers in a large cohort of IPAA patients

We extracted 453 non-overlapping signature genes by differential expression and outlier analysis^27^ (Fig. 5A, Supplementary Table 1) as a signature matrix for the 12 T cell, 5 B cell and 5 myeloid cell subsets identified by scRNA-seq. We also extracted gene expression data for genome wide association study (GWAS) risk genes that are associated with IBD^3^ from our scRNA-seq signature (Fig. 5A, Supplementary Figure 7). 17 of the GWAS-implicated IBD risk alleles were associated with specific immune cells populations (Fig. 5A). For example, *IFNG* was associated with CD8+ T cells, *CCL20* was associated with Th17 cells and *LY9* was associated with follicular B cells. 45 other IBD-associated genes were also found in the scRNA-seq dataset but were not found to be significant markers of our immune cell signatures (Supplementary Figure 7) indicating that expression of these genes was not specific to a particular immune cell population.

**Figure 5.**
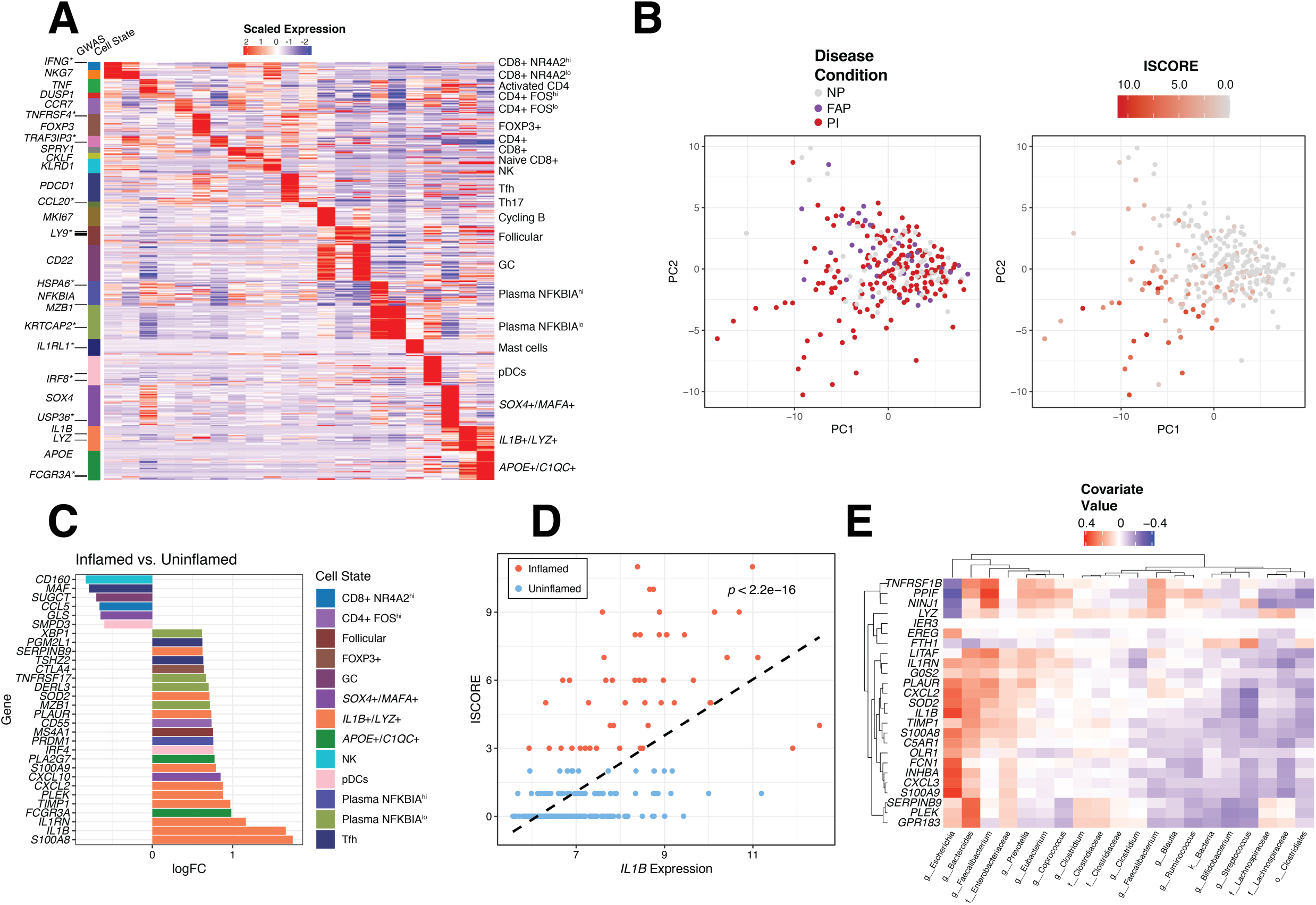
Identification of immune cell type specific transcripts and analysis of an independent dataset of patients with IPAA. (A) Heatmap illustration of a signature matrix of 453 marker genes for 22 identifiable T, B and Myeloid cell states from scRNA-seq. Asterisks indicate GWAS genes related to inflammatory bowel disease and the expression of these genes in the specific cell types on the right. (B) Principal component analysis of 250 IPAA patient samples with familial adenomatous polyposis (FAP), active pouchitis (PI) or no pouchitis (NP) based on the 453-gene signature and colored by Disease Condition (left) and inflammation score (ISCORE) (right). (C) Expression of *IL-1B+/LYZ+* monocyte/macrophage cell type specific transcripts from biopsy samples collected from inflamed (ISCORE > 2) versus uninflamed (ISCORE < 2) patient samples. Shown are log fold change values of the most differentially expressed genes from the 453 cell type specific markers colored by the corresponding cell type from scRNA-seq. (D) Linear model of the association between IL1B expression and the numeric inflammation score for 250 patient samples of IPAA. (E) sPLS analysis of a 25-gene signature for *IL- 1B+/LYZ+* monocyte/macrophages compared to microbial taxa measured by 16S rRNA sequencing of the 250 patient samples of IPAA. Directionality of the association between gene expression and bacterial abundance is colored in red or blue as shown in the legend.

After generating an immune cell signature matrix, we examined the expression of these genes in a microarray dataset from a larger study of IPAA patients (n=250) with active pouchitis (PI), uninflamed pouches (NP) and familial adenomatous polyposis (FAP)^28^. This dataset also included 16S rRNA microbial profiling data from paired biopsies. Based on the 453-gene signature matrix generated by our single cell data, there were no appreciable differences between groups of IPAA patient samples by PCA (Fig. 5B). However, we found greater separation of patient samples in relation to the composite inflammation score (ISCORE)^28^. Patients with the highest ISCORE contributed most to the separation in PCA space (Fig. 5B). We therefore split the patient samples according to the ISCORE values into a low (ISCORE < 2, *n*=223) and high (ISCORE > 2, *n*=50) inflammation group and performed differential expression analysis by limma^29^ version 3.38.3 limited to the 453 gene signature from our single cell dataset. The most differentially expressed transcripts in inflamed patient samples were related to the proinflammatory *IL1B*+/*LYZ*+ monocyte/macrophages, marked by *IL1B, S100A8, IL1RN* and *CXCL2* (Fig. 5C). We also found that expression of the inflammatory marker *IL1B* was significantly related to ISCORE (Fig. 5D). To determine if there are microbial associations with this set of host-transcripts, we employed sPLS regression to identify microbial taxa most associated with the *IL1B*+/*LYZ*+ monocyte/macrophages signature genes. We find that the expression of these genes is positively correlated with the abundance of *Escherichia, Bacteroides, Faecalibacterium* and inversely correlated with abundance for the order *Clostridiales* (Fig. 5E). Hence, there may be particular macrophage-microbe interactions that drive this anti-microbial IL-1B signature in inflammatory macrophages specifically during inflammation of the pouch.

### Evaluating response to clinical therapies and inflammation status in ulcerative colitis

In order to identify cytokine signaling networks and immunoregulatory mechanisms between the 22 immune cell subsets in UC, we next investigated receptor-ligand pair networks between cells^30^ to construct a cellular interaction network (Fig. 6A). Using a curated database of receptor-ligand interactions from cellPhoneDB^31^, we identified pairs of interacting cell subsets based on our 453 gene signature matrix. Across the 22 immune cell subsets we identified 629 significant interactions between receptor-ligand pairs. *IL1B*+/*LYZ*+ and *APOE*+/*C1QC*+ monocyte/macrophages had the most interacting pairs and are the largest nodes of this network by number of connections, with many significant interactions between the two cell types. Additionally, Th17 cells are an important interacting node with these macrophage populations (Fig. 6A). The interactions between Th17 cells and *IL1B*+/*LYZ*+ monocyte/macrophages and *APOE*+/*C1QC*+ monocyte/macrophages include activity involving the chemokines *CCL3, CCL4, CCL5* and *CCL20* with the receptors *CCR1, CCR4, CCR5, CCR6* (Fig. 6B). Costimulatory molecule interactions with their ligands such as *CTLA-4, ICOS, PD-1* and *CD40* were also significant between Th17 and *FOXP3*+ T cells with the *IL1B*+/*LYZ*+ and *APOE*+/*C1QC*+ monocyte/macrophages populations (Fig. 6B). Some of the other notable interactions relate to cytokines such as *TNF* and *CSF*, which are important in disease pathogenesis^32^. In summary, this approach enabled us to determine that *IL1B*+/*LYZ*+ monocyte/macrophages interactions with Th17 cells could be an important component of the intestinal immune response during inflammation for these UC patients.

**Figure 6.**
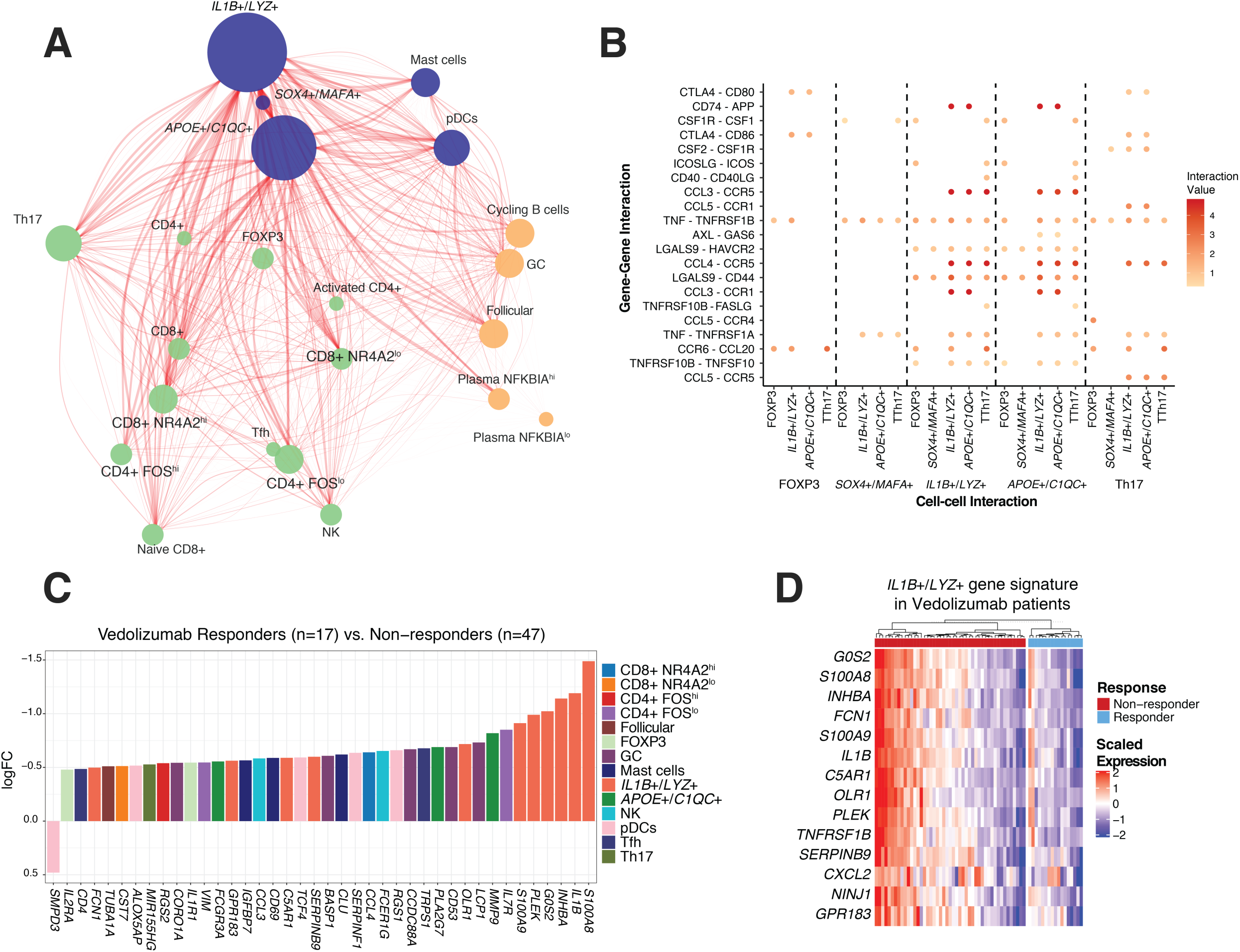
Evaluating response to clinical therapies and inflammation status in ulcerative colitis. (A) Analysis of receptor-ligand network connections between 22 of the major cell populations identified in scRNA-seq. Each node represents a cell population and each edge a significant receptor-ligand association according to curated database, cellPhoneDB. The size of each node is proportional to the number of connected edges and the thickness of the edges is proportional to the significance value of the connection. (B) Visualization of selected ligand–receptor interactions that are specifically enriched between FOXP3+ Tregs, *SOX4+/MAFA+, IL1B+/LYZ+, APOE+/C1QC+* monocyte/macrophages and Th17 cells. Interaction values are indicated by intensity, scale on right. (C) Expression of *IL-1B+/LYZ+* monocyte/macrophage cell type specific transcripts from biopsy samples collected from ulcerative colitis patient Responders versus Non-responders to treatment with the anti-α4β7 integrin antibody Vedolizumab. Log fold change values of the most differentially expressed genes from a matrix of 453 cell-type specific markers between Responders and Non-responders to treatment are shown. D) Heatmap showing the heterogeneity in the expression of *IL-1B+/LYZ+* monocyte/macrophage cell type specific transcripts among Non-responders to Vedolizumab treatment. Normalized expression values shown for individual patients from Responders (n=17) and Non-responders (n=47).

Several new therapeutic agents developed for treating UC patients are designed to block immune cell interactions. Etrolizumab and Vedolizumab target the α4β7 integrin, and Golimumab targets TNFα, but there is considerable heterogeneity in patient responsiveness that is poorly understood. We next assessed if the cell specific signature of immune cell subsets could distinguish between responders and non-responders for 3 different treatments for UC, Vedolizumab^33^ (GSE73661), Etrolizumab^34^ (GSE72819), and Golimumab^35^ (GSE92415). We examined differential expression between responders and non-responder in for the 453-gene signature matrix for these treatment studies. Transcripts associated with *IL1B*+/*LYZ*+ monocyte/macrophages were the most significantly different between responders and non-responders for Vedolizumab treatment. The non-responders were most enriched in transcripts for the proinflammatory *IL1B*+/*LYZ*+ monocyte/macrophages subset (Fig. 6C,D), indicating that the presence of these macrophages could be an indicator of resistance to α4β7 integrin blockade. In addition to *IL1B*, calprotectin components *S100A8/A9* were more highly expressed in non-responders. However, there is considerable heterogeneity, in that approximately 50% of non-responders have high expression of *IL1B*+/*LYZ*+ monocyte/macrophages transcripts (Fig. 6D). Non-responders to Etrolizumab treatment also exhibited higher expression of proinflammatory genes like *IL1B, IL1RN* and *S100A9* compared to responders (Supplementary Figure 8). Golimumab non-responders, while still slightly enriched for expression of these transcripts, are less strikingly associated with *IL1B*+/*LYZ*+ monocyte/macrophages transcripts (Supplementary Figure 8), perhaps reflecting the different mechanism of action. Hence, we find that a proportion of non-responders to α4β7 integrin blockade in UC patients are associated with increased transcripts for an inflammatory monocyte/macrophage population that expresses an anti-microbial signature.

## DISCUSSION

In this study, we found that inflammation in the ileal-anal J pouch, a novel organ created from ileal tissue has an inflammatory landscape similar to the colon of ulcerative colitis patients. We identified *FOXP3*+/*BATF*+ T cells and 3 different monocyte/macrophage populations (*IL1B+/LYZ+, SOX4+/MAFA+* and *APOE+/C1Q+*) with inflamed tissues in both UC and pouchitis. Of these cells, the *IL1B+/LYZ+* monocyte/macrophages were the most highly connected cell type in the inflammatory network and their signature was associated with the lack of responsiveness to α4β7 integrin blockade and an increased abundance of *Bacteroides* and *Escherichia* bacterial populations. We hypothesize that the increased activation of these *IL1B+/LYZ+* monocyte/macrophages may be representative of a different intestinal inflammation state that is indicative of an immune response that cannot be modulated by α4β7 integrin blockade.

Our initial goal was to compare immune infiltration for the J-pouch with the colon, as two organs performing a similar function but with the J-pouch originating from the small intestine. This analysis concludes that the inflammatory response for pouchitis and UC is remarkably similar despite the different origin tissues. While the pouch has more CD8+ T cells of different phenotypes, which may reflect the increased proportion of CD8+ T cells observed in the ileum compared with colon^21,22^, this is not significantly altered by the inflammatory response. This data supports previous studies indicating that pouchitis and UC are driven by similar inflammatory mechanisms^15^ and may therefore respond to similar therapeutics^36–40^. Notably, we did not characterize the CD45+ immune cells of the pouch for patients with FAP, who develop pouchitis less frequently and may reveal features unique for the UC associated pouch even without the presence of inflammation. Recently, secondary bile acids and associated microbes were found to distinguish UC and FAP pouches, and may result in the pro-inflammatory conditions preceding pouchitis in UC patients^41^. Efforts are underway in determining the distinct roles of secondary bile acids and butyrate, both of which are byproducts of similar bacterial taxa that mediate Th17 and Treg polarization, and discerning the action of these metabolites on antigen presenting cells versus T cells remains an important area of research^11,41–45^. Thus, it is important to identify any association between the *IL1B+/LYZ+* monocyte/macrophages and other metabolites such as bile acids beyond butyrate^18^.

The relationship between microbes and antimicrobial macrophages is particularly relevant to the pathogenesis of IBD. ScRNA-seq of human monocyte derived macrophages treated with butyrate had previously been shown to induce an antimicrobial signature through *HDAC3* including upregulation of autophagy-related processes^18^. Here we provide evidence that macrophages with an overlapping antimicrobial signature can directly be identified from intestinal biopsies of UC patients. This observation may reflect the recruitment of monocytes to the gut where they differentiate into macrophages to counteract a breach in the epithelial barrier, a characteristic of IBD patients who are colonized by pro-inflammatory bacteria related to *Bacteroides* and *Escherichia* species^46–48^. In contrast to healthy individuals, local cytokine responses and inefficient autophagy may prevent macrophages from resolving the breach in the barrier, leading to a detrimental pro-inflammatory effect of macrophages^49^. Consistent with this possibility, our previous work indicates that antimicrobial monocytes and macrophages recruited to the gut are beneficial when damage to the colon is temporary, even with an inflammatory cytokine signature exacerbated by the absence of autophagy^50–52^.

The most consistent finding between our scRNA-seq data and the meta-analysis of data from public datasets is the expansion of *FOXP3*+/*BATF*+ Tregs in actively inflamed IBD. Expression of *BATF* in FOXP3+ Tregs is indicative of tissue residency^53^. In the visceral adipose tissue, FOXP3+ Tregs require *BATF* for differentiation downstream of *ST2* and *PPARG* activity^54^. In the intestine, *BATF* regulates expression of *CCR9* and a4b7 and *BATF* deficient mice have reduced effector as well as FOXP3+ T cells in the intestine^55^. Hence, *BATF* is likely an important transcription factor for the differentiation and recruitment of FOXP3+ Tregs to the intestinal tissues during inflammation. Notably, a recent report on scRNA-seq analysis of immune cell populations in immune checkpoint inhibitor-induced colitis also reveals the persistence and expansion of Tregs^56^. The increased accumulation of FOXP3+ Tregs in the inflamed colon is likely driven by the need to restrain inflammation, but why these Tregs are not successful in controlling inflammation requires further study. One possibility may be that the presence of inflammatory monocyte/macrophage populations can somehow inhibit the appropriate function of these regulatory cells.

We also observe expansion of *APOE*+/*C1QC*+ monocyte/macrophages with a phagocytic signature in both the inflamed pouch and colon, which are reminiscent of the tumor associated macrophage (TAM) populations recently described in colorectal cancer (CRC) patients^57^. In that study, the C1QC+ macrophages were found to be closely connected with IL1B+ macrophages, which we also find in this study. This indicates that there are some similar characteristics between the myeloid infiltration observed in CRC and UC. However, the role of these macrophages beyond having a phagocytic signature remain unclear and will require further study. Notably, we did not observe the *SPP1*+ macrophage described in CRC patients^17,57^.

We had previously found increase in Th17 cells in inflamed biopsies^58,59^ linked by expression of *SAA1* with *Bacteroides* abundance, but this was by flow cytometry and intracellular cytokine staining. Serum amyloid A (SAA) proteins produced by intestinal epithelial cells can drive differentiation of inflammatory Th17 cells according to the tissue environment^60^, however expression of SAA proteins was not detected in the present study because we selected for immune cells. We also do not observe significantly more Th17 cells by transcriptional signature (Supplementary Figure 4D). Nonetheless, this population of cells is highly connected with the *IL1B*+/*LYZ*+ and *APOE*+/*C1QC*+ populations that are expanded in inflamed samples and hence their activity could be highly dependent on the interaction with these macrophages. Causal relationships between these populations may be discerned in the future through *in vitro* co-culture assays but purifying specific populations of these immune cells will entail surgical specimens rather than the mucosal pinch biopsies utilized in this study.

IL-1B release can be triggered by activating *NLRP3* and other inflammasomes in macrophages exposed to invasive microbes. Elegant experiments examining very early onset IBD (VEOIBD) patients and mouse models genetically deficient in IL-10 signaling indicate that inflammasome-triggered IL-1B production by macrophages polarizes CD4+ T cells that mediate colitis^61,62^. Blocking IL-1B signaling was effective in two IL10R-deficient patients with treatment-refractory disease^62^, suggesting that macrophage-T cell interactions (Fig. 6A-B) drive disease in the absence of IL-10 and other immuno-suppressive Treg effectors. This would be consistent with predictions from the receptor ligand analysis performed in this study (Fig. 6). If the presence of *IL1B*+ macrophages are indicative of unresponsiveness to α4β7 integrin blockade as suggested by our results, therapies that target *IL1B* itself or JAK/STAT inhibitors that broadly target signaling downstream of IL-1B-induced cytokines^63^ may be promising alternatives for UC patients with this signature. Detailed analysis of tissue from patients receiving JAK/STAT inhibitors will be highly informative.

In conclusion, this work utilizes scRNA-seq to identify unique features of pouchitis and a specific population of *IL1B*+ macrophages that could potentially be targeted in a subset of UC patients who are not responding to treatment with α4β7 integrin antagonists. Hence, this study provides an example of how utilizing precision medicine to identify changes in cell proportions, gene expression and cell-cell signaling by scRNA-seq, followed by further analyses of publicly available datasets, could be used to improve our understanding of individual patient responsiveness to IBD therapies and provide a hypothesis for alternative treatment options.

## METHODS

### Study Participants

#### Participants were recruited and consented and are part of an institutional

review board–approved study (S12-01137; “Mucosal immune profiling in patients with inflammatory bowel disease”) by NYU Langone Health. We identified 15 patients with a J-pouch and 13 patients with UC (Supplementary Table 2). All J-pouch patients had preoperative UC with 14 (93%) and 1 (7%) undergoing an IPAA for medically refractory disease and colitis-associated neoplasia, respectively. Of J-pouch patients, 10 (67%) had endoscopic evidence of inflammation (endoscopic PDAI ≥2) referred to as pouchitis and 5 (33%) had endoscopically normal appearing pouches referred to as uninflamed pouches (Supplementary Table 2). All UC patients had moderate to severe endoscopic activity (endoscopic Mayo ≥2). Patients with UC or a J-pouch at the Inflammatory Bowel Disease Center at NYU Langone Health, New York, were approached for recruitment on presentation for routine endoscopy (pouchoscopy, colonoscopy, or flexible sigmoidoscopy) performed for disease activity assessment. Potential participants were excluded if they were unable to or did not consent to provide tissue. The endoscopic appearance determined the inflammatory activity. UC patients were limited to those with active inflammation denoted by an Mayo endoscopic subscore of ≥2^64^. J-pouch patients were stratified into an endoscopic pouchitis cohort for those with an pouchitis disease activity index (PDAI) endoscopic subscore ≥2 or a normal J-pouch cohort for those with an PDAI endoscopic subscore <2^65^. Further patient details and stratification are described in Supplementary Table 2.

### Biopsies

Approximately four to ten mucosal pinch biopsies were obtained from each patient. For UC patients, all biopsies were obtained from the rectum. For J-pouch patients, all biopsies were obtained from the pouch body or inlet. If active endoscopic inflammation was present, this area was targeted for biopsy. For each location sampled, one biopsy was collected for standard histopathology assessment and read by two expert pathologists for the PDAI histology subscore and histologic pouch activity score (PAS)^65,66^.

### Patient metadata

At endoscopy, clinical data was collected including demographics, initial IBD subtype and phenotype, smoking status, the presence of primary sclerosing cholangitis (PSC), extra-intestinal manifestations, and comorbidities, age and indication for IPAA, previous and current medication exposures, and clinical indices of disease activity including the partial Mayo score^64^ for patients with UC and PDAI clinical subscore for patients with a J-pouch^65^ (Supplementary Table 2).

### Biopsy specimen processing

All biopsies were collected in ice cold complete RPMI 1640 (10% FBS, 100x penicillin/streptomycin/glutamine, 50uM 2-mercaptoethanol, Sigma) during endoscopy and subsequently cryopreserved in freezing media (90% FBS + 10% DMSO) for long-term storage. Cryopreserved biopsies were gently thawed at 37°C and enzymatically digested in collagenase VIII (Sigma) and DNase (Sigma) for 1h to obtain a single cell suspension. After live/dead cell staining with near-IR stain (Invitrogen), cell surface markers were labeled with the following antibodies: CD45 PE-Cy7, CD3 PerCP-Cy5.5, CD19 PE, CD14 FITC, and CD16 Pacific Blue (BioLegend). Sorted CD45+ cellular suspensions were isolated using the Sony SY3200 cell sorter and prepared for single-cell RNA sequencing.

### Single cell library and sequencing

From pinch biopsies all samples were sorted and CD45+ cellular suspensions were loaded on a 10x Genomics Chromium instrument to generate single-cell gel beads in emulsion (GEMs). Approximately 10,000 cells were loaded per channel. Single-cell RNA-Seq libraries were prepared using the following Single Cell 3’ Reagent Kits v2: Chromium™ Single Cell 3’ Library & Gel Bead Kit v2, Single Cell 3’ Chip Kit v2, and i7 Multiplex Kit (catalog# PN-120237, PN-120236, # PN-120262, 10x Genomics)^67^ and following the Single Cell 3’ Reagent Kits v2 User Guide (Manual Part # CG00052), Rev A. Libraries were run on an Illumina HiSeq 4000 as 2 × 150 paired-end reads, one full lane per sample, for approximately >90% sequencing saturation.

### Single cell analysis pipeline

The Cellranger software suite (https://support.10xgenomics.com/single-cell-gene-expression/software/pipelines/latest/what-is-cell-ranger) from 10X was used to demultiplex cellular barcodes, align reads to the human genome (GRCh38 ensemble, http://useast.ensembl.org/Homo_sapiens/Info/Index) and perform UMI counting. From filtered counts Seurat^16^ version 3.1.3 was used to process the single cell data including dimension reduction, UMAP representation and differential expression to identify cell type specific markers and differentially expressed genes between pouch and UC conditions by a Wilcox test. All single cell processing steps, code and gene expression tables are described in detail on our github at https://github.com/ruggleslab/Pouch. We also used additional single cell analysis software including diffusion map and pseudotime analysis from the R package Slingshot^68^ version 1.0.0.

### Immunohistochemistry staining for FOXP3+ cells

Image panels were acquired with ×10 lens using an Olympus BX53 microscope equipped with an Olympus DP27 digital color camera (Olympus, Center Valley, PA, USA). FOXp53 nuclear expression, stained with 3,3-Diaminobenzidine (DAB) was digitally quantified by QuPath^69^ using hematoxylin as a background stain. Areas of interest were drawn with the line or polygon drawing tool. *FOXP3* cells and background immune cells were annotated. The images were thresholded using a binary categorization of positive (DAB, brown stain) and negative (blue stain). Default software settings were used for the final analysis with the help of cell analysis, positive cell detection command as previously published^70^. The measurement table was then exported to excel spread sheet for statistical analysis.

### Generating Receptor-ligand networks

Receptor-ligand networks were generated using the software CellPhoneDB^31^ using the default databases and methods as described in their documentation (https://github.com/Teichlab/cellphonedb).

## DATA AND CODE AVAILABILITY

Raw sequence data are deposited in the NCBI Sequence Read Archive under BioProject accession number XXX and gene expression omnibus (GEO) accession number GSEXXX. All processing was performed in R^71^ version 3.5.1 and complete analysis scripts can be found on github at https://github.com/ruggleslab/Pouch.

## Acknowledgements

We wish to thank the NYU School of Medicine Flow Cytometry and Cell Sorting, Microscopy, Genome Technology, and Histology Cores for use of their instruments and technical assistance (supported in part by National Institute of Health (NIH) grant P31CA016087, S10OD01058, and S10OD018338).

## Author contributions

Design of experiments, data analysis, data discussion, and interpretation: J.C.D., J.A., S.C., K.V.R., D.H., K.C. and P.L.; primary responsibility for execution of experiments: A.M.H., J.D.L., Single cell analysis: J.C.D., K.V.R. All authors discussed data and commented on the manuscript.

