## Supplemental Figures for "Transcriptional Atlas of Ileal-Anal Pouch Immune Cells from Ulcerative Colitis Patients"

**Supplementary Tables**

**Supplementary Table 1.** Signature matrix of 22 identified immune cell subsets from scRNA-seq of pouch and ulcerative colitis patient samples

**Supplementary Table 2.** Clinical metadata of pouch and ulcerative colitis patients included in our study

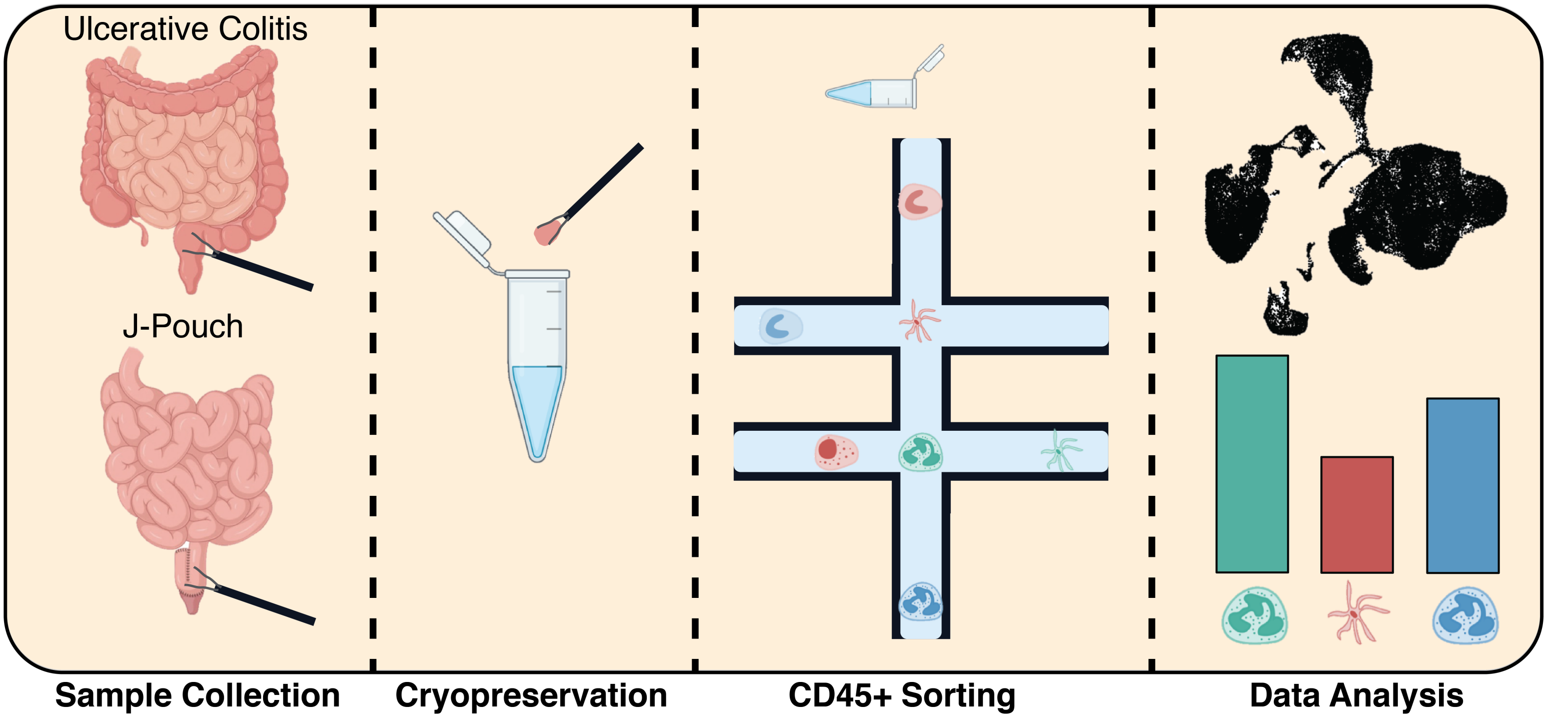
Ulcerative Colitis
J-Pouch
Sample Collection
Cryopreservation
CD45+ Sorting
Data Analysis

**Supplementary Figure 1.** Workflow for collection and processing of pinch biopsy samples of ulcerative colitis patients with active disease or with a J-pouch for scRNA-seq.

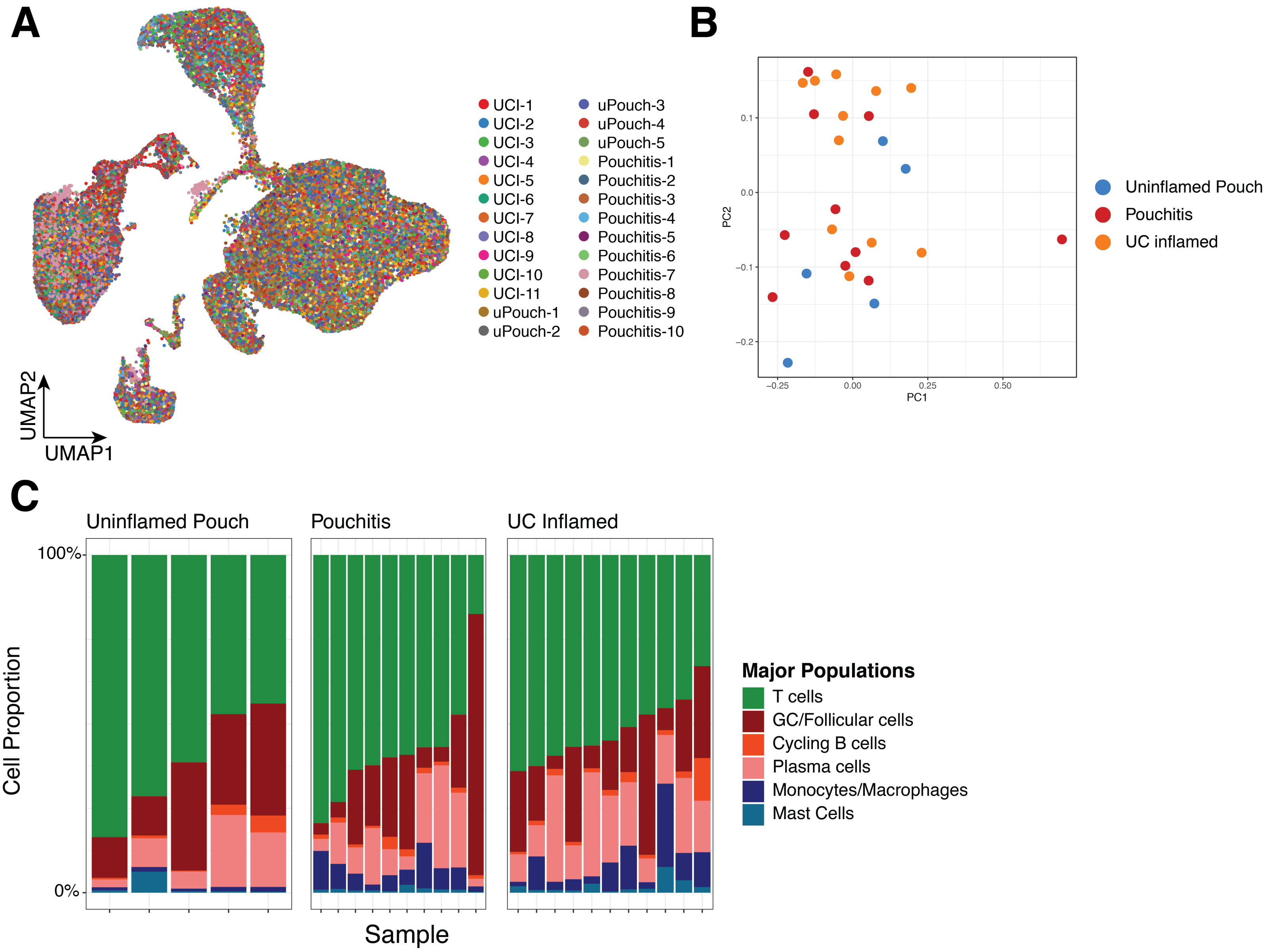
A
UCI-1
UCI-2
UCI-3
UCI-4
UCI-5
UCI-6
UCI-7
UCI-8
UCI-9
UCI-10
UCI-11
uPouch-1
uPouch-2
uPouch-3
uPouch-4
uPouch-5
Pouchitis-1
Pouchitis-2
Pouchitis-3
Pouchitis-4
Pouchitis-5
Pouchitis-6
Pouchitis-7
Pouchitis-8
Pouchitis-9
Pouchitis-10
UMAP2
UMAP1
B
PC2
PC1
Uninflamed Pouch
Pouchitis
UC inflamed
C
Uninflamed Pouch
Pouchitis
UC Inflamed
100%
0%
Cell Proportion
Sample
Major Populations
T cells
GC/Follicular cells
Cycling B cells
Plasma cells
Monocytes/Macrophages
Mast Cells

**Supplementary Figure 2.** (A) UMAP Representation of 26 patient samples of CD45+ cells colored by patient. (B) Principle component analysis of relative percentages of 6 major cell populations colored by disease condition. (C) Bar plot of relative percentages of 6 major cell populations.

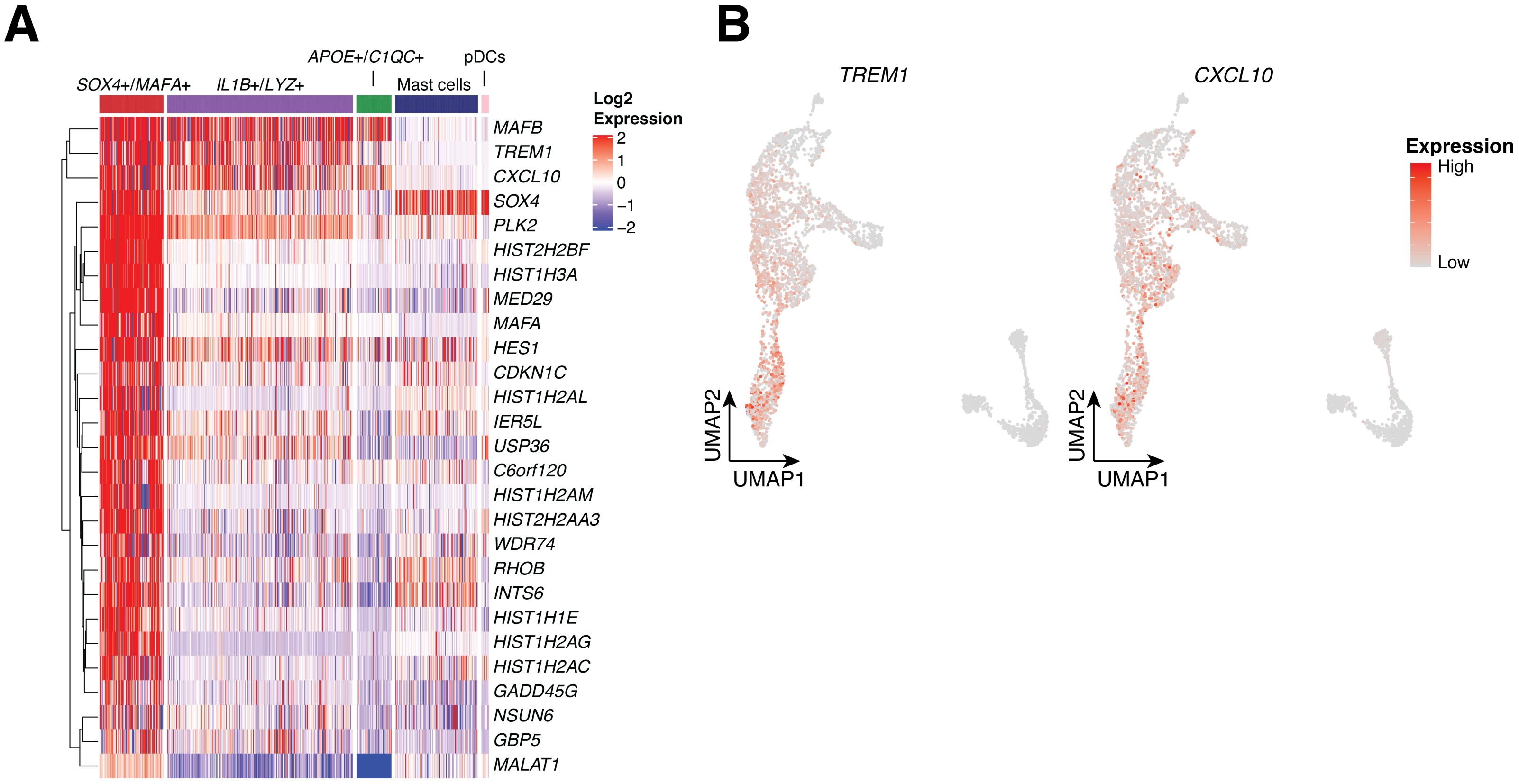
A
APOE+/C1QC+
pDCs
SOX4+/MAFA+
IL1B+/LYZ+
Mast cells
MAFB
TREM1
CXCL10
SOX4
PLK2
HIST2H2BF
HIST1H3A
MED29
MAFA
HES1
CDKN1C
HIST1H2AL
IER5L
USP36
C6orf120
HIST1H2AM
HIST2H2AA3
WDR74
RHOB
INTS6
HIST1H1E
HIST1H2AG
HIST1H2AC
GADD45G
NSUN6
GBP5
MALAT1
Log2
Expression
2
1
0
−1
−2
B
TREM1
CXCL10
Expression
High
Low
UMAP2
UMAP1
UMAP2
UMAP1

**Supplementary Figure 3.** (A) Heatmap of Monocyte/Macrophage 1 specific genes in each myeloid cell population. (B) Expression of *TREM1* and *CXCL10* in myeloid cell populations.

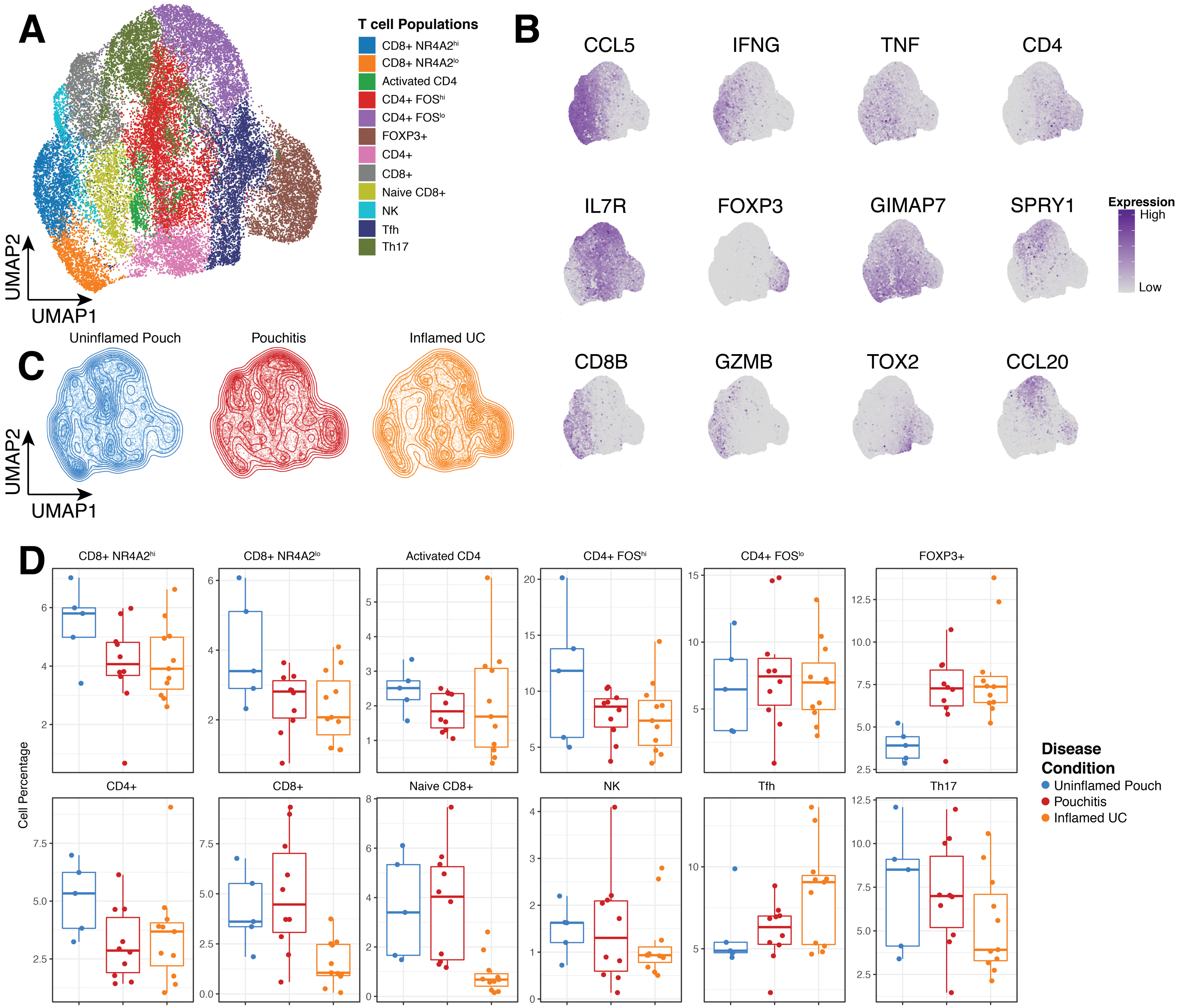

A
T cell Populations
CD8+ NR4A2hi
CD8+ NR4A2lo
Activated CD4
CD4+ FOShi
CD4+ FOSlo
FOXP3+
CD4+
CD8+
Naive CD8+
NK
Tfh
Th17
UMAP2
UMAP1
B
CCL5
IFNG
TNF
CD4
IL7R
FOXP3
GIMAP7
SPRY1
CD8B
GZMB
TOX2
CCL20
Expression
High
Low
C
Uninflamed Pouch
Pouchitis
Inflamed UC
D
Cell Percentage
Disease Condition
Uninflamed Pouch
Pouchitis
Inflamed UC

**Supplementary Figure 4.** (A) UMAP representation of 12 major T cell populations. (B) Expression of representative gene markers for each of the 12 T cell populations. (C) Contour plots of cell density on top of UMAP representation for uninflamed pouches, pouchitis and UC inflamed patient samples.

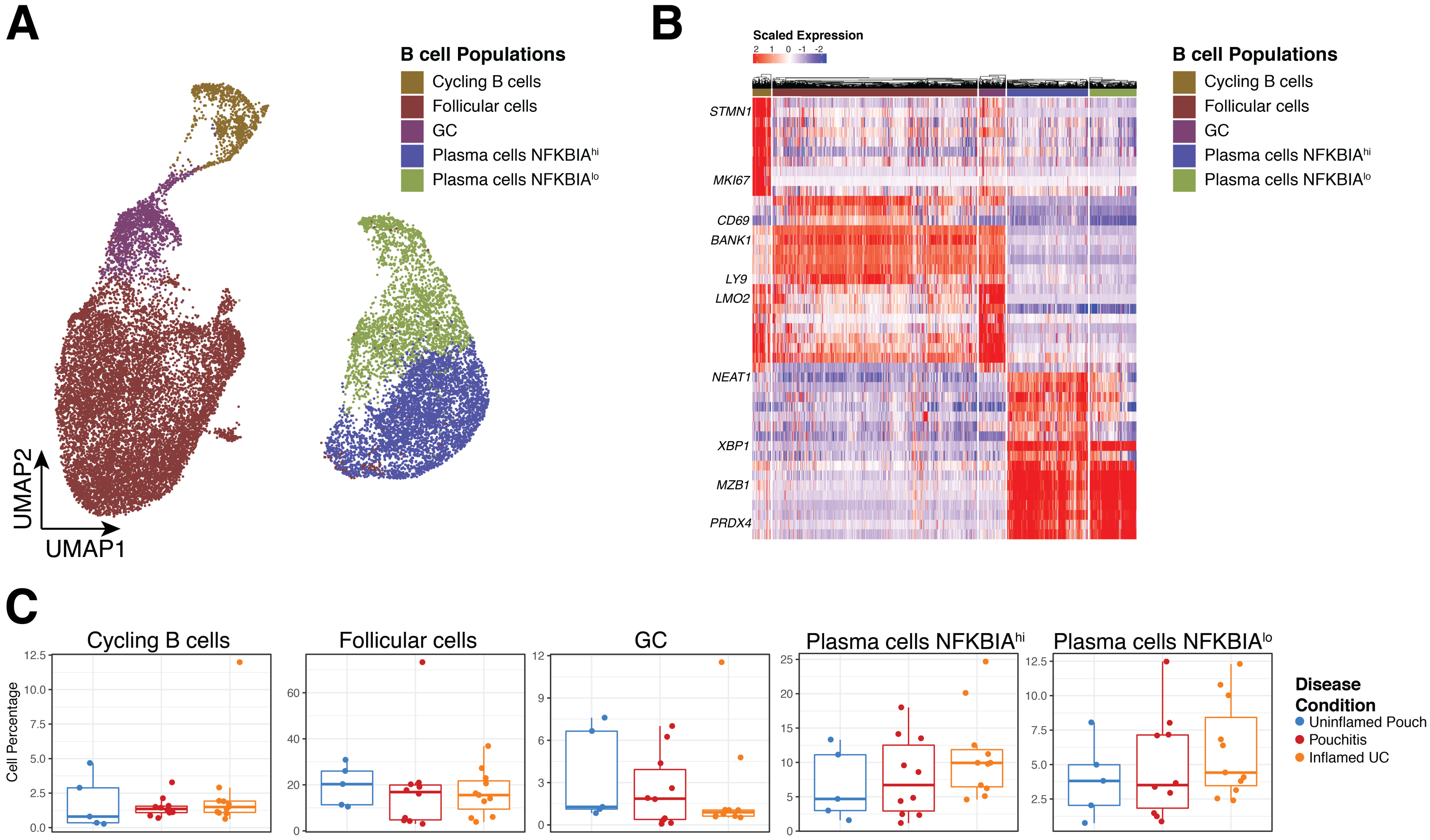

A
B cell Populations
Cycling B cells
Follicular cells
GC
Plasma cells NFKBIAhi
Plasma cells NFKBIAlo
UMAP2
UMAP1
B
Scaled Expression
2 1 0 -1 -2
STMN1
MKI67
CD69
BANK1
LY9
LMO2
NEAT1
XBP1
MZB1
PRDX4
B cell Populations
Cycling B cells
Follicular cells
GC
Plasma cells NFKBIAhi
Plasma cells NFKBIAlo
C
Cycling B cells
Follicular cells
GC
Plasma cells NFKBIAhi
Plasma cells NFKBIAlo
Cell Percentage
Disease Condition
Uninflamed Pouch
Pouchitis
Inflamed UC

**Supplementary Figure 5.** (A) UMAP representation of 5 major B cell populations. (B) Heatmap of marker genes for each of the 5 major B cell populations. (C) Relative percentages of each major B cell population in uninflamed pouches, pouchitis and UC inflamed patient samples.

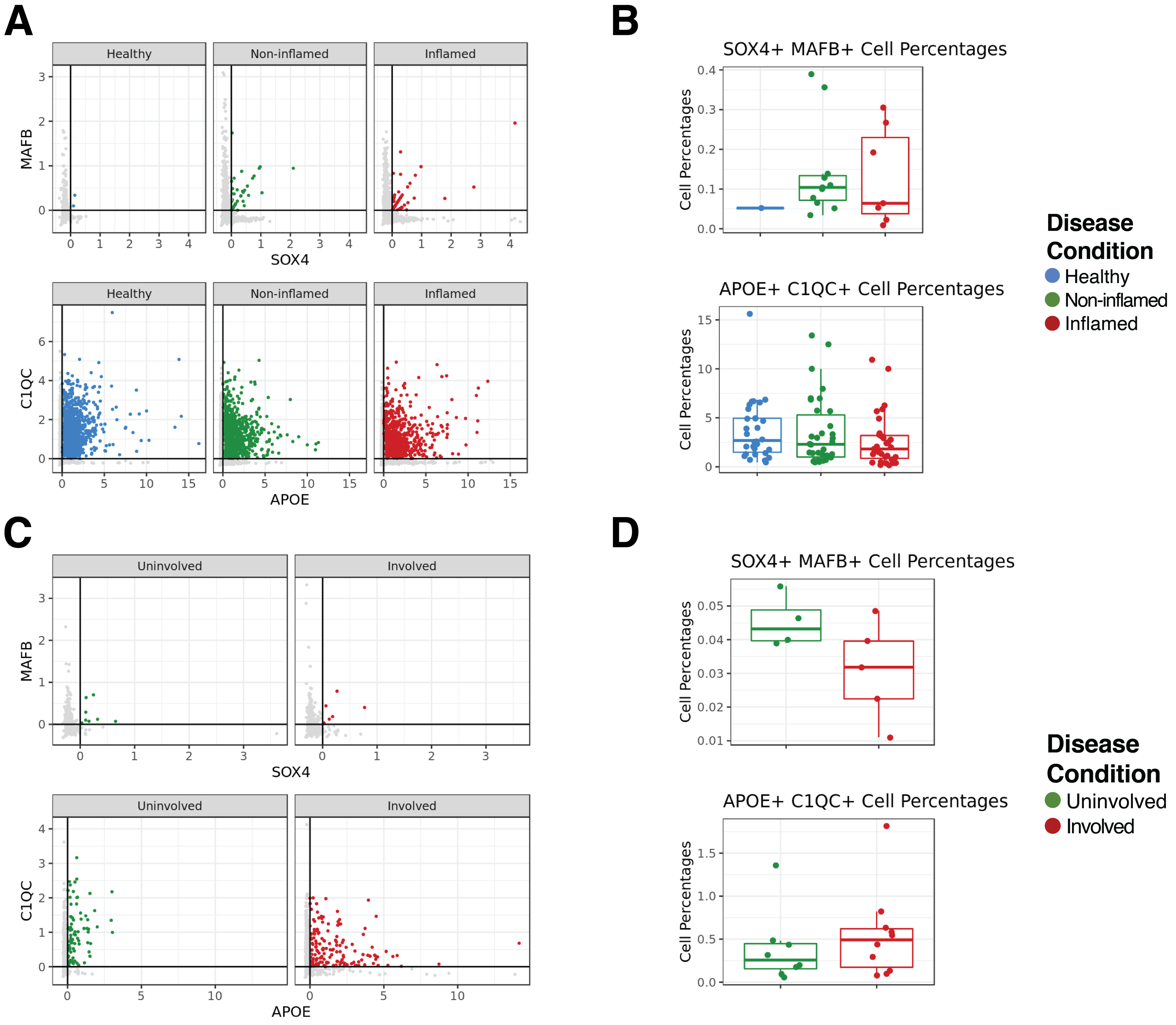

A
Healthy
Non-inflamed
Inflamed
MAFB
SOX4
C1QC
APOE
B
SOX4+ MAFB+ Cell Percentages
Cell Percentages
APOE+ C1QC+ Cell Percentages
Disease Condition
Healthy
Non-inflamed
Inflamed
C
Uninvolved
Involved
MAFB
SOX4
C1QC
APOE
D
SOX4+ MAFB+ Cell Percentages
APOE+ C1QC+ Cell Percentages
Cell Percentages
Disease Condition
Uninvolved
Involved

**Supplementary Figure 6.** (A) SOX4 by MAFB expression plot (top) and APOE by C1QC expression plot (bottom) in healthy, non-inflamed and inflamed ulcerative colitis patient samples from Smillie et al. (B) Relative percentages of SOX4+/MAFB+ myeloid cells (top) and APOE+/C1QC+ myeloid cells (bottom) in healthy, non-inflamed and inflamed ulcerative colitis patient samples from Smillie et al. (C) SOX4 by MAFB expression plot (top) and APOE by C1QC expression plot (bottom) in matched involved and uninvolved Crohns disease patient samples from Martins et al. (D) Relative percentages of SOX4+/MAFB+ myeloid cells (top) and APOE+/C1QC+ myeloid cells (bottom) in matched involved and uninvolved Crohns disease patient samples from Martins et al.

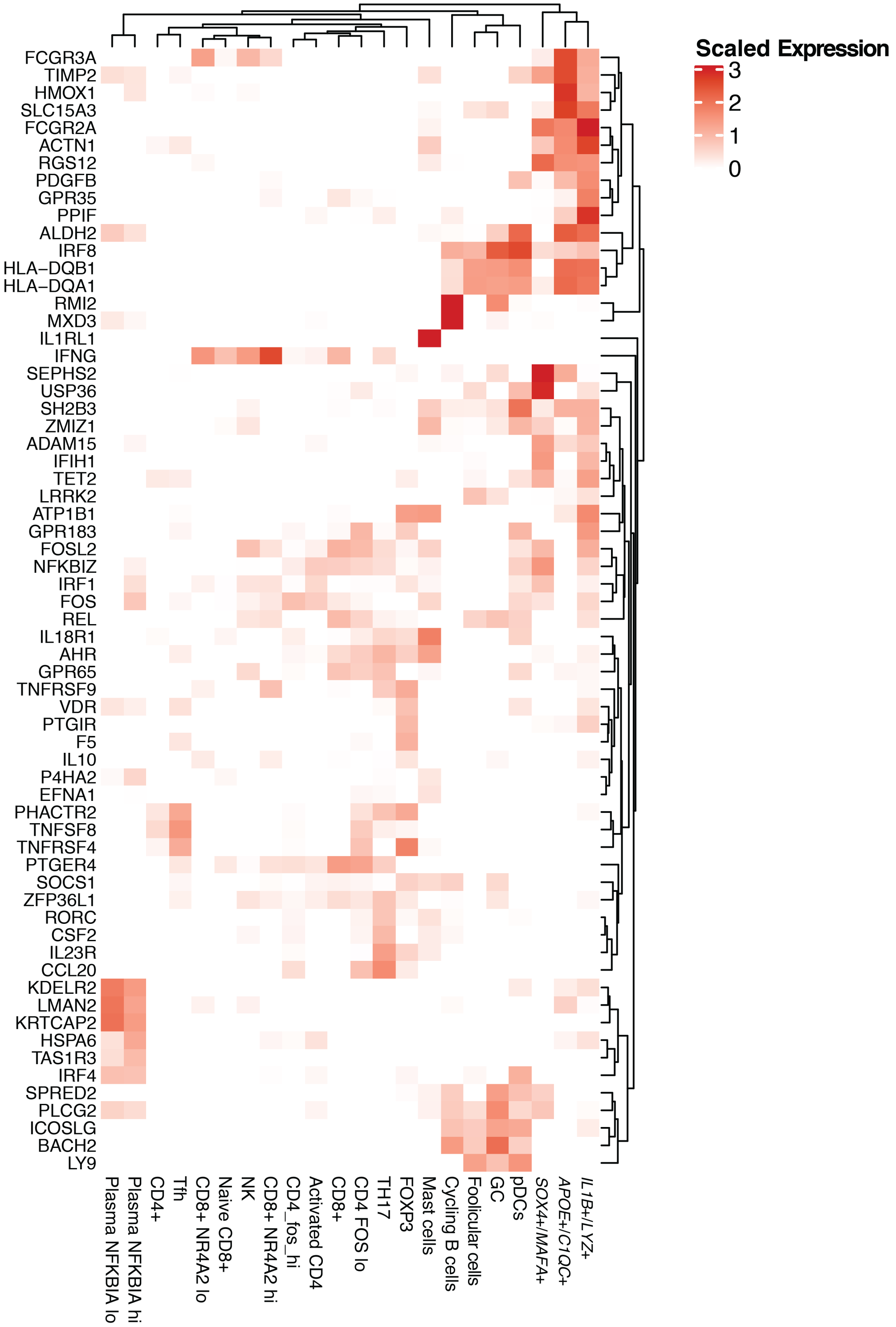

Scaled Expression
3
2
1
0
FCGR3A
TIMP2
HMOX1
SLC15A3
FCGR2A
ACTN1
RGS12
PDGFB
GPR35
PPIF
ALDH2
IRF8
HLA−DQB1
HLA−DQA1
RMI2
MXD3
IL1RL1
IFNG
SEPHS2
USP36
SH2B3
ZMIZ1
ADAM15
IFIH1
TET2
LRRK2
ATP1B1
GPR183
FOSL2
NFKBIZ
IRF1
FOS
REL
IL18R1
AHR
GPR65
TNFRSF9
VDR
PTGIR
F5
IL10
P4HA2
EFNA1
PHACTR2
TNFSF8
TNFRSF4
PTGER4
SOCS1
ZFP36L1
RORC
CSF2
IL23R
CCL20
KDELR2
LMAN2
KRTCAP2
HSPA6
TAS1R3
IRF4
SPRED2
PLCG2
ICOSLG
BACH2
LY9
Plasma NFKBIA lo
Plasma NFKBIA hi
CD4+
Tfh
CD8+ NR4A2 lo
Naive CD8+
NK
CD8+ NR4A2 hi
CD4_fos_hi
Activated CD4
CD8+
CD4 FOS lo
TH17
FOXP3
Mast cells
Cycling B cells
Follicular cells
GC
pDCs
SOX4+/MAFA+
APOE+/C1QC+
IL1B+/LYZ+

**Supplementary Figure 7.** Average expression level of 62 GWAS genes in each of the 22 immune cell types identified by our scRNA-Seq analysis.

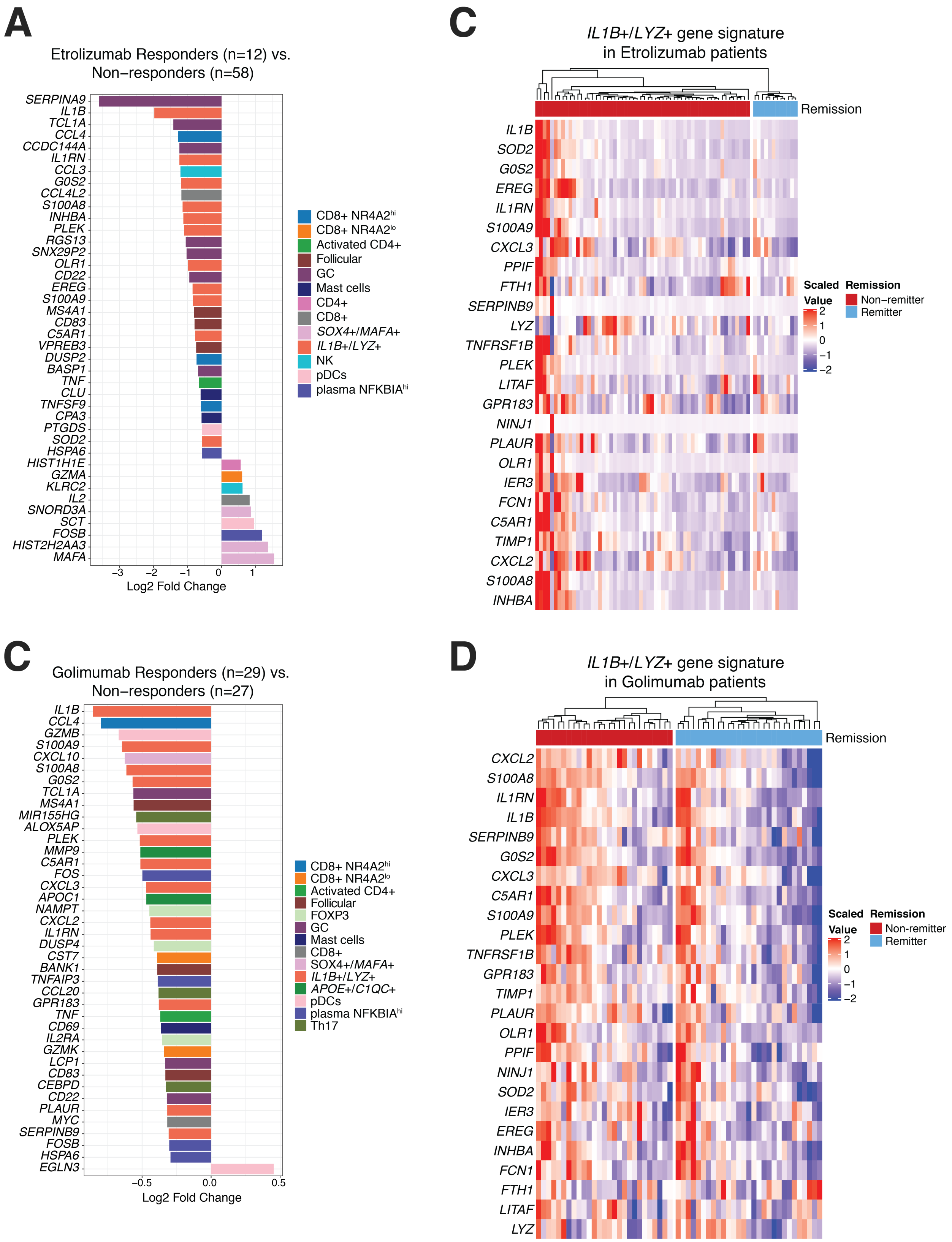
A
Etrolizumab Responders (n=12) vs.
Non−responders (n=58)
SERPINA9
IL1B
TCL1A
CCL4
CCDC144A
IL1RN
CCL3
G0S2
CCL4L2
S100A8
INHBA
PLEK
RGS13
SNX29P2
OLR1
CD22
EREG
S100A9
MS4A1
CD83
C5AR1
VPREB3
DUSP2
BASP1
TNF
CLU
TNFSF9
CPA3
PTGDS
SOD2
HSPA6
HIST1H1E
GZMA
KLRC2
IL2
SNORD3A
SCT
FOSB
HIST2H2AA3
MAFA
−3
−2
−1
0
1
Log2 Fold Change
CD8+ NR4A2hi
CD8+ NR4A2lo
Activated CD4+
Follicular
GC
Mast cells
CD4+
CD8+
SOX4+/MAFA+
IL1B+/LYZ+
NK
pDCs
plasma NFKBIAhi
C
IL1B+/LYZ+ gene signature
in Etrolizumab patients
Remission
IL1B
SOD2
G0S2
EREG
IL1RN
S100A9
CXCL3
PPIF
FTH1
SERPINB9
LYZ
TNFRSF1B
PLEK
LITAF
GPR183
NINJ1
PLAUR
OLR1
IER3
FCN1
C5AR1
TIMP1
CXCL2
S100A8
INHBA
Scaled
Value
2
1
0
−1
−2
Remission
Non−remitter
Remitter
C
Golimumab Responders (n=29) vs.
Non−responders (n=27)
IL1B
CCL4
GZMB
S100A9
CXCL10
S100A8
G0S2
TCL1A
MS4A1
MIR155HG
ALOX5AP
PLEK
MMP9
C5AR1
FOS
CXCL3
APOC1
NAMPT
CXCL2
IL1RN
DUSP4
CST7
BANK1
TNFAIP3
CCL20
GPR183
TNF
CD69
IL2RA
GZMK
LCP1
CD83
CEBPD
CD22
PLAUR
MYC
SERPINB9
FOSB
HSPA6
EGLN3
−0.5
0.0
0.5
Log2 Fold Change
CD8+ NR4A2hi
CD8+ NR4A2lo
Activated CD4+
Follicular
FOXP3
GC
Mast cells
CD8+
SOX4+/MAFA+
IL1B+/LYZ+
APOE+/C1QC+
pDCs
plasma NFKBIAhi
Th17
D
IL1B+/LYZ+ gene signature
in Golimumab patients
Remission
CXCL2
S100A8
IL1RN
IL1B
SERPINB9
G0S2
CXCL3
C5AR1
S100A9
PLEK
TNFRSF1B
GPR183
TIMP1
PLAUR
OLR1
PPIF
NINJ1
SOD2
IER3
EREG
INHBA
FCN1
FTH1
LITAF
LYZ
Scaled
Value
2
1
0
−1
−2
Remission
Non−remitter
Remitter

**Supplementary Figure 8.** (A) Log fold change of differentially expressed genes in responders versus non-responders to Etrolizumab and Golimumab (C). (B) Gene expression of Monocyte/Macrophage 2 signature genes in responders and non-responders to Etrolizumab and Golimumab (D)
