## Supplementary Table 1 for "Transcriptional Atlas of Ileal-Anal Pouch Immune Cells from Ulcerative Colitis Patients"

| | A | B | C | D | E | F | G | H | I | J | K | L | M | N | O | P | Q | R | S | T | U | V | W | X |
|---|---|---|---|---|---|---|---|---|---|---|---|---|---|---|---|---|---|---|---|---|---|---|---|---|
| 1 | GeneSymbol | act_CD8_NR4A2 | act_CD8_NR4A2 | activated_CD4 | CD4_fos_hi | CD4_fos_lo | Foxp3_cells | memory_CD4 | memory_CD8 | naive_CD8 | NK_cells | Tfh_cells | Th17_cells | cycling_b_cells | follicular_cells | gc_cells | plasma_NFKBIA | plasma_NFKBIA | mast_cells | pdcs | SOX4/MAFA | IL1B/LYZ | APOE/C1QC | CellState |
| 2 | IFNG | 2.244611002 | 1.403850675 | 0.573814488 | 0.439093585 | 0.250843841 | 0.084668338 | 0.184825238 | 1.117558763 | 0.902425025 | 1.144390611 | 0.162779636 | 0.973203215 | 0.335451864 | 0.037200876 | 0.032902389 | 0.262613131 | 0.042503032 | 0.079420129 | 0.006215151 | 0.043019612 | 0.054429794 | 0.074859121 | act_CD8_NR4A2_hi |
| 3 | CCL4 | 2.987128837 | 2.392236308 | 0.801131132 | 0.355758223 | 0.238554862 | 0.168567553 | 0.314403493 | 1.895640632 | 1.237494595 | 2.097577404 | 0.143758577 | 1.622620667 | 0.549041411 | 0.427590794 | 0.362473705 | 0.515096311 | 0.20265267 | 0.347466871 | 0.643934274 | 1.314623974 | 2.020781531 | 1.522062235 | act_CD8_NR4A2_hi |
| 4 | CRTAM | 0.93107243 | 0.438457143 | 0.088949926 | 0.026178513 | 0.021775673 | 0.012722354 | 0.044840396 | 0.53452331 | 0.32397665 | 0.572292849 | 0.014897358 | 0.098666839 | 0.049177546 | 0.007437488 | 0.00981165 | 0.059605511 | 0.010782749 | 0.040005527 | 0.007606819 | 0.013143526 | 0.079392293 | 0.05518 | act_CD8_NR4A2_hi |
| 5 | TNFSF9 | 1.044677664 | 0.321205549 | 0.298137872 | 0.181790134 | 0.129635453 | 0.098513504 | 0.040486429 | 0.599826119 | 0.24269284 | 0.38489425 | 0.055399848 | 0.4251822 | 0.157219347 | 0.328277036 | 0.236164295 | 0.197918879 | 0.033674126 | 0.104747292 | 0.320575486 | 0.236529458 | 0.192047039 | 0.151280081 | act_CD8_NR4A2_hi |
| 6 | CCL5 | 3.444115846 | 3.378924562 | 1.236364896 | 1.513255923 | 0.918279576 | 0.156041508 | 1.085576306 | 3.501887769 | 3.543680438 | 2.989754008 | 0.140445358 | 2.081353852 | 0.54265056 | 0.120766562 | 0.118767462 | 0.567984681 | 0.136374564 | 0.221925931 | 0.07018882 | 0.248558178 | 0.261336402 | 0.186393082 | act_CD8_NR4A2_hi |
| 7 | DUSP2 | 2.634100253 | 1.121277687 | 1.242847166 | 1.203932246 | 1.610516903 | 1.625127243 | 0.298855893 | 1.87369896 | 0.850963032 | 1.986714783 | 1.269015014 | 1.668876531 | 0.840981807 | 1.327713957 | 1.140254474 | 1.360762975 | 0.266785855 | 0.700449228 | 0.444215715 | 0.882501517 | 1.599565929 | 0.687947762 | act_CD8_NR4A2_hi |
| 8 | APOBEC3G | 0.883699296 | 0.970532901 | 0.103700204 | 0.160588059 | 0.198740044 | 0.233345653 | 0.154440412 | 0.304416292 | 0.304563755 | 0.861015866 | 0.142279061 | 0.388452587 | 0.338897307 | 0.275876164 | 0.330138179 | 0.232643508 | 0.195861394 | 0.163911304 | 0.228238168 | 0.036109524 | 0.149438248 | 0.131933216 | act_CD8_NR4A2_hi |
| 9 | TUBA4A | 1.026288976 | 0.593061733 | 0.43552064 | 0.477090835 | 0.486525312 | 0.425477859 | 0.399684049 | 0.614316729 | 0.528311389 | 0.553922787 | 0.496129414 | 0.451466189 | 0.774185225 | 0.298262774 | 0.527402088 | 0.604881104 | 0.377352728 | 0.129446706 | 0.01531479 | 0.238685944 | 0.143553152 | 0.118079117 | act_CD8_NR4A2_hi |
| 10 | TMEM2 | 0.985326859 | 0.437756397 | 0.475930399 | 0.313139664 | 0.434196838 | 0.674826505 | 0.283754747 | 0.479058504 | 0.261106075 | 0.763109436 | 0.740181552 | 0.516006588 | 0.237811271 | 0.274421045 | 0.131150762 | 0.188908479 | 0.028714276 | 0.161766221 | 0.057801507 | 0.29533896 | 0.202156311 | 0.101010831 | act_CD8_NR4A2_hi |
| 11 | GZMK | 1.761324082 | 1.958340291 | 0.050020383 | 0.072729803 | 0.087167639 | 0.026764884 | 0.140546956 | 0.109490212 | 0.134648057 | 0.822559648 | 0.051560822 | 0.282233432 | 0.088489628 | 0.011071795 | 0.013098323 | 0.096926528 | 0.018851876 | 0.014112459 | 0 | 0.005756233 | 0.01954866 | 0.01896319 | act_CD8_NR4A2_lo |
| 12 | NKG7 | 2.276473277 | 2.590114926 | 0.124060245 | 0.197998139 | 0.174905412 | 0.073355463 | 0.199568338 | 1.351937482 | 1.400604243 | 2.441061667 | 0.040961581 | 1.024510564 | 0.379027349 | 0.03052754 | 0.025889848 | 0.178113367 | 0.029629103 | 0.190267589 | 0.349802633 | 0.034971754 | 0.177496222 | 0.169029647 | act_CD8_NR4A2_lo |
| 13 | GZMH | 0.803909947 | 1.09720562 | 0.02238524 | 0.009668754 | 0.012235582 | 0.018652109 | 0.020568473 | 0.078177461 | 0.104895924 | 0.345367394 | 0.01109429 | 0.078503086 | 0.090375052 | 0.00557443 | 0.003443968 | 0.044259183 | 0.008467459 | 0.01244528 | 0.069251335 | 0.003181275 | 0.008733116 | 0.011736808 | act_CD8_NR4A2_lo |
| 14 | GZMA | 1.60641028 | 2.092630665 | 0.061120419 | 0.167697079 | 0.184141316 | 0.100965595 | 0.163984911 | 0.665318286 | 0.575145353 | 1.458642557 | 0.067588191 | 1.427760354 | 0.442016568 | 0.014294911 | 0.013966702 | 0.182010922 | 0.025411123 | 0.044873052 | 0.008107145 | 0.021675563 | 0.04070269 | 0.02935986 | act_CD8_NR4A2_lo |
| 15 | CST7 | 1.554560689 | 1.688779121 | 0.083397043 | 0.235384637 | 0.265227545 | 0.450043622 | 0.202022551 | 0.676309204 | 0.560536247 | 1.167447379 | 0.186235136 | 0.619190022 | 0.17852482 | 0.015702928 | 0.013724765 | 0.134701711 | 0.018472391 | 0.05010377 | 0.031552794 | 0.034189548 | 0.285653506 | 0.123721885 | act_CD8_NR4A2_lo |
| 16 | CMC1 | 0.831371819 | 1.055826859 | 0.063907143 | 0.147112626 | 0.151245873 | 0.206740439 | 0.147926958 | 0.190619202 | 0.197702571 | 0.933911593 | 0.175805926 | 0.248334004 | 0.322958703 | 0.116284059 | 0.170666496 | 0.2045543 | 0.178822018 | 0.233637863 | 0.24000781 | 0.051172178 | 0.185629082 | 0.245266398 | act_CD8_NR4A2_lo |
| 17 | KLRG1 | 0.776521414 | 1.071160987 | 0.114765576 | 0.156187641 | 0.151926537 | 0.058818264 | 0.189991401 | 0.05629074 | 0.079381642 | 0.266805905 | 0.052461324 | 0.273468483 | 0.057350907 | 0.017168187 | 0.029148109 | 0.05622722 | 0.017187131 | 0.351201625 | 0.013496281 | 0.034840824 | 0.016424813 | 0.026420794 | act_CD8_NR4A2_lo |
| 18 | SAMD3 | 0.625425724 | 0.968430499 | 0.119825126 | 0.114046667 | 0.116307715 | 0.04574189 | 0.158166561 | 0.115065116 | 0.135822513 | 0.652973908 | 0.02721933 | 0.124825478 | 0.05486122 | 0.004723356 | 0.004901882 | 0.036611668 | 0.007404303 | 0.030693501 | 0.009420133 | 0.001339351 | 0.008691593 | 0.010122555 | act_CD8_NR4A2_lo |
| 19 | PRF1 | 0.433701968 | 0.741483105 | 0.039168275 | 0.046653499 | 0.040740894 | 0.076205249 | 0.052026366 | 0.157200999 | 0.174567255 | 0.678918905 | 0.041435224 | 0.269967739 | 0.104051621 | 0.003273327 | 0.004335406 | 0.041913041 | 0.006489721 | 0.021817461 | 0.008107145 | 0.001339351 | 0.011489032 | 0.012382725 | act_CD8_NR4A2_lo |
| 20 | HNRNPA2B1 | 2.13899975 | 2.083069062 | 2.916638186 | 2.102507526 | 1.891014612 | 2.140493827 | 1.954621127 | 2.072113564 | 2.097688904 | 2.257031779 | 2.057042096 | 2.13613073 | 2.638701351 | 2.150726385 | 2.283707154 | 2.037429377 | 1.469909044 | 2.229972076 | 2.356179466 | 2.507133757 | 2.013965885 | 1.544297852 | activated_CD4 |
| 21 | PPP1R15A | 2.108122183 | 1.425055555 | 2.869101485 | 2.347514192 | 2.078077624 | 2.112599606 | 1.193185257 | 2.307276108 | 1.988793447 | 2.21217815 | 1.76961956 | 2.275265979 | 1.434941645 | 1.964066167 | 1.530443624 | 2.688711107 | 1.169222233 | 2.187928213 | 2.502258406 | 3.034865971 | 2.313374087 | 1.19291151 | activated_CD4 |
| 22 | SLC38A2 | 1.13538793 | 0.947733096 | 2.01681773 | 1.06927749 | 0.96268991 | 1.101163654 | 0.964633094 | 1.033673838 | 1.06209163 | 1.152505708 | 0.961270515 | 1.044870117 | 0.734337406 | 0.799020475 | 0.677064543 | 1.489584159 | 1.013659406 | 1.097507754 | 1.021801023 | 1.176704688 | 0.918773731 | 0.577691119 | activated_CD4 |
| 23 | HSPA1B | 2.574017978 | 2.075306524 | 3.364743559 | 3.254335932 | 1.409568429 | 1.811863027 | 1.796629919 | 2.519664333 | 3.009370447 | 2.263653398 | 1.652917812 | 2.12659718 | 1.140345644 | 2.207983429 | 1.066790576 | 3.013958572 | 1.331903546 | 2.517643836 | 2.845728605 | 2.728447266 | 2.500942098 | 1.554776075 | activated_CD4 |
| 24 | PTCH2 | 0.06586581 | 0.067487863 | 0.531303376 | 0.088917935 | 0.041013589 | 0.051655417 | 0.074608746 | 0.0629585 | 0.117775242 | 0.121145246 | 0.049257759 | 0.093283308 | 0.073262888 | 0.057549346 | 0.046831056 | 0.445326653 | 0.22896739 | 0.146726763 | 0.176575528 | 0.312492029 | 0.076792517 | 0.057184343 | activated_CD4 |
| 25 | PHYH | 0.333083791 | 0.31578949 | 0.971129734 | 0.498634581 | 0.185717236 | 0.17226995 | 0.295602649 | 0.353305512 | 0.561115173 | 0.30722733 | 0.154829438 | 0.253081124 | 0.148874597 | 0.124685476 | 0.101545947 | 0.340596426 | 0.166281212 | 0.313161761 | 0.368302908 | 0.168094651 | 0.103152081 | 0.084094286 | activated_CD4 |
| 26 | SAT1 | 1.729233642 | 1.314648562 | 2.439426812 | 1.600814579 | 1.56578419 | 2.460544693 | 1.202860453 | 1.67485276 | 1.518153523 | 1.459962556 | 1.483455204 | 1.584676132 | 1.210586273 | 1.550351801 | 1.28540777 | 1.921012691 | 1.123535146 | 1.959871411 | 1.730257502 | 4.087332341 | 3.573436675 | 2.976035562 | activated_CD4 |
| 27 | GADD45B | 1.194387143 | 1.066516035 | 1.845931138 | 1.32464214 | 0.832505368 | 0.825984561 | 0.688923857 | 1.066175436 | 1.421793201 | 1.612365165 | 0.5409261 | 0.905417308 | 0.963706909 | 1.454258953 | 1.317512788 | 1.728248071 | 0.745142153 | 1.439972483 | 1.062058832 | 2.369946287 | 1.855358303 | 1.119382953 | activated_CD4 |
| 28 | TSPYL2 | 1.166594845 | 0.672705602 | 1.752083524 | 1.038057078 | 0.857295194 | 1.099692497 | 0.685016328 | 1.312126253 | 0.969240536 | 0.996982403 | 0.967372227 | 0.941700536 | 0.34939542 | 0.544800481 | 0.37997151 | 0.707756138 | 0.331857802 | 0.851973441 | 1.32695366 | 0.983497078 | 0.370037373 | 0.186200323 | activated_CD4 |
| 29 | KMT2E-AS1 | 0.437694487 | 0.529105621 | 1.081868738 | 0.516997623 | 0.310651721 | 0.399918661 | 0.527935442 | 0.411916326 | 0.551457054 | 0.47028322 | 0.350235671 | 0.460523033 | 0.35846564 | 0.395024115 | 0.542300205 | 0.799688282 | 0.484678329 | 0.382658352 | 0.315940081 | 0.91239569 | 0.358258528 | 0.277376649 | activated_CD4 |
| 30 | TNF | 0.681420419 | 0.931127738 | 1.463511306 | 0.911456314 | 0.341864623 | 0.254962717 | 0.824227016 | 0.490362312 | 0.894291155 | 0.364116439 | 0.360008084 | 0.778949723 | 0.436494418 | 0.501191141 | 0.565011257 | 0.206630338 | 0.063456658 | 0.229648569 | 0.069191819 | 0.959688771 | 0.509811301 | 0.352541816 | activated_CD4 |
| 31 | NABP1 | 0.785962233 | 0.572824454 | 1.51767148 | 1.133429202 | 0.700158934 | 1.098262916 | 0.707998263 | 0.987147657 | 0.974259462 | 0.555995323 | 0.867260748 | 0.776403129 | 0.385711799 | 0.500051444 | 0.376142615 | 0.66017162 | 0.285907923 | 0.694693894 | 0.40213815 | 1.199902034 | 1.071998662 | 0.592150444 | activated_CD4 |
| 32 | VMP1 | 1.314750698 | 1.176182443 | 1.829219487 | 1.265545828 | 1.035681586 | 1.641834052 | 1.062878226 | 1.256171145 | 1.390530575 | 1.019167275 | 1.319642106 | 1.258535625 | 0.951120499 | 0.497786625 | 0.680737945 | 0.76573642 | 0.327890188 | 1.075517802 | 0.879264468 | 1.503015473 | 1.612160728 | 1.008931793 | activated_CD4 |
| 33 | BRD2 | 1.039546944 | 0.675363269 | 1.311723829 | 0.684669633 | 0.892752563 | 0.945808142 | 0.488793395 | 0.826773535 | 0.64723147 | 1.07953889 | 0.728897227 | 0.94078887 | 0.8270975 | 0.670056723 | 0.869737581 | 1.123285751 | 0.635921435 | 1.029223384 | 0.950048524 | 1.244538663 | 0.729793488 | 0.437808515 | activated_CD4 |
| 34 | POLR2A | 0.410607973 | 0.269205358 | 0.838738333 | 0.278062737 | 0.311599746 | 0.378332115 | 0.237332944 | 0.354530401 | 0.282796255 | 0.400778365 | 0.325628726 | 0.391761997 | 0.43827621 | 0.210595847 | 0.339870843 | 0.316712931 | 0.1785405 | 0.360303325 | 0.538422881 | 0.739598337 | 0.248984317 | 0.181375971 | activated_CD4 |
| 35 | DNAJB1 | 3.188069309 | 2.335005425 | 3.408020104 | 3.442037154 | 2.071410075 | 2.385116165 | 1.736111854 | 2.786705327 | 2.932851364 | 2.615555068 | 2.007228904 | 2.530829026 | 1.448350787 | 2.444096213 | 1.547811037 | 2.956345049 | 1.178386855 | 2.249016276 | 2.40090494 | 2.592343349 | 2.363811553 | 1.403269727 | CD4_fos_hi |
| 36 | DUSP1 | 2.621541268 | 1.56366667 | 2.827895784 | 3.048272381 | 1.992461741 | 2.336684606 | 1.212824241 | 2.412262002 | 2.16756629 | 2.372727221 | 2.27104672 | 2.387035228 | 1.238379716 | 2.230822796 | 1.191100119 | 2.621638895 | 1.073172048 | 2.895761198 | 2.334273087 | 1.993014733 | 2.849201179 | 1.849254761 | CD4_fos_hi |
| 37 | HSP90AA1 | 4.357263339 | 3.25702039 | 4.420657979 | 4.419400665 | 3.529235959 | 3.727537231 | 2.690069039 | 4.137538632 | 3.859198651 | 3.906330821 | 3.495672703 | 3.858220297 | 3.31974439 | 3.935839549 | 3.02607347 | 4.487806868 | 2.375592177 | 3.650387038 | 4.71827118 | 3.362215031 | 3.818478896 | 2.936630967 | CD4_fos_hi |
| 38 | HSPA1A | 2.954172669 | 2.058533659 | 3.146663467 | 3.302209838 | 1.795871711 | 2.264408449 | 1.527287371 | 2.918401041 | 2.915953355 | 2.797807559 | 1.899692703 | 2.555763093 | 1.235645261 | 2.194465575 | 0.955502033 | 3.248901369 | 1.441279919 | 2.910491708 | 3.262117212 | 2.516006311 | 3.320722183 | 2.104744873 | CD4_fos_hi |
| 39 | HSPH1 | 1.965020301 | 0.888791393 | 2.20432098 | 2.039915819 | 1.431441178 | 1.505134084 | 0.523058921 | 2.102398236 | 1.572200979 | 1.694929991 | 1.328444672 | 1.727833033 | 1.135061083 | 1.65027988 | 1.031271119 | 2.183721003 | 0.5254972 | 1.523340041 | 2.132711183 | 1.419942297 | 1.926235471 | 1.01731957 | CD4_fos_hi |
| 40 | RGS2 | 0.7767608 | 0.459928285 | 1.111962876 | 1.098189088 | 0.664821442 | 0.476080973 | 0.44809236 | 0.977151912 | 0.896501661 | 0.659340758 | 0.724125143 | 1.111118079 | 0.712033418 | 0.461864985 | 0.773008545 | 1.003317622 | 0.38941516 | 0.909269609 | 2.068350986 | 2.079318401 | 1.512778041 | 0.932885927 | CD4_fos_hi |
| 41 | LMNA | 0.613517359 | 0.126281032 | 0.340716719 | 0.395115469 | 1.204106594 | 0.798966191 | 0.088521896 | 0.808356518 | 0.182619006 | 0.578589043 | 0.645226863 | 0.551926849 | 0.244771222 | 0.227731666 | 0.069296674 | 0.71314479 | 0.258724147 | 1.575514221 | 0.160592647 | 0.327045218 | 0.950046591 | 0.437079542 | CD4_fos_lo |
| 42 | VIM | 1.940945715 | 1.656257014 | 0.956712078 | 1.923795361 | 2.542865376 | 1.354947761 | 1.396070986 | 2.51217313 | 2.060178896 | 2.003762058 | 0.974125648 | 1.923688245 | 1.611783591 | 0.863948748 | 0.380153516 | 1.918673803 | 1.650312181 | 3.050086919 | 2.509330486 | 1.938445973 | 3.268698242 | 2.208683828 | CD4_fos_lo |
| 43 | IL7R | 0.854363897 | 0.640857429 | 1.213127359 | 2.580183985 | 2.681235568 | 1.240904901 | 2.17231709 | 2.514620113 | 2.16775008 | 0.779926977 | 1.199223283 | 2.1032131 | 0.27969099 | 0.071551556 | 0.060141133 | 0.528024876 | 0.084004545 | 0.750127143 | 0.061683194 | 0.31138226 | 0.674266048 | 0.495728024 | CD4_fos_lo |
| 44 | EZR | 1.222435194 | 0.587967107 | 0.539282694 | 0.620394966 | 1.588937753 | 1.253171306 | 0.370332779 | 1.513321835 | 0.595195125 | 1.000597447 | 1.039812881 | 1.093815631 | 1.581078199 | 1.111446418 | 1.681770763 | 0.858891224 | 0.333749029 | 0.424700546 | 1.537367504 | 0.290409951 | 0.926986176 | 0.417456078 | CD4_fos_lo |
| 45 | CD55 | 0.334638075 | 0.134245274 | 0.26265487 | 0.347043298 | 0.791788253 | 0.215634128 | 0.193156449 | 0.798935996 | 0.336681346 | 0.659798286 | 0.193182585 | 0.410848144 | 0.331332735 | 0.962335408 | 0.509497054 | 0.459785823 | 0.265256039 | 0.57653787 | 0.714550316 | 0.383334868 | 1.185386443 | 0.297689318 | CD4_fos_lo |
| 46 | MYADM | 1.253678793 | 0.419900243 | 0.808167562 | 1.194529257 | 1.572093143 | 0.668038641 | 0.231742211 | 1.788578273 | 0.83080826 | 1.180571 | 0.501241548 | 1.270732429 | 0.395166132 | 0.586533625 | 0.216201388 | 0.840012192 | 0.218392843 | 1.193246584 | 1.002206962 | 0.224216239 | 0.939393547 | 0.378089842 | CD4_fos_lo |
| 47 | PABPC1 | 1.686246003 | 1.188012978 | 1.448167076 | 1.82592909 | 2.281699219 | 1.692416872 | 1.548975147 | 2.107748104 | 1.632519395 | 1.926971868 | 1.852797117 | 1.897335711 | 1.850437642 | 1.804917961 | 2.212617764 | 1.363472625 | 1.026775579 | 1.650815306 | 2.09223934 | 1.057075571 | 2.201442131 | 1.779719946 | CD4_fos_lo |
| 48 | KDM6B | 1.044671487 | 0.259309733 | 0.681475783 | 0.760741418 | 1.383620425 | 0.99956722 | 0.167509837 | 1.360507252 | 0.444994403 | 1.107180233 | 0.743154149 | 1.086608761 | 0.421398366 | 0.978549905 | 0.547450161 | 0.554686737 | 0.094209286 | 0.907709038 | 1.429558173 | 0.743557676 | 1.528594569 | 0.535215337 | CD4_fos_lo |
| 49 | ARL4C | 1.357758461 | 1.377753163 | 0.876755667 | 1.214241991 | 1.794500341 | 1.185641883 | 1.01137115 | 1.278095934 | 0.992380099 | 0.928184137 | 1.121687838 | 1.41830801 | 0.474680017 | 0.343510432 | 0.119640168 | 0.389145198 | 0.084885678 | 0.319337403 | 2.073751566 | 0.333088274 | 1.18824055 | 0.833466016 | CD4_fos_lo |
| 50 | TSC22D3 | 1.483657135 | 0.78123285 | 1.428819488 | 1.549528047 | 2.198384191 | 1.969085611 | 0.851622141 | 1.945240563 | 0.971980117 | 1.457728069 | 1.591997303 | 1.657307651 | 0.793243892 | 1.332750899 | 0.848646951 | 1.614621389 | 0.75626078 | 1.401868365 | 1.161354706 | 1.123917071 | 1.074853696 | 0.689781052 | CD4_fos_lo |
| 51 | S100A10 | 1.106745914 | 1.159917434 | 0.260269147 | 1.130144553 | 1.484741024 | 0.767306802 | 0.919871256 | 1.343192098 | 1.38113589 | 1.126446603 | 0.641467972 | 0.870713769 | 1.387342435 | 0.355058996 | 0.899745993 | 0.448414115 | 0.254337434 | 1.061937159 | 0.516317687 | 0.571301792 | 1.970157914 | 1.206286974 | CD4_fos_lo |
| 52 | CRIP1 | 1.141925308 | 0.988267394 | 0.597271757 | 1.032645644 | 1.677569289 | 1.057893293 | 0.723284139 | 1.516788418 | 1.020269469 | 1.238183738 | 0.988552774 | 1.408908511 | 1.368387284 | 0.48931779 | 0.437130192 | 0.657923943 | 0.297110459 | 0.571196402 | 0.624230849 | 0.219803877 | 0.751995682 | 0.369402872 | CD4_fos_lo |
| 53 | PDE4D | 0.370679939 | 0.184076669 | 0.229148797 | 0.383915104 | 0.822137723 | 0.359472528 | 0.224940158 | 0.676088964 | 0.31203152 | 0.392389523 | 0.455244131 | 0.953285824 | 0.277963212 | 0.079620008 | 0.158554572 | 0.226495822 | 0.060690698 | 0.233629568 | 0.071719726 | 0.057355562 | 0.131473094 | 0.038574749 | CD4_fos_lo |
| 54 | CCR7 | 0.162341704 | 0.08479734 | 0.090441868 | 0.20945973 | 0.509516995 | 0.281220812 | 0.302388884 | 0.076993221 | 0.054841482 | 0.077300766 | 0.39688237 | 0.084001516 | 0.077919593 | 0.470268711 | 0.059139919 | 0.060014516 | 0.012461666 | 0.042155252 | 0.208193699 | 0.054495059 | 0.380056258 | 0.0665697 | CD4_fos_lo |
| 55 | KLF2 | 1.168239936 | 0.939956253 | 0.748550235 | 0.890853474 | 1.199444066 | 0.540601959 | 0.60676163 | 0.422493227 | 0.332924168 | 0.873763221 | 0.574441986 | 0.469821715 | 0.42330642 | 1.626784236 | 0.318260214 | 1.487835257 | 0.569144466 | 1.12202692 | 0.508161509 | 1.697422437 | 0.867972085 | 0.410808256 | CD4_fos_lo |
| 56 | S100A11 | 0.607682139 | 0.742748788 | 0.191070321 | 0.96377671 | 1.14460529 | 0.515184034 | 0.800296691 | 0.822570364 | 0.904460639 | 0.808989691 | 0.646130775 | 0.771147135 | 0.494416236 | 0.190810679 | 0.252037766 | 0.583271809 | 0.544349522 | 1.447068582 | 0.959528529 | 0.985360269 | 2.412497458 | 2.641607616 | CD4_fos_lo |
| 57 | FAM107B | 1.06128214 | 0.852338446 | 0.772531933 | 0.910099063 | 1.574338102 | 1.213384431 | 0.80801735 | 1.190781186 | 0.707778272 | 1.136896652 | 1.432211595 | 1.44010729 | 1.614469419 | 0.89857243 | 1.403347617 | 0.911113866 | 0.498385339 | 0.736793722 | 0.629328119 | 0.437786549 | 0.939153731 | 0.571398802 | CD4_fos_lo |
| 58 | BATF | 0.35027594 | 0.409115666 | 0.285802744 | 0.34925313 | 0.329436214 | 1.50268327 | 0.268726537 | 0.310886978 | 0.320784929 | 0.231256666 | 0.719607777 | 0.568103943 | 0.370039284 | 0.109620543 | 0.126999476 | 0.185984011 | 0.05548467 | 0.223997252 | 0.019980852 | 0.22222596 | 0.225221019 | 0.136375786 | Foxp3_cells |
| 59 | TNFRSF18 | 0.116447691 | 0.106273865 | 0.206900488 | 0.180541818 | 0.380305037 | 1.264116154 | 0.140964925 | 0.093675985 | 0.042997412 | 0.735541933 | 0.654428118 | 0.485945725 | 0.284548933 | 0.034889865 | 0.035485113 | 0.335321147 | 0.272981986 | 0.371804389 | 0.089159725 | 0.05249221 | 0.130809141 | 0.062180315 | Foxp3_cells |
| 60 | TNFRSF4 | 0.068826992 | 0.065441204 | 0.230535623 | 0.226785073 | 0.635698265 | 1.208083572 | 0.241975209 | 0.04571749 | 0.03092991 | 0.150516732 | 0.822494951 | 0.347429593 | 0.169575896 | 0.020947774 | 0.021536463 | 0.23526448 | 0.163901071 | 0.259889066 | 0.110639127 | 0.107244748 | 0.139571468 | 0.131617648 | Foxp3_cells |
| 61 | CTLA4 | 0.117744686 | 0.062047931 | 0.352838989 | 0.137089488 | 0.192063286 | 1.255975796 | 0.132667633 | 0.060969572 | 0.035400567 | 0.058259111 | 0.838591227 | 0.49674056 | 0.239982725 | 0.012783708 | 0.011169788 | 0.11390437 | 0.01490945 | 0.020008394 | 0 | 0.029444732 | 0.034543334 | 0.023897508 | Foxp3_cells |
| 62 | IL2RA | 0.035802084 | 0.017363741 | 0.150603037 | 0.087028257 | 0.288113111 | 0.842907883 | 0.062047232 | 0.163828662 | 0.043432365 | 0.126129966 | 0.142928084 | 0.203874808 | 0.118067662 | 0.067493259 | 0.02070547 | 0.072282616 | 0.006319088 | 0.11963649 | 0 | 0.035148837 | 0.103932526 | 0.208936776 | Foxp3_cells |
| 63 | NAMPT | 0.634728634 | 0.26707633 | 0.683273161 | 0.38917759 | 0.693699174 | 1.498882545 | 0.176302982 | 0.540419546 | 0.285957024 | 0.504580728 | 0.75633649 | 0.774192333 | 0.458025162 | 0.483207873 | 0.440435961 | 0.631968931 | 0.205216537 | 0.608858017 | 0.47957605 | 1.701508507 | 2.506747743 | 1.27795969 | Foxp3_cells |
| 64 | ARID5B | 0.633370749 | 0.392937592 | 0.492180678 | 0.345594007 | 0.619196445 | 1.567873138 | 0.376718726 | 0.268206107 | 0.15147909 | 0.390711945 | 1.167760129 | 0.490919255 | 0.554325319 | 0.569001007 | 0.632005907 | 0.535471881 | 0.280186089 | 0.262832809 | 0.045954417 | 0.183343239 | 0.294562135 | 0.203618886 | Foxp3_cells |
| 65 | RTKN2 | 0.047564717 | 0.071726267 | 0.105563995 | 0.047250922 | 0.044927929 | 0.663816123 | 0.074634738 | 0.015997904 | 0.017285085 | 0.043624206 | 0.143399717 | 0.019300562 | 0.119600652 | 0.009735305 | 0.023231997 | 0.036575012 | 0.004764536 | 0.053466456 | 0 | 0.004746737 | 0.010402355 | 0.018119237 | Foxp3_cells |
| 66 | FOXP3 | 0.009365577 | 0.00389413 | 0.041464177 | 0.019524125 | 0.013586185 | 0.700767217 | 0.022964505 | 0.00748429 | 0.003467774 | 0.00396313 | 0.066719886 | 0.037814971 | 0.041408508 | 0.004443446 | 0.00357709 | 0.036214629 | 0.002685303 | 0.003057805 | 0.017151997 | 0.015225341 | 0.012608898 | 0.014766714 | Foxp3_cells |
| 67 | GADD45A | 0.479282018 | 0.237379627 | 0.483196344 | 0.363556872 | 0.458060479 | 1.096249652 | 0.156371986 | 0.377359877 | 0.274682667 | 0.551840543 | 0.577012946 | 0.558674588 | 0.408092353 | 0.191375846 | 0.236345101 | 0.800890629 | 0.471525103 | 0.397275863 | 0.455445251 | 0.392388231 | 0.371355734 | 0.237672236 | Foxp3_cells |
| 68 | DUSP4 | 0.678409285 | 0.185890279 | 0.295578474 | 0.081861209 | 0.206900335 | 1.101339486 | 0.052361231 | 0.397072059 | 0.176817173 | 0.609994218 | 0.544442088 | 0.551120008 | 0.352612171 | 0.107779577 | 0.083391602 | 0.280068997 | 0.035510258 | 0.148720569 | 0.067975945 | 0.058317389 | 0.431637914 | 0.140505947 | Foxp3_cells |
| 69 | TBC1D4 | 0.116889013 | 0.104694781 | 0.327675092 | 0.184967592 | 0.160648044 | 1.064631607 | 0.347949052 | 0.083589743 | 0.070930008 | 0.082536551 | 0.832899652 | 0.12167287 | 0.157521133 | 0.019288177 | 0.083258598 | 0.081625471 | 0.011842868 | 0.070755688 | 0.279472561 | 0.016169618 | 0.100135935 | 0.09319491 | Foxp3_cells |
| 70 | MIR4435-1HG | 0.185626645 | 0.183034926 | 0.125777992 | 0.068650644 | 0.121895322 | 0.857193633 | 0.082967633 | 0.135127095 | 0.116698295 | 0.159245145 | 0.309610456 | 0.328959333 | 0.326426761 | 0.017404214 | 0.053175484 | 0.268313944 | 0.105565 | 0.185430935 | 0.168633623 | 0.456052756 | 0.511190098 | 0.327381626 | Foxp3_cells |
| 71 | IKZF2 | 0.179319017 | 0.204313218 | 0.156486403 | 0.03575873 | 0.03315075 | 0.779918526 | 0.07374969 | 0.053625805 | 0.055490713 | 0.390208362 | 0.184601227 | 0.185699816 | 0.139487796 | 0.036659618 | 0.053546959 | 0.050063393 | 0.009641538 | 0.144963609 | 0.080959421 | 0.015336274 | 0.021531761 | 0.015547909 | Foxp3_cells |
| 72 | IL1R1 | 0.007233145 | 0.003316489 | 0.09596549 | 0.034437178 | 0.076145217 | 0.63029125 | 0.018908959 | 0.00325479 | 0.001827383 | 0.014173349 | 0.056293265 | 0.154217885 | 0.054458829 | 0.003788572 | 0.004633314 | 0.044336469 | 0.006498038 | 0.315109209 | 0.035364717 | 0.042345992 | 0.20032872 | 0.065896476 | Foxp3_cells |
| 73 | CARD16 | 0.344989105 | 0.489205862 | 0.126168906 | 0.2648018 | 0.260901647 | 0.949452986 | 0.269703424 | 0.225293141 | 0.314796013 | 0.322188387 | 0.361371326 | 0.421711675 | 0.29927005 | 0.209308668 | 0.160630071 | 0.219192661 | 0.15941372 | 0.326864153 | 0.120710208 | 0.41595404 | 0.89931326 | 0.627544324 | Foxp3_cells |
| 74 | TIGIT | 0.647483435 | 0.74285381 | 0.24213105 | 0.12484249 | 0.091797149 | 1.123827268 | 0.224912717 | 0.489068934 | 0.387078007 | 0.572569152 | 0.832130027 | 0.275613274 | 0.21966421 | 0.028979004 | 0.049410331 | 0.14489415 | 0.025812855 | 0.010600809 | 0.053885965 | 0.013715926 | 0.031752051 | 0.023622898 | Foxp3_cells |
| 75 | UGP2 | 0.282658323 | 0.361024616 | 0.447621324 | 0.407176808 | 0.320947452 | 0.996749502 | 0.373791525 | 0.408538971 | 0.488278542 | 0.354295189 | 0.353926701 | 0.318925835 | 0.628984995 | 0.240142828 | 0.415778369 | 0.277595717 | 0.212962864 | 0.372154154 | 0.263877759 | 0.256741926 | 0.375744506 | 0.347101271 | Foxp3_cells |
| 76 | IL32 | 2.126231996 | 2.327119418 | 0.677295752 | 1.750692685 | 1.82503027 | 2.720911019 | 1.618255451 | 1.950345517 | 2.092160393 | 1.732816847 | 1.893197143 | 2.042511534 | 0.756664038 | 0.07188351 | 0.075855226 | 0.529824999 | 0.09354494 | 0.514399363 | 0.030157341 | 0.103213757 | 0.172151849 | 0.167229069 | Foxp3_cells |
| 77 | PELI1 | 0.181663829 | 0.111164355 | 0.424412033 | 0.219528754 | 0.32692992 | 0.88447859 | 0.125341056 | 0.208724316 | 0.112719591 | 0.185265053 | 0.506194712 | 0.35000791 | 0.477019846 | 0.260200472 | 0.516433857 | 0.622271258 | 0.18184037 | 0.201404554 | 0.118951807 | 0.406643663 | 0.604046988 | 0.221238446 | Foxp3_cells |
| 78 | ICA1 | 0.010777387 | 0.027537724 | 0.022988364 | 0.032514889 | 0.023570838 | 0.639417847 | 0.051879211 | 0.013324162 | 0.011519006 | 0.030765121 | 0.58639604 | 0.046268025 | 0.091955293 | 0.011042627 | 0.069183773 | 0.038985128 | 0.006564496 | 0.042270024 | 0 | 0.013190247 | 0.021174573 | 0.030812464 | Foxp3_cells |
| 79 | GK | 0.045544186 | 0.048124152 | 0.102534919 | 0.041953887 | 0.117952282 | 0.597840546 | 0.048333104 | 0.026330017 | 0.011555247 | 0.055795031 | 0.441681514 | 0.115580057 | 0.186086917 | 0.053252282 | 0.110531444 | 0.097197516 | 0.029484146 | 0.133138092 | 0.158001064 | 0.818724177 | 1.209046578 | 0.33291499 | Foxp3_cells |
| 80 | ZC3H12D | 0.155102792 | 0.106641514 | 0.563959373 | 0.29555484 | 0.58535397 | 0.972469891 | 0.238540157 | 0.286107469 | 0.148272223 | 0.065102675 | 0.627491772 | 0.349033363 | 0.279300752 | 0.142736155 | 0.374573158 | 0.150777824 | 0.03320524 | 0.046920135 | 0.029706113 | 0.068693863 | 0.055284653 | 0.091965834 | Foxp3_cells |
| 81 | PMAIP1 | 0.890377904 | 0.378065037 | 0.732294481 | 0.586462695 | 0.777780472 | 1.251363804 | 0.179063018 | 0.897337974 | 0.428974205 | 0.863685487 | 0.537414508 | 0.737921862 | 0.803085699 | 1.046660927 | 0.994334766 | 0.503118187 | 0.142229393 | 0.31927586 | 1.659698457 | 0.897608915 | 0.95266253 | 0.376886785 | Foxp3_cells |
| 82 | GIMAP7 | 0.818628172 | 1.659051211 | 0.47962065 | 1.09657678 | 0.495965001 | 0.421863158 | 1.529123043 | 0.520978812 | 1.406549748 | 0.755222305 | 0.942186426 | 0.771050426 | 0.268601325 | 0.026258332 | 0.027644725 | 0.159413124 | 0.033553591 | 0.240043926 | 0.008048118 | 0.111207481 | 0.158116901 | 0.418018742 | memory_CD4 |
| 83 | GIMAP4 | 0.683754478 | 1.239608355 | 0.341179766 | 0.783311746 | 0.408377097 | 0.294659743 | 1.130223766 | 0.40913253 | 0.897031905 | 0.429470183 | 0.702912799 | 0.547588754 | 0.180457572 | 0.020243855 | 0.021520126 | 0.115418323 | 0.021304752 | 0.144897839 | 0 | 0.12648472 | 0.250269369 | 0.522504166 | memory_CD4 |
| 84 | TXNIP | 1.107627629 | 1.684908691 | 0.972446342 | 1.596800921 | 0.732483407 | 1.012372922 | 1.686767375 | 1.008411924 | 1.811679759 | 1.202156716 | 0.959579788 | 0.894821042 | 0.390174556 | 1.222893492 | 0.321011704 | 1.337361494 | 1.227126519 | 1.519711334 | 0.384377313 | 0.862823148 | 1.030880357 | 1.167023043 | memory_CD4 |
| 85 | KIAA1551 | 0.99843942 | 1.372153073 | 0.990440858 | 0.98866814 | 0.685311437 | 0.943945636 | 1.500927723 | 0.782536817 | 1.070812927 | 0.772161387 | 1.273797443 | 0.856677556 | 0.539619365 | 0.737696985 | 0.558925257 | 0.538385599 | 0.382606115 | 0.371401919 | 0.402289062 | 0.263307013 | 0.385922432 | 0.348409754 | memory_CD4 |
| 86 | ATM | 0.711049707 | 1.063848566 | 0.755233935 | 0.655888422 | 0.524185319 | 0.495343908 | 1.018325834 | 0.433456529 | 0.646540359 | 0.782131808 | 0.724400747 | 0.497276806 | 0.638144909 | 0.759652609 | 1.062194515 | 0.723757604 | 0.653580478 | 0.433295316 | 0.369889994 | 0.32425648 | 0.274663905 | 0.357534888 | memory_CD4 |
| 87 | N4BP2L2 | 1.093473661 | 1.381425677 | 1.314251667 | 1.201047264 | 1.078595194 | 1.322863376 | 1.583997101 | 1.070030129 | 1.262875785 | 1.082855855 | 1.428967186 | 1.217098538 | 0.760048891 | 1.191265467 | 0.929256742 | 1.134665691 | 0.944601991 | 1.229998305 | 1.131109296 | 0.80238437 | 0.720288322 | 0.612190725 | memory_CD4 |
| 88 | TRAF3IP3 | 0.453041173 | 0.747212494 | 0.398300771 | 0.606613451 | 0.428671834 | 0.598964004 | 0.889904867 | 0.443491529 | 0.697067685 | 0.461573166 | 0.682700733 | 0.487690706 | 0.293985959 | 0.417648213 | 0.253867301 | 0.274157602 | 0.227099198 | 0.264248952 | 0.22990316 | 0.105983651 | 0.183123605 | 0.120790207 | memory_CD4 |
| 89 | NKTR | 0.865246046 | 1.076354146 | 1.08895667 | 0.889155474 | 0.721778885 | 0.786506871 | 1.223606414 | 0.725899651 | 0.977022454 | 0.923594629 | 0.964627435 | 0.811315028 | 1.012928906 | 0.762913573 | 1.351641571 | 0.854973624 | 0.682997924 | 0.825133207 | 0.804212436 | 0.484886103 | 0.430190047 | 0.371720896 | memory_CD4 |
| 90 | HIST1H1E | 0.14230433 | 0.272345062 | 0.486967487 | 0.143124409 | 0.130593562 | 0.17998414 | 0.424962948 | 0.13691259 | 0.167650012 | 0.226574764 | 0.225515338 | 0.136424298 | 0.604929861 | 0.23350618 | 0.303324768 | 0.364177834 | 0.284158252 | 0.127723272 | 0.239816507 | 0.618199996 | 0.085105896 | 0.088685415 | memory_CD4 |
| 91 | GIMAP5 | 0.343976549 | 0.622557548 | 0.269880672 | 0.442188538 | 0.215339021 | 0.175622129 | 0.605743972 | 0.250361747 | 0.506800154 | 0.271697215 | 0.335099348 | 0.370293433 | 0.096354808 | 0.012263485 | 0.009865971 | 0.071659979 | 0.014678573 | 0.093082178 | 0 | 0.177843961 | 0.125345386 | 0.232438597 | memory_CD4 |
| 92 | DGKA | 0.157285327 | 0.2048733 | 0.263925833 | 0.294914221 | 0.20681232 | 0.287926954 | 0.486377057 | 0.146598388 | 0.243210293 | 0.0923488 | 0.450344731 | 0.191920113 | 0.126985356 | 0.139208825 | 0.129206906 | 0.107389902 | 0.076911553 | 0.08472329 | 0.085871269 | 0.159923024 | 0.08110253 | 0.049090329 | memory_CD4 |
| 93 | HIST1H1D | 0.148342841 | 0.254688516 | 0.404183066 | 0.218087708 | 0.255879458 | 0.209070697 | 0.436173161 | 0.193012387 | 0.216672793 | 0.140606976 | 0.299125171 | 0.178730283 | 0.866401679 | 0.225997579 | 0.189281361 | 0.459466942 | 0.372604088 | 0.114340281 | 0.405601548 | 0.196447616 | 0.068392438 | 0.069884132 | memory_CD4 |
| 94 | IL2 | 0.080810524 | 0.024482534 | 0.126828093 | 0.265478954 | 0.170781535 | 0.015002794 | 0.085975866 | 0.501530998 | 0.399644535 | 0.099402551 | 0.026044407 | 0.139130376 | 0.050121662 | 0.006886327 | 0.008423441 | 0.048484615 | 0.010356227 | 0.031224799 | 0.006891616 | 0.008469297 | 0.010774395 | 0.0020501 | memory_CD8 |
| 95 | CCL4L2 | 0.683294319 | 0.556446518 | 0.299392323 | 0.191238533 | 0.11042715 | 0.040551206 | 0.144212855 | 0.852257634 | 0.622178204 | 0.510776441 | 0.02281 | 0.386764239 | 0.080123919 | 0.069408863 | 0.063305622 | 0.091175906 | 0.043238906 | 0.137220796 | 0.043217747 | 0.18142013 | 0.296786729 | 0.250193874 | memory_CD8 |
| 96 | SPRY1 | 0.226224852 | 0.049994213 | 0.481670788 | 0.581433036 | 0.49833477 | 0.076962636 | 0.158361051 | 1.543201755 | 0.866936194 | 0.658994265 | 0.17383522 | 1.123681896 | 0.204266686 | 0.125009059 | 0.03014857 | 0.17725428 | 0.037403614 | 0.427649821 | 0.052426634 | 0.043065464 | 0.044769982 | 0.040289381 | memory_CD8 |
| 97 | RASGEF1B | 0.791723117 | 0.190869291 | 0.476427072 | 0.57435034 | 0.647690587 | 0.162693121 | 0.098613647 | 1.225052636 | 0.399396424 | 0.908098678 | 0.359074848 | 0.64264697 | 0.431745407 | 0.922335412 | 0.685806679 | 0.349787836 | 0.074678035 | 0.606925541 | 1.197512067 | 0.619871988 | 1.278193318 | 0.566210037 | memory_CD8 |
| 98 | ANXA1 | 1.907464378 | 1.248008041 | 0.962060638 | 1.914532678 | 2.403291299 | 0.206485574 | 0.853666274 | 2.52726815 | 1.778770787 | 1.284420968 | 0.407444947 | 1.501617102 | 0.395317743 | 0.072460299 | 0.064740138 | 0.44277063 | 0.089484723 | 2.02653872 | 0.098943933 | 0.496685073 | 1.670861481 | 1.092753865 | memory_CD8 |
| 99 | MYC | 0.217696171 | 0.095144641 | 0.236473076 | 0.3193851 | 0.387667303 | 0.094654055 | 0.182468901 | 0.686685203 | 0.396411394 | 0.34185966 | 0.125112801 | 0.244185914 | 0.220795658 | 0.521796736 | 0.211147028 | 0.232680172 | 0.073033941 | 0.112092102 | 0.016709858 | 0.103109325 | 0.109798407 | 0.077233007 | memory_CD8 |
| 100 | CD8B | 0.743213732 | 0.754989159 | 0.066827346 | 0.09112968 | 0.055979137 | 0.017444069 | 0.146313824 | 0.979995521 | 1.180463103 | 0.233859615 | 0.058515366 | 0.220731658 | 0.104357497 | 0.007133791 | 0.008355648 | 0.065849461 | 0.007911142 | 0.015294165 | 0.008018686 | 0.00503038 | 0.01152094 | 0.010840875 | naive_CD8 |
| 101 | CD8A | 1.010172221 | 1.025881244 | 0.124408759 | 0.121862548 | 0.098392733 | 0.04545837 | 0.177100235 | 1.180786192 | 1.257186738 | 0.383724304 | 0.024876219 | 0.695856779 | 0.159757699 | 0.011765816 | 0.013094764 | 0.095589869 | 0.017111257 | 0.032160248 | 0.01848101 | 0.014612985 | 0.023875953 | 0.023913481 | naive_CD8 |
| 102 | ITGA1 | 0.326991976 | 0.216094833 | 0.249133048 | 0.382705342 | 0.241193083 | 0.026290151 | 0.329152735 | 0.883070068 | 0.981807537 | 0.589286067 | 0.022850296 | 0.589133222 | 0.142106297 | 0.011190254 | 0.012835272 | 0.096647049 | 0.012400097 | 0.151239217 | 0.009141119 | 0.02919583 | 0.02724527 | 0.033939267 | naive_CD8 |
| 103 | CISH | 0.051057678 | 0.140500679 | 0.043424554 | 0.122837595 | 0.025792285 | 0.108275641 | 0.133187284 | 0.084673064 | 0.349858652 | 0.069341436 | 0.024164766 | 0.086837177 | 0.024386796 | 0.008713842 | 0.005012187 | 0.026581463 | 0.010920255 | 0.029762372 | 0.082126131 | 0.079707253 | 0.057989372 | 0.051278104 | naive_CD8 |
| 104 | CAPG | 0.150036843 | 0.10221936 | 0.034776597 | 0.1742345 | 0.163305147 | 0.04743888 | 0.121840745 | 0.361956525 | 0.443920533 | 0.452225634 | 0.043909046 | 0.271092223 | 0.16600444 | 0.302956035 | 0.0803973 | 0.096452255 | 0.071891505 | 1.04315834 | 0.744656604 | 0.098788524 | 0.59611096 | 0.728063415 | naive_CD8 |
| 105 | CD3G | 1.161058373 | 1.470848145 | 0.367155216 | 1.214196155 | 1.046052957 | 1.11348647 | 1.258928741 | 1.357844934 | 1.6831422 | 0.594346633 | 1.228267612 | 1.490619981 | 0.457559365 | 0.031757632 | 0.033852087 | 0.297636478 | 0.042515525 | 0.089936447 | 0 | 0.023775691 | 0.066710629 | 0.052891807 | naive_CD8 |
| 106 | CKLF | 0.667194284 | 0.748521463 | 0.188940067 | 0.738297426 | 0.669721181 | 0.73194281 | 0.588882047 | 1.00436838 | 1.207708012 | 0.87966274 | 0.678412769 | 1.140342268 | 0.61359239 | 0.184779796 | 0.176448253 | 0.41326252 | 0.281828272 | 0.725435921 | 0.321517182 | 0.071734927 | 0.504594959 | 0.359406403 | naive_CD8 |
| 107 | GNLY | 0.547880149 | 0.892283602 | 0.093175789 | 0.038134896 | 0.074297522 | 0.081609728 | 0.046834085 | 0.305683625 | 0.18594766 | 2.320167538 | 0.038866727 | 0.470620541 | 0.236078109 | 0.009012457 | 0.005049909 | 0.063729958 | 0.022023571 | 0.169278701 | 0.104341705 | 0.023687577 | 0.037403852 | 0.048316759 | NK_cells |
| 108 | XCL1 | 0.339198279 | 0.16684207 | 0.081489196 | 0.045298125 | 0.020728856 | 0.01412557 | 0.026485611 | 0.998000915 | 0.490085715 | 1.857147448 | 0.008106504 | 0.042992011 | 0.07448805 | 0.005711205 | 0.005790067 | 0.042724995 | 0.010885081 | 0.096847036 | 0 | 0.002758645 | 0.013834213 | 0.0102451 | NK_cells |
| 109 | XCL2 | 0.806073158 | 0.532637149 | 0.106951373 | 0.037319005 | 0.014087404 | 0.015924737 | 0.055982516 | 1.232394037 | 0.721383992 | 2.095211704 | 0.007878498 | 0.058042665 | 0.09889789 | 0.009441753 | 0.004365757 | 0.06741977 | 0.015877164 | 0.074493106 | 0 | 0.010374144 | 0.022489675 | 0.014289119 | NK_cells |
| 110 | TYROBP | 0.091117626 | 0.190373275 | 0.028567768 | 0.009938151 | 0.008054956 | 0.010073321 | 0.012231052 | 0.021858968 | 0.021991965 | 1.257354415 | 0.005209745 | 0.024707575 | 0.05691212 | 0.012757013 | 0.012459735 | 0.029769255 | 0.030749946 | 1.087137991 | 1.759870068 | 0.927302785 | 2.397956789 | 2.695096386 | NK_cells |
| 111 | FCER1G | 0.044444543 | 0.097848872 | 0.017597478 | 0.005682668 | 0.006481718 | 0.008961961 | 0.007319395 | 0.019837922 | 0.020992718 | 1.109995665 | 0.004398825 | 0.008281059 | 0.051243277 | 0.006391278 | 0.003559875 | 0.01102425 | 0.005580325 | 1.197844691 | 1.479354852 | 0.489665362 | 2.06025025 | 2.123203774 | NK_cells |
| 112 | KLRC1 | 0.160016501 | 0.144988591 | 0.073131462 | 0.016487316 | 0.007495087 | 0.0036024 | 0.03435862 | 0.36226742 | 0.377657434 | 1.029844883 | 0.00221873 | 0.10717236 | 0.056833985 | 0.001551375 | 0.001465296 | 0.022385459 | 0.003969889 | 0.106880145 | 0 | 0 | 0.007640487 | 0.00998188 | NK_cells |
| 113 | KLRD1 | 0.753302907 | 0.746523334 | 0.137817987 | 0.047140173 | 0.032718842 | 0.015509146 | 0.08314203 | 0.862042254 | 0.720752605 | 1.558714964 | 0.014994493 | 0.28253348 | 0.121775944 | 0.007806063 | 0.003077223 | 0.06592771 | 0.009109094 | 0.042458 | 0 | 0.007915933 | 0.019910411 | 0.021009644 | NK_cells |
| 114 | HOPX | 0.516070195 | 0.546972926 | 0.050862362 | 0.236543702 | 0.277128294 | 0.025546837 | 0.146569033 | 0.725840132 | 0.720403105 | 1.448196205 | 0.048507651 | 0.682595803 | 0.448005064 | 0.129960943 | 0.659640707 | 0.095793407 | 0.021083334 | 0.115856027 | 0 | 0.017059249 | 0.033217395 | 0.03275059 | NK_cells |
| 115 | CD160 | 0.297999082 | 0.252027115 | 0.037507525 | 0.038591384 | 0.013933492 | 0.00924151 | 0.039498659 | 0.529375995 | 0.597101578 | 0.916407471 | 0.009989376 | 0.160378779 | 0.045018422 | 0.005142552 | 0.004656557 | 0.033567979 | 0.008625568 | 0.009447069 | 0.008306775 | 0 | 0.007007107 | 0.005709236 | NK_cells |
| 116 | KLRF1 | 0.083500704 | 0.171191119 | 0.001971595 | 0.002751987 | 0.001281381 | 0.001541826 | 0.003683047 | 0.008252665 | 0.007619033 | 0.584021889 | 0.001447608 | 0.023914424 | 0.015396472 | 0.004119457 | 0.001169076 | 0.006885271 | 0.006151396 | 0.005461431 | 0.009777078 | 0 | 0.004465154 | 0 | NK_cells |
| 117 | KLRC2 | 0.099472906 | 0.07987325 | 0.0183702 | 0.007626407 | 0.006087845 | 0.003708492 | 0.03087452 | 0.271170505 | 0.251044207 | 0.652301728 | 0.000705675 | 0.017536218 | 0.041612613 | 0.000780799 | 0.000629746 | 0.010884096 | 0.002721532 | 0.016616831 | 0 | 0 | 0.003275475 | 0 | NK_cells |
| 118 | CTSW | 0.734700882 | 1.013120687 | 0.067469345 | 0.107770545 | 0.097877896 | 0.017507613 | 0.089791619 | 0.406367891 | 0.496445201 | 1.160011285 | 0.01560765 | 0.318444768 | 0.111912032 | 0.013983429 | 0.00920378 | 0.074587857 | 0.044591729 | 0.296278705 | 0.027572309 | 0.015320368 | 0.022194933 | 0.017775592 | NK_cells |
| 119 | IL2RB | 0.408151087 | 0.445017902 | 0.084398006 | 0.179244761 | 0.238216221 | 0.519209222 | 0.150534022 | 0.382342347 | 0.299688542 | 1.140647843 | 0.285783993 | 0.394738796 | 0.175380168 | 0.010639896 | 0.009034644 | 0.085105668 | 0.011229895 | 0.092590261 | 0.013083281 | 0.014904637 | 0.026553148 | 0.025663583 | NK_cells |
| 120 | GSTP1 | 0.576746384 | 0.699181752 | 0.108628786 | 0.406233578 | 0.391519006 | 0.28025055 | 0.35933875 | 0.570789244 | 0.694370155 | 1.238543235 | 0.251135835 | 0.470451519 | 1.42746493 | 0.462114071 | 1.161682085 | 1.094411519 | 1.289153201 | 1.057372508 | 1.339045403 | 0.25004768 | 1.425871935 | 1.749353281 | NK_cells |
| 121 | CCL3 | 1.08453226 | 0.809983579 | 0.229373926 | 0.086871567 | 0.059700838 | 0.06663628 | 0.058519187 | 0.326406792 | 0.227361915 | 1.351376863 | 0.054342939 | 0.487269279 | 0.338013653 | 0.243079654 | 0.19944241 | 0.39414509 | 0.169591055 | 0.117431694 | 0.81863979 | 1.639270473 | 2.477538537 | 1.775617417 | NK_cells |
| 122 | MAF | 0.322876074 | 0.397214067 | 0.675795526 | 0.429424496 | 0.525062489 | 1.395613965 | 0.536864599 | 0.139153771 | 0.162306527 | 0.123845031 | 1.478747476 | 0.983769544 | 0.273027717 | 0.025771519 | 0.030938316 | 0.219985137 | 0.031139593 | 0.322635691 | 0.035353569 | 0.475680252 | 0.304114208 | 0.594978943 | Tfh_cells |
| 123 | TOX2 | 0.038761028 | 0.043405989 | 0.053190892 | 0.023314653 | 0.020235952 | 0.25549533 | 0.064657076 | 0.014618391 | 0.014191691 | 0.067361989 | 0.545251046 | 0.058876415 | 0.092581748 | 0.006413014 | 0.010461961 | 0.050159001 | 0.017906391 | 0.100857921 | 0.195803956 | 0.003444113 | 0.013064245 | 0.010885702 | Tfh_cells |
| 124 | PASK | 0.121196094 | 0.113869647 | 0.09478526 | 0.21243546 | 0.230255213 | 0.207282505 | 0.274785391 | 0.120053206 | 0.111593389 | 0.037835852 | 0.680969613 | 0.101982984 | 0.134741097 | 0.042375311 | 0.210369163 | 0.065646652 | 0.044378747 | 0.033865597 | 0.00728113 | 0.014104641 | 0.021782399 | 0.034015785 | Tfh_cells |
| 125 | RNF19A | 1.309390141 | 0.838454142 | 0.94474483 | 0.616673985 | 1.006350284 | 1.143996882 | 0.595815512 | 0.884866923 | 0.452667377 | 0.87898933 | 1.739691545 | 1.397204386 | 0.704163193 | 0.618219427 | 0.637418675 | 0.538728402 | 0.287664891 | 0.451179205 | 0.412047157 | 0.428136873 | 0.331857631 | 0.265173947 | Tfh_cells |
| 126 | TSHZ2 | 0.017656659 | 0.026733475 | 0.17090607 | 0.1155474 | 0.159417458 | 0.156567054 | 0.283813711 | 0.004820613 | 0.003988852 | 0.006245109 | 0.554588065 | 0.021828754 | 0.05654807 | 0.011747851 | 0.012822102 | 0.093379328 | 0.070163862 | 0.100840286 | 0 | 0.010063166 | 0.005035089 | 0.002366069 | Tfh_cells |
| 127 | FKBP5 | 0.188606167 | 0.170353882 | 0.167963019 | 0.358908564 | 0.317287278 | 0.226972059 | 0.45438906 | 0.212310221 | 0.235930121 | 0.149681378 | 0.80129649 | 0.343244938 | 0.428223776 | 0.074982432 | 0.189400972 | 0.174020984 | 0.075613058 | 0.154661468 | 0.424934217 | 0.181798969 | 0.379988327 | 0.299928918 | Tfh_cells |
| 128 | CORO1B | 0.197845509 | 0.263919932 | 0.115057192 | 0.346147907 | 0.361692769 | 0.72595306 | 0.371513172 | 0.218641867 | 0.284059476 | 0.151129037 | 0.853932617 | 0.247818441 | 0.353297673 | 0.082262651 | 0.175227291 | 0.235528241 | 0.194665259 | 0.204744576 | 0.231021206 | 0.067907487 | 0.345205468 | 0.512212514 | Tfh_cells |
| 129 | ITM2A | 0.607959365 | 0.598292765 | 0.180697222 | 0.553495456 | 0.736920014 | 0.608505568 | 0.476804002 | 0.652064804 | 0.507521646 | 0.553584525 | 1.15590532 | 0.522474599 | 0.313986198 | 0.062589341 | 0.01776335 | 0.2837478 | 0.161938729 | 0.43801078 | 0.026486936 | 0.021426244 | 0.033714203 | 0.040709483 | Tfh_cells |
| 130 | SESN3 | 0.065089989 | 0.09222335 | 0.110495306 | 0.131948468 | 0.153820684 | 0.343448141 | 0.225775044 | 0.087589803 | 0.075584296 | 0.066899323 | 0.576733266 | 0.11373395 | 0.200386799 | 0.433057681 | 0.346662285 | 0.065482245 | 0.013883842 | 0.105816614 | 0.072416591 | 0.104665791 | 0.090939614 | 0.10187495 | Tfh_cells |
| 131 | IL6ST | 0.189961481 | 0.09336761 | 0.322826179 | 0.246761427 | 0.332945405 | 0.645280401 | 0.32667029 | 0.211253283 | 0.099268132 | 0.184321364 | 0.790353673 | 0.309877925 | 0.147614112 | 0.123390568 | 0.065300555 | 0.273452151 | 0.212414488 | 0.395821121 | 0.297558446 | 0.142977484 | 0.360781045 | 0.219020101 | Tfh_cells |
| 132 | RP5-1028K7.2 | 0.078763552 | 0.150719826 | 0.016080527 | 0.032352636 | 0.035061315 | 0.01437975 | 0.067386933 | 0.004952867 | 0.009330116 | 0.023773874 | 0.37523973 | 0.019691235 | 0.037429803 | 0.04978368 | 0.011820697 | 0.01180289 | 0.003479168 | 0.013388973 | 0 | 0.003122497 | 0.005086569 | 0.003356255 | Tfh_cells |
| 133 | TRPS1 | 0.368428817 | 0.352648332 | 0.473390546 | 0.196990827 | 0.186527913 | 0.633072129 | 0.274761301 | 0.164322897 | 0.171049057 | 0.142540363 | 0.779396378 | 0.271977181 | 0.217457002 | 0.117346884 | 0.112454599 | 0.125775451 | 0.030269654 | 0.205927751 | 0.139639491 | 0.461237806 | 0.419628305 | 0.215740006 | Tfh_cells |
| 134 | AHI1 | 0.227149951 | 0.155659711 | 0.198250123 | 0.136529821 | 0.192069931 | 0.212955839 | 0.185594152 | 0.184708355 | 0.144520504 | 0.428757822 | 0.57877395 | 0.20821834 | 0.309280105 | 0.193632253 | 0.128085802 | 0.330951693 | 0.263169031 | 0.503811523 | 0.717393939 | 0.145694002 | 0.133597933 | 0.085051815 | Tfh_cells |
| 135 | LIMS1 | 0.333074185 | 0.288939 | 0.322451716 | 0.519244853 | 0.624297968 | 0.916586839 | 0.453514792 | 0.476453783 | 0.378907917 | 0.536118653 | 1.129202773 | 0.825761257 | 0.615083875 | 0.14152001 | 0.539991579 | 0.325652311 | 0.113826926 | 0.519506384 | 0.186536599 | 0.708499311 | 1.321226553 | 0.95532694 | Tfh_cells |
| 136 | SLC9A9 | 0.092314096 | 0.170752082 | 0.165046411 | 0.097765857 | 0.064984572 | 0.21644615 | 0.251048028 | 0.045398677 | 0.068495653 | 0.074232321 | 0.480136562 | 0.0766492 | 0.061800171 | 0.072722786 | 0.063655336 | 0.064380971 | 0.03671092 | 0.071272955 | 0.008107145 | 0.168046269 | 0.089010262 | 0.148866504 | Tfh_cells |
| 137 | PDCD1 | 0.11499002 | 0.132113389 | 0.044382571 | 0.064605332 | 0.087508437 | 0.12628627 | 0.062540138 | 0.028913424 | 0.026438832 | 0.024631631 | 0.417052643 | 0.231552354 | 0.087022068 | 0.008325717 | 0.010577626 | 0.033000473 | 0.004071023 | 0.007323704 | 0 | 0.01177177 | 0.010402622 | 0.008891479 | Tfh_cells |
| 138 | THADA | 0.091567684 | 0.09370823 | 0.169186571 | 0.07861397 | 0.083068043 | 0.255771708 | 0.155769733 | 0.084183136 | 0.082123222 | 0.094105895 | 0.432042678 | 0.097062366 | 0.148935037 | 0.095475973 | 0.134661683 | 0.104604491 | 0.075850479 | 0.079376921 | 0.226930785 | 0.054448973 | 0.080213021 | 0.083023664 | Tfh_cells |
| 139 | SMCO4 | 0.019808804 | 0.027021966 | 0.03554019 | 0.082230027 | 0.082454512 | 0.176614364 | 0.077665108 | 0.048850211 | 0.049802601 | 0.072227452 | 0.412136203 | 0.242871538 | 0.213340823 | 0.025085234 | 0.137578409 | 0.09485828 | 0.067675242 | 0.055272523 | 0.29171013 | 0.138226708 | 0.429784894 | 0.425176063 | Tfh_cells |
| 140 | IL6R | 0.011632183 | 0.015906451 | 0.083829641 | 0.076061305 | 0.092223102 | 0.296840186 | 0.183804986 | 0.003449315 | 0.003644443 | 0.012242467 | 0.409371031 | 0.035608167 | 0.06560271 | 0.014116831 | 0.016193437 | 0.11733934 | 0.097564599 | 0.0937905 | 0.222504055 | 0.280797364 | 0.648050959 | 0.291605654 | Tfh_cells |
| 141 | ST8SIA1 | 0.097255771 | 0.153883749 | 0.091171959 | 0.095406896 | 0.04325824 | 0.056076593 | 0.187109626 | 0.034289586 | 0.064571451 | 0.02716351 | 0.321391513 | 0.030957936 | 0.025120742 | 0.003421677 | 0.004531667 | 0.016716657 | 0.004473595 | 0.061796017 | 0 | 0.008398286 | 0.013432175 | 0.013060682 | Tfh_cells |
| 142 | TOX | 0.196068087 | 0.219885012 | 0.110954232 | 0.065268155 | 0.052271587 | 0.281214455 | 0.107136008 | 0.077840186 | 0.067270531 | 0.248023649 | 0.449137248 | 0.182580609 | 0.250423092 | 0.047384492 | 0.425948505 | 0.073653064 | 0.030329149 | 0.121922053 | 0.046503895 | 0.00748758 | 0.019532043 | 0.019826938 | Tfh_cells |

| | A | B | C | D | E | F | G | H | I | J | K | L | M | N | O | P | Q | R | S | T | U | V | W | X |
|---|---|---|---|---|---|---|---|---|---|---|---|---|---|---|---|---|---|---|---|---|---|---|---|---|
| 143 | CD4 | 0.023085217 | 0.034650127 | 0.060522237 | 0.21284817 | 0.202184182 | 0.403995623 | 0.264872735 | 0.011285851 | 0.027258234 | 0.006896286 | 0.491893025 | 0.185468447 | 0.109185727 | 0.006841698 | 0.013021062 | 0.054670348 | 0.006802868 | 0.060861881 | 0.99635259 | 0.068365683 | 0.412778868 | 0.504838969 | Tfh_cells |
| 144 | PGM2L1 | 0.074255666 | 0.088076264 | 0.146582131 | 0.093773608 | 0.096594088 | 0.165980074 | 0.138447716 | 0.074943954 | 0.085717637 | 0.111060484 | 0.44261944 | 0.093171038 | 0.07785807 | 0.09883288 | 0.028109193 | 0.053563333 | 0.027293913 | 0.146208952 | 0.120039303 | 0.007702621 | 0.025859785 | 0.046030408 | Tfh_cells |
| 145 | ETV6 | 0.144090125 | 0.165514219 | 0.212507207 | 0.209974227 | 0.242175682 | 0.292954443 | 0.256078259 | 0.133702244 | 0.128090142 | 0.177255298 | 0.542625896 | 0.227862754 | 0.153750829 | 0.285867567 | 0.104704808 | 0.265032178 | 0.181452403 | 0.343807837 | 0.924443533 | 0.655941834 | 0.741534797 | 0.317434991 | Tfh_cells |
| 146 | MAGEH1 | 0.067037833 | 0.075416284 | 0.070406042 | 0.081453716 | 0.055190614 | 0.385426928 | 0.099755424 | 0.06164204 | 0.086013012 | 0.067872634 | 0.397920938 | 0.049973481 | 0.081887901 | 0.03867832 | 0.054604826 | 0.164734868 | 0.161757241 | 0.078259776 | 0.037240044 | 0.022162429 | 0.026151968 | 0.027241252 | Tfh_cells |
| 147 | NR3C1 | 0.601804623 | 0.455609586 | 0.615844679 | 0.61213953 | 0.670412576 | 0.73704007 | 0.589979205 | 0.80188047 | 0.586291971 | 0.731985472 | 1.127887738 | 0.745063625 | 0.666887462 | 0.431896887 | 0.849627982 | 0.455393381 | 0.261630593 | 0.650112452 | 1.265218026 | 0.471008752 | 0.56772213 | 0.334028746 | Tfh_cells |
| 148 | PPP1CC | 0.343655613 | 0.348448577 | 0.179562069 | 0.344399627 | 0.360600764 | 0.431902916 | 0.346138396 | 0.335249647 | 0.325203942 | 0.357426265 | 0.716266769 | 0.406799474 | 0.900480961 | 0.447896361 | 0.900532348 | 0.511255801 | 0.41469864 | 0.409681472 | 0.323507316 | 0.071467303 | 0.339248351 | 0.406873857 | Tfh_cells |
| 149 | CD40LG | 0.055792193 | 0.059594293 | 0.250172758 | 0.617390701 | 0.467798366 | 0.107246702 | 0.465091291 | 0.037917421 | 0.065247997 | 0.018254358 | 0.644734933 | 0.476565914 | 0.099535367 | 0.0077301 | 0.008198464 | 0.093865881 | 0.012939617 | 0.044373064 | 0.008085853 | 0.018444327 | 0.01639659 | 0.020001449 | Tfh_cells |
| 150 | C9orf16 | 0.649586134 | 0.738982379 | 0.175740461 | 0.539841831 | 0.660782695 | 0.991545936 | 0.573382184 | 0.585077306 | 0.669070173 | 0.846865539 | 1.082026506 | 0.887216505 | 0.805761082 | 0.488076815 | 0.693035335 | 0.605090913 | 0.564857905 | 0.680462037 | 0.618794328 | 0.137171956 | 0.460849839 | 0.542298517 | Tfh_cells |
| 151 | FYB | 0.952142183 | 0.963511118 | 0.599278235 | 1.006903812 | 0.948172338 | 1.138228623 | 1.174043111 | 1.003493911 | 1.049795248 | 0.774541157 | 1.404198316 | 0.913786213 | 0.378218018 | 0.047471945 | 0.040040651 | 0.313520601 | 0.042512123 | 0.242172707 | 0.540960747 | 0.663553292 | 0.876807398 | 0.771901335 | Tfh_cells |
| 152 | TCF7 | 0.424841333 | 0.360944705 | 0.29830207 | 0.607551429 | 0.570470869 | 0.18810288 | 0.682123903 | 0.465112725 | 0.516004201 | 0.521460951 | 0.767454692 | 0.255211363 | 0.159016965 | 0.113424267 | 0.105628103 | 0.170084609 | 0.035544961 | 0.282292819 | 0.095352767 | 0.027958212 | 0.047280061 | 0.038668763 | Tfh_cells |
| 153 | IL17A | 0.012306682 | 0.008026742 | 0.171409858 | 0.137613219 | 0.139770159 | 0.20896959 | 0.020541415 | 0.038921458 | 0.024556213 | 0.009197389 | 0.169428871 | 0.695792877 | 0.124407931 | 0.006858066 | 0.012880679 | 0.099374199 | 0.010690843 | 0.024671545 | 0.010781018 | 0.02562262 | 0.009485078 | 0.003927656 | Th17_cells |
| 154 | CCL20 | 0.12605243 | 0.059946987 | 0.439003488 | 0.827837586 | 1.2514905 | 0.907032576 | 0.168540501 | 0.34866609 | 0.178102026 | 0.081556287 | 0.363756406 | 2.043933249 | 0.348301519 | 0.050350893 | 0.035704773 | 0.310419422 | 0.054753011 | 0.222530705 | 0.014418875 | 0.157589452 | 0.509372588 | 0.283304369 | Th17_cells |
| 155 | CEBPD | 0.180504731 | 0.12547679 | 0.156015977 | 0.15630241 | 0.314061521 | 0.055108766 | 0.05050031 | 0.113683575 | 0.069471749 | 0.529648349 | 0.036918566 | 0.780713461 | 0.138911899 | 0.012994898 | 0.028915537 | 0.100358854 | 0.028388704 | 0.374462139 | 0.035306385 | 0.649265369 | 1.133684612 | 0.901783874 | Th17_cells |
| 156 | PRR5 | 0.131349106 | 0.176910648 | 0.076902115 | 0.241129709 | 0.2648897 | 0.154930181 | 0.125583765 | 0.198756351 | 0.190593955 | 0.173785469 | 0.098227489 | 0.803135814 | 0.097811237 | 0.032615838 | 0.013365654 | 0.128078366 | 0.074412632 | 0.213828582 | 0.018396174 | 0.025884811 | 0.021080995 | 0.038834696 | Th17_cells |
| 157 | AC092580.4 | 0.337256738 | 0.340239337 | 0.151872814 | 0.278488444 | 0.324274349 | 0.17903413 | 0.211915942 | 0.50423403 | 0.50938357 | 0.5658093 | 0.293303394 | 0.902095958 | 0.256800261 | 0.013131876 | 0.014937723 | 0.119760692 | 0.019748714 | 0.080750123 | 0.052830191 | 0.052770397 | 0.026078285 | 0.02377674 | Th17_cells |
| 158 | MIR155HG | 0.289676122 | 0.156851419 | 0.223590154 | 0.316723274 | 0.361524555 | 0.426415869 | 0.092067273 | 0.200760808 | 0.086166261 | 0.18962328 | 0.282064323 | 0.64716775 | 0.324339568 | 0.266191721 | 0.269411309 | 0.360926212 | 0.112275542 | 0.112225682 | 0.195501989 | 0.248889379 | 0.550922518 | 0.139301028 | Th17_cells |
| 159 | HMGB2 | 0.610030271 | 0.536828645 | 0.402414864 | 0.460998252 | 0.470221852 | 0.611709083 | 0.434033774 | 0.590710495 | 0.519732441 | 0.633328763 | 0.565727288 | 0.57992006 | 2.77605767 | 0.255914085 | 0.729114705 | 0.40570423 | 0.193428824 | 0.44425105 | 0.236086563 | 0.349509775 | 0.35914852 | 0.331901888 | cycling_b_cells |
| 160 | STMN1 | 0.173324032 | 0.119917968 | 0.066289323 | 0.093176996 | 0.118569445 | 0.127334039 | 0.064303404 | 0.130618937 | 0.090126856 | 0.259903159 | 0.095354638 | 0.123154374 | 2.349587078 | 0.146689925 | 0.557644051 | 0.067713092 | 0.034783417 | 0.601928205 | 1.466033799 | 0.076023434 | 0.070296838 | 0.217769734 | cycling_b_cells |
| 161 | HMGN2 | 0.832666229 | 0.893562795 | 0.438894654 | 0.840968173 | 0.754854118 | 1.036530679 | 0.865826234 | 0.845201762 | 0.996529943 | 0.950087121 | 1.021577043 | 0.932893367 | 2.970536383 | 1.072092994 | 1.754970663 | 0.830298871 | 0.629362664 | 1.039018676 | 1.278313074 | 0.691584975 | 0.959744396 | 1.159211753 | cycling_b_cells |
| 162 | TUBA1B | 0.862653636 | 0.676490522 | 0.820642227 | 0.77994598 | 0.83115586 | 0.795904091 | 0.553710643 | 0.795866313 | 0.786603979 | 0.920394904 | 0.723007864 | 0.804790633 | 2.600631415 | 0.814523548 | 0.706753593 | 0.698840662 | 0.472039634 | 1.080392327 | 1.362228724 | 0.469741872 | 1.115305473 | 1.536576921 | cycling_b_cells |
| 163 | H2AFZ | 1.154606482 | 0.86373064 | 0.663923014 | 0.896933056 | 1.179372482 | 1.229749278 | 0.707725328 | 1.122452717 | 0.929205504 | 1.175942471 | 1.229900105 | 1.134733857 | 2.694540765 | 0.715014761 | 1.206705978 | 0.957323232 | 0.785102669 | 0.963316818 | 0.784878323 | 1.154528718 | 1.014397359 | 0.935059524 | cycling_b_cells |
| 164 | TUBB | 0.352683603 | 0.411378241 | 0.143495375 | 0.316298849 | 0.383983312 | 0.424649534 | 0.309564208 | 0.37794179 | 0.422222743 | 0.443567999 | 0.365721867 | 0.408442012 | 2.026846205 | 0.395497091 | 0.57423779 | 0.249831105 | 0.181689166 | 0.523092684 | 0.734517353 | 0.277124202 | 0.599135434 | 0.782519299 | cycling_b_cells |
| 165 | CENPF | 0.123949698 | 0.03923277 | 0.077062227 | 0.02503837 | 0.041750274 | 0.057515353 | 0.032681752 | 0.058160269 | 0.030011388 | 0.109200983 | 0.073750641 | 0.072886192 | 1.196852973 | 0.019427113 | 0.110829219 | 0.034706058 | 0.007968672 | 0.033429854 | 0.024748805 | 0.016724634 | 0.025267479 | 0.028370058 | cycling_b_cells |
| 166 | MKI67 | 0.015506699 | 0.011036454 | 0.038606431 | 0.002390478 | 0.003076818 | 0.008736112 | 0.007284504 | 0.005737899 | 0.006153828 | 0.034705113 | 0.007008365 | 0.015640573 | 1.321350652 | 0.003183941 | 0.093886197 | 0.023845099 | 0.008028175 | 0.005318642 | 0 | 0 | 0.009282716 | 0.022449144 | cycling_b_cells |
| 167 | HMGB1 | 1.663389026 | 1.904238722 | 1.586431626 | 1.630524074 | 1.468320725 | 1.513156517 | 1.725300766 | 1.573010334 | 1.824304341 | 1.762843386 | 1.751849987 | 1.697453979 | 3.418874482 | 1.682093211 | 2.433919973 | 1.776024449 | 1.489606325 | 1.79603545 | 2.113600164 | 1.12143958 | 1.369484542 | 1.500664969 | cycling_b_cells |
| 168 | KIAA0101 | 0.0081379 | 0.013352146 | 0.013107948 | 0.002808118 | 0.005240998 | 0.012790243 | 0.009665973 | 0.006378623 | 0.002745046 | 0.050253872 | 0.011583634 | 0.015582678 | 1.324217035 | 0.007751241 | 0.264473907 | 0.013525267 | 0.007497878 | 0.019015947 | 0.009924031 | 0.004064041 | 0.00773166 | 0.048269326 | cycling_b_cells |
| 169 | DUT | 0.30858824 | 0.382169386 | 0.171133472 | 0.38816942 | 0.378681891 | 0.30980209 | 0.39338319 | 0.381505803 | 0.434361794 | 0.39549086 | 0.396835026 | 0.400896957 | 1.734930578 | 0.338257491 | 0.490082271 | 0.283369175 | 0.208972421 | 0.43381118 | 0.443148871 | 0.118694691 | 0.277501175 | 0.377518807 | cycling_b_cells |
| 170 | GAPDH | 1.669498866 | 1.667923266 | 0.470345537 | 1.325870769 | 1.587363334 | 1.852393622 | 1.200717838 | 1.692222357 | 1.742387523 | 1.96237626 | 1.773286517 | 1.905900953 | 3.272892884 | 1.22672533 | 2.824762838 | 1.390421973 | 1.264997149 | 1.675661074 | 1.256805144 | 0.743052105 | 2.433533063 | 2.318987832 | cycling_b_cells |
| 171 | PTTG1 | 0.257673509 | 0.209119844 | 0.109845151 | 0.124939806 | 0.140064467 | 0.324353874 | 0.08038295 | 0.219054437 | 0.178415213 | 0.197796608 | 0.187725864 | 0.212390703 | 1.37053929 | 0.078429506 | 0.584173737 | 0.11556963 | 0.052500966 | 0.085833898 | 0.142331133 | 0.064504356 | 0.135482353 | 0.150944681 | cycling_b_cells |
| 172 | TOP2A | 0.023989609 | 0.015642455 | 0.014507018 | 0.004119142 | 0.010545702 | 0.016088792 | 0.005412065 | 0.010823058 | 0.00640003 | 0.014652166 | 0.021816648 | 0.019589354 | 1.098760393 | 0.008287121 | 0.053359735 | 0.012155549 | 0.004706901 | 0.004559007 | 0.008233163 | 0.004960273 | 0.012054601 | 0.028628825 | cycling_b_cells |
| 173 | DEK | 0.629705905 | 0.731101832 | 0.287680167 | 0.581599865 | 0.680673918 | 0.59369722 | 0.66026991 | 0.743002329 | 0.712264099 | 0.836572024 | 0.76753354 | 0.891904444 | 1.945341624 | 0.559469613 | 0.958935261 | 0.197171107 | 0.041912815 | 0.687638942 | 0.687408106 | 0.364779821 | 0.861414529 | 0.733999348 | cycling_b_cells |
| 174 | NUSAP1 | 0.132145132 | 0.037219668 | 0.116372704 | 0.033956993 | 0.082829259 | 0.155698781 | 0.016109507 | 0.099359846 | 0.026368095 | 0.063880509 | 0.082726646 | 0.151565046 | 1.222266798 | 0.041169583 | 0.228574566 | 0.065201748 | 0.020175903 | 0.036964361 | 0.091844795 | 0.097087656 | 0.062212724 | 0.067681604 | cycling_b_cells |
| 175 | SMC4 | 0.345489874 | 0.214883173 | 0.248511245 | 0.166488709 | 0.226974445 | 0.4511611 | 0.16048038 | 0.24546393 | 0.146121366 | 0.403558936 | 0.323860528 | 0.352023377 | 1.487024607 | 0.165101374 | 0.414267305 | 0.313715866 | 0.213295287 | 0.325964623 | 0.334727525 | 0.096809502 | 0.196258156 | 0.149451921 | cycling_b_cells |
| 176 | ASPM | 0.013459127 | 0.014496861 | 0.012978201 | 0.003191006 | 0.006912841 | 0.012882012 | 0.006860049 | 0.00464785 | 0.002336376 | 0.01653902 | 0.006550439 | 0.010250597 | 0.917272937 | 0.003453991 | 0.038477494 | 0.023257011 | 0.008107491 | 0.010594839 | 0.037003188 | 0.014836005 | 0.006314037 | 0.003134309 | cycling_b_cells |
| 177 | H2AFV | 0.632758127 | 0.574833806 | 0.266572034 | 0.60680777 | 0.790629703 | 0.741091607 | 0.466279739 | 0.703367306 | 0.575815453 | 0.697027928 | 0.749876974 | 0.871343221 | 1.914105288 | 0.409887269 | 0.947639541 | 0.605258673 | 0.508770522 | 0.671013704 | 0.964776105 | 0.137427862 | 0.625057566 | 0.61514019 | cycling_b_cells |
| 178 | TYMS | 0.008314309 | 0.004796524 | 0.006291616 | 0.003770577 | 0.004113202 | 0.010191636 | 0.004745893 | 0.004695401 | 0.00232619 | 0.052208174 | 0.008554865 | 0.016452224 | 1.062425769 | 0.003404823 | 0.039579829 | 0.015912462 | 0.00926373 | 0.018505394 | 0.007991449 | 0.006714458 | 0.00648348 | 0.035102377 | cycling_b_cells |
| 179 | HIST1H4C | 1.009931478 | 0.757027834 | 1.129683274 | 0.93942451 | 1.149331475 | 0.970781472 | 0.71102147 | 1.145984128 | 0.920117259 | 1.006022747 | 1.018937279 | 0.91198595 | 2.505891621 | 0.932774718 | 1.059982507 | 1.221143815 | 0.791899394 | 0.876301464 | 0.828145903 | 1.126939659 | 0.390850497 | 0.43181888 | cycling_b_cells |
| 180 | CD83 | 0.390595161 | 0.172637606 | 0.591240366 | 0.478559134 | 0.473391742 | 0.462063058 | 0.113758846 | 0.474521975 | 0.291415133 | 0.499291322 | 0.300627449 | 0.622923091 | 0.732444405 | 2.312874518 | 1.574757113 | 0.361496095 | 0.091737958 | 0.868389278 | 2.087258715 | 1.357508788 | 2.200637844 | 1.103553714 | follicular_cells |
| 181 | HLA-DRA | 0.486691263 | 0.428582995 | 0.321994884 | 0.131316138 | 0.143713772 | 0.214535515 | 0.140176271 | 0.15819581 | 0.142622485 | 0.270193161 | 0.165710321 | 0.204988203 | 2.116165368 | 3.824004421 | 3.673832203 | 0.362204979 | 0.24246844 | 0.401340205 | 3.543603343 | 1.311401677 | 4.069205815 | 4.47010321 | follicular_cells |
| 182 | BANK1 | 0.015481786 | 0.007554587 | 0.097006478 | 0.014720941 | 0.008964665 | 0.019815307 | 0.011101624 | 0.009135597 | 0.010437692 | 0.013252004 | 0.01743386 | 0.017399691 | 0.390804515 | 1.577772095 | 0.914376281 | 0.146997354 | 0.11828125 | 0.025601124 | 0.125046345 | 0.074886039 | 0.059012333 | 0.070241232 | follicular_cells |
| 183 | HLA-DRB1 | 0.916674799 | 0.901724742 | 0.246993479 | 0.099809316 | 0.138745989 | 0.316370113 | 0.116528724 | 0.198408146 | 0.174915544 | 0.421869397 | 0.144856625 | 0.28089219 | 1.682360484 | 3.008090844 | 2.855933169 | 0.454752253 | 0.317584618 | 0.477116052 | 3.072090739 | 1.168482783 | 3.408480536 | 3.926667206 | follicular_cells |
| 184 | HLA-DPB1 | 0.808800022 | 0.989896021 | 0.161004197 | 0.099218943 | 0.122358612 | 0.251630238 | 0.126518052 | 0.222480671 | 0.261137931 | 0.38630773 | 0.155143197 | 0.234581684 | 1.45089279 | 2.644226731 | 2.66034071 | 0.246928793 | 0.120161143 | 0.295635573 | 2.503118703 | 0.639773336 | 2.891284352 | 3.566368079 | follicular_cells |
| 185 | HLA-DQA1 | 0.221796709 | 0.171745145 | 0.170521691 | 0.029638483 | 0.029678854 | 0.068480353 | 0.03200101 | 0.052060319 | 0.023761728 | 0.074787245 | 0.039016379 | 0.052733327 | 1.038529491 | 2.178390373 | 2.224251017 | 0.216459344 | 0.132954237 | 0.095067149 | 1.57331729 | 0.618704643 | 2.029273836 | 2.598569831 | follicular_cells |
| 186 | CD37 | 0.994284018 | 1.050698266 | 0.418087021 | 1.066510754 | 1.038702583 | 1.055817675 | 0.979560231 | 1.016266335 | 1.072179726 | 0.949909501 | 1.206364599 | 0.992237132 | 1.397161558 | 2.533459856 | 2.681236986 | 0.471214724 | 0.270858518 | 1.024140221 | 1.333456213 | 0.21386708 | 0.72261142 | 0.52543295 | follicular_cells |
| 187 | LY9 | 0.194988364 | 0.231579705 | 0.140387061 | 0.160684291 | 0.110966862 | 0.122895678 | 0.185571289 | 0.152388796 | 0.209747002 | 0.135496139 | 0.173875621 | 0.126087923 | 0.264680087 | 1.212932488 | 0.769214372 | 0.299249946 | 0.166929035 | 0.052828739 | 0.881711316 | 0.04524734 | 0.114973415 | 0.104925961 | follicular_cells |
| 188 | HLA-DQB1 | 0.50422965 | 0.334099289 | 0.202611289 | 0.07376799 | 0.106316455 | 0.199038938 | 0.051602002 | 0.099756024 | 0.085122879 | 0.20535479 | 0.118002614 | 0.144875902 | 0.97538607 | 2.162742855 | 2.193780643 | 0.300498788 | 0.207874736 | 0.146969101 | 1.803948955 | 0.563167711 | 2.20623685 | 2.552567435 | follicular_cells |
| 189 | REL | 2.020599294 | 0.774716104 | 1.080425085 | 1.077322166 | 1.801142905 | 1.738893306 | 0.392410388 | 2.137821099 | 0.858800917 | 2.090211247 | 1.435361019 | 1.76016016 | 1.323720396 | 1.981326584 | 1.923197425 | 0.618340562 | 0.098226293 | 1.163185981 | 1.694404278 | 1.274260579 | 1.960265803 | 0.969788882 | follicular_cells |
| 190 | CD52 | 1.542364977 | 1.714364155 | 0.581510177 | 1.723928394 | 1.803815578 | 1.372551348 | 1.719176129 | 1.826763902 | 2.055587708 | 1.335432917 | 1.617204176 | 2.027027675 | 1.675816487 | 2.287954901 | 2.229724615 | 0.625499453 | 0.209882866 | 1.088284007 | 0.524776365 | 0.348175934 | 0.984359558 | 0.806405101 | follicular_cells |
| 191 | LINC00926 | 0.01642797 | 0.011865044 | 0.050965844 | 0.013107721 | 0.011798305 | 0.015122253 | 0.011108285 | 0.00981751 | 0.010871873 | 0.009368252 | 0.014025679 | 0.013716561 | 0.066659925 | 0.844192836 | 0.292767365 | 0.038205863 | 0.021799918 | 0.018438416 | 0.007982577 | 0.049541869 | 0.031823253 | 0.044733107 | follicular_cells |
| 192 | MS4A1 | 0.09618946 | 0.091114992 | 0.162833937 | 0.043729502 | 0.030352361 | 0.055210103 | 0.053691763 | 0.046551441 | 0.039821786 | 0.045743934 | 0.058440378 | 0.045521528 | 1.377001924 | 2.436555745 | 2.930094004 | 0.106596229 | 0.062420576 | 0.048581313 | 0.027483124 | 0.121214494 | 0.062239223 | 0.105539897 | follicular_cells |
| 193 | CD74 | 0.98635857 | 1.069891347 | 0.504168683 | 0.44557374 | 0.535999937 | 0.832947631 | 0.415721283 | 0.517103557 | 0.501160514 | 0.758091311 | 0.541778107 | 0.660572909 | 2.256082698 | 3.377647786 | 3.553314569 | 1.352860307 | 1.36082025 | 0.641327316 | 3.654560968 | 1.802555167 | 3.31209155 | 3.747647281 | follicular_cells |
| 194 | ZNF331 | 1.504867096 | 0.46055346 | 0.64591837 | 0.619991369 | 1.298734729 | 1.327619721 | 0.229460398 | 1.406623544 | 0.466864938 | 1.338530808 | 1.351972737 | 1.055364486 | 0.391232803 | 0.995552385 | 0.414672178 | 0.484399348 | 0.11714461 | 0.608381359 | 1.491972123 | 1.205993317 | 1.289306593 | 0.815235993 | follicular_cells |
| 195 | NFKBID | 0.791651297 | 0.310328636 | 0.734534613 | 0.609099681 | 0.524046994 | 0.386080233 | 0.156305847 | 0.801450095 | 0.52267536 | 0.750415855 | 0.410381951 | 0.527157892 | 0.491083845 | 1.263244103 | 0.855593207 | 0.505790281 | 0.097536539 | 0.662278714 | 0.937514485 | 0.327206358 | 0.640512279 | 0.300995901 | follicular_cells |
| 196 | RPS27 | 4.393412684 | 4.512183177 | 2.670608076 | 4.435879843 | 4.589947254 | 4.271196505 | 4.422936735 | 4.315003149 | 4.421529339 | 4.263106891 | 4.498411268 | 4.180827574 | 3.968199375 | 4.760872673 | 4.422498118 | 3.426068933 | 3.291805443 | 3.892206018 | 3.909382602 | 2.51541187 | 3.592000443 | 3.543098741 | follicular_cells |
| 197 | TUBA1A | 1.179183099 | 0.893406813 | 1.474610665 | 1.210798659 | 1.156346211 | 1.075887837 | 0.902837096 | 1.335704297 | 1.166806307 | 1.073160566 | 1.069239406 | 1.055291492 | 0.619943651 | 1.055498313 | 0.490811582 | 0.359302671 | 0.079971134 | 1.461770574 | 0.664268907 | 1.171579806 | 0.930160359 | 0.72871207 | follicular_cells |
| 198 | HLA-DRB5 | 0.2105628 | 0.199658128 | 0.091314392 | 0.020356025 | 0.021086155 | 0.072747064 | 0.017518397 | 0.035015736 | 0.031219427 | 0.083471418 | 0.029677528 | 0.058165153 | 0.641509131 | 1.187335415 | 1.10715081 | 0.117474757 | 0.072247937 | 0.14876151 | 0.881841539 | 0.495232161 | 1.711423698 | 1.901890237 | follicular_cells |
| 199 | VPREB3 | 0.013687267 | 0.012324187 | 0.064679599 | 0.012067622 | 0.008355812 | 0.01667655 | 0.005355129 | 0.013282826 | 0.006122899 | 0.017825717 | 0.01197376 | 0.014315764 | 0.839224038 | 1.579976199 | 1.82400324 | 0.101004755 | 0.086833482 | 0.016430871 | 0.013947312 | 0.049959912 | 0.02280411 | 0.088682972 | follicular_cells |
| 200 | MARCKSL1 | 0.259051793 | 0.137747701 | 0.31183482 | 0.15353399 | 0.209382126 | 0.151876911 | 0.099282147 | 0.214179365 | 0.142609381 | 0.227978533 | 0.157305046 | 0.184620787 | 1.081730479 | 0.557582734 | 2.155593097 | 0.186902628 | 0.097912063 | 0.247127446 | 0.346700558 | 0.386145415 | 0.566109144 | 0.246048415 | gc_cells |
| 201 | LMO2 | 0.001793275 | 0.00240603 | 0.009689964 | 0.002800368 | 0.000420851 | 0.002796063 | 0.003152807 | 0.000889015 | 0.001873412 | 0.006655254 | 0.003425482 | 0.002075155 | 0.420857696 | 0.138930026 | 1.4289686 | 0.005870815 | 0.002844666 | 0.109789113 | 0.04745033 | 0.061243213 | 0.370278362 | 0.383471868 | gc_cells |
| 202 | ACTG1 | 2.349880199 | 2.669357572 | 0.954705811 | 2.117652829 | 2.144664149 | 2.369969931 | 2.082955926 | 2.192692529 | 2.561833974 | 2.610891041 | 2.434441823 | 2.38994064 | 3.384171787 | 2.158180844 | 3.753454642 | 1.510815254 | 1.071405697 | 1.87549271 | 2.202842798 | 1.338026505 | 2.651779243 | 2.619745613 | gc_cells |
| 203 | RGS13 | 0.008830432 | 0.005683523 | 0.008321209 | 0.003628943 | 0.005600436 | 0.014464362 | 0.002101471 | 0.007738222 | 0.002302014 | 0.014750891 | 0.010364632 | 0.007305363 | 0.758559937 | 0.035279815 | 1.271953121 | 0.052253858 | 0.032144738 | 0.453252548 | 0.383443175 | 0.003231019 | 0.026811755 | 0.014655616 | gc_cells |
| 204 | LPP | 0.2429454 | 0.244351768 | 0.167038636 | 0.210969354 | 0.21929117 | 0.270422671 | 0.182884445 | 0.228311625 | 0.219815963 | 0.363088198 | 0.278919374 | 0.234887849 | 0.867478253 | 0.258539249 | 1.420183297 | 0.120523018 | 0.045128025 | 0.331406791 | 0.37302196 | 0.150822151 | 0.356182855 | 0.279777413 | gc_cells |
| 205 | LRMP | 0.218215132 | 0.18637619 | 0.154651152 | 0.099924025 | 0.113049641 | 0.075045631 | 0.116094459 | 0.149627312 | 0.114514247 | 0.231655515 | 0.212809786 | 0.156398884 | 1.098419659 | 0.357554838 | 1.675588325 | 0.215997838 | 0.149205137 | 0.136309801 | 0.181373688 | 0.104703075 | 0.188914075 | 0.137820691 | gc_cells |
| 206 | NEIL1 | 0.017010957 | 0.026675893 | 0.029862057 | 0.016017745 | 0.018164394 | 0.023514932 | 0.020741013 | 0.017752168 | 0.021250562 | 0.054603593 | 0.013125294 | 0.01523603 | 0.611270778 | 0.262069359 | 1.374971557 | 0.088261472 | 0.067541134 | 0.03449015 | 0 | 0.015494595 | 0.009435865 | 0.015162499 | gc_cells |
| 207 | BASP1 | 0.012446232 | 0.013462833 | 0.051821475 | 0.011483303 | 0.014524721 | 0.064697508 | 0.008096867 | 0.01174906 | 0.00776218 | 0.025297477 | 0.042022169 | 0.017520295 | 1.18871844 | 0.928785742 | 2.14741228 | 0.486266581 | 0.340134832 | 0.282137818 | 0.150894192 | 0.208039399 | 1.113817243 | 0.539659199 | gc_cells |
| 208 | SYNE2 | 0.97424482 | 1.186476478 | 1.16393248 | 0.882300082 | 0.967787928 | 1.173174797 | 1.078330071 | 1.019519198 | 0.987277281 | 0.695582487 | 0.872188781 | 0.965098579 | 1.281425816 | 0.20249048 | 1.448240894 | 0.690388383 | 0.509906621 | 0.229620649 | 0.050011801 | 0.084667951 | 0.066608727 | 0.064967978 | gc_cells |
| 209 | TCL1A | 0.007037684 | 0.004483102 | 0.015401636 | 0.002629345 | 0.003467249 | 0.006718893 | 0.004809896 | 0.003057248 | 0.00776466 | 0.004993696 | 0.004891189 | 0.004607556 | 1.120129531 | 0.577681163 | 1.613616132 | 0.012939376 | 0.012600066 | 0.005513983 | 0.745067595 | 0.014647991 | 0.006594206 | 0.017897339 | gc_cells |
| 210 | CD79B | 0.044628445 | 0.103223019 | 0.09279292 | 0.060640189 | 0.04666791 | 0.234272789 | 0.098060618 | 0.029617987 | 0.03102888 | 0.062572375 | 0.148525111 | 0.072808666 | 1.144830237 | 1.240782766 | 2.253276149 | 0.383686824 | 0.385216563 | 0.100630265 | 0.031138644 | 0.046377863 | 0.040104657 | 0.081116068 | gc_cells |
| 211 | HMGN1 | 0.918918558 | 0.935423463 | 0.585194672 | 1.061973007 | 1.02109322 | 1.130343386 | 1.113438232 | 1.067554676 | 1.11933387 | 1.186065581 | 1.302388612 | 1.19605638 | 2.37343366 | 1.250657619 | 2.461931111 | 0.960491284 | 0.770949199 | 1.229999536 | 1.460285148 | 0.46115948 | 0.853250987 | 0.964698718 | gc_cells |
| 212 | ACTB | 2.943931941 | 3.321888939 | 1.497078531 | 2.734702084 | 2.604686136 | 3.082795679 | 2.708267208 | 2.78381546 | 3.277405304 | 3.21487756 | 2.964013942 | 3.114427802 | 3.961982749 | 2.868002469 | 3.975062004 | 1.848163893 | 1.134690745 | 2.58866209 | 2.947540499 | 2.122639058 | 3.87971385 | 3.88869118 | gc_cells |
| 213 | SMIM14 | 0.060740775 | 0.072862458 | 0.060311758 | 0.061759257 | 0.053904996 | 0.032811173 | 0.058187823 | 0.053271213 | 0.058841585 | 0.102603088 | 0.041102505 | 0.043311118 | 0.699385835 | 1.070559652 | 1.847333394 | 0.160495072 | 0.178786454 | 0.222729703 | 0.534982355 | 0.254812145 | 0.213173198 | 0.277795525 | gc_cells |
| 214 | SNX29P2 | 0.147139769 | 0.121307717 | 0.110580528 | 0.109996579 | 0.117500506 | 0.13826234 | 0.122573079 | 0.131791693 | 0.097460008 | 0.160005906 | 0.150657996 | 0.121405557 | 0.726468346 | 0.641429975 | 1.535595784 | 0.123607861 | 0.075845558 | 0.107501554 | 0.347038387 | 0.100764866 | 0.141707892 | 0.108645728 | gc_cells |
| 215 | SUGCT | 0.000517592 | 0.001414781 | 0.002360597 | 0.00098597 | 0.000826094 | 0.000353924 | 0.002040112 | 0.001703069 | 0.002562442 | 0 | 0.000239907 | 0.001741058 | 0.401425206 | 0.012829716 | 0.731060524 | 0.001617786 | 0.000822307 | 0.003171638 | 0 | 0 | 0.001225485 | 0.001154297 | gc_cells |
| 216 | RFTN1 | 0.177690883 | 0.209790872 | 0.10885813 | 0.114027776 | 0.151699997 | 0.222378528 | 0.126558198 | 0.170974231 | 0.113614398 | 0.190205002 | 0.145335044 | 0.16358365 | 0.713111503 | 0.306509364 | 1.281661869 | 0.115012607 | 0.052714404 | 0.212981233 | 0.574639616 | 0.118719003 | 0.306742218 | 0.171188728 | gc_cells |
| 217 | LIMD2 | 0.774262515 | 0.956379942 | 0.59012142 | 0.701094395 | 0.658194393 | 0.768676628 | 0.856442243 | 0.660625487 | 0.89352392 | 0.777770401 | 0.836963002 | 0.662947819 | 1.257456284 | 1.013743814 | 1.94237436 | 0.320113318 | 0.167287844 | 0.414487764 | 0.456210884 | 0.298989594 | 0.469375923 | 0.383894467 | gc_cells |
| 218 | CORO1A | 1.30571641 | 1.605149984 | 0.63913652 | 1.1198249 | 1.091677346 | 1.283010918 | 1.178552801 | 1.238188188 | 1.413193787 | 1.411359634 | 1.039789588 | 1.382442838 | 2.055382706 | 1.340272214 | 2.304722901 | 0.477510777 | 0.226894676 | 0.387175607 | 1.125384893 | 0.441603308 | 0.997671272 | 0.879416549 | gc_cells |
| 219 | ARPC1B | 0.991870666 | 1.28365746 | 0.289573621 | 0.905847269 | 1.007133519 | 1.425599658 | 0.969474879 | 0.986465512 | 1.24666603 | 1.062405575 | 1.272078558 | 1.254007182 | 1.829394086 | 0.988959312 | 2.039454717 | 0.489786864 | 0.273419911 | 0.806389537 | 1.414231205 | 0.329238682 | 1.66510193 | 1.946439093 | gc_cells |
| 220 | MYBL1 | 0.263169541 | 0.436359231 | 0.19810564 | 0.26096216 | 0.245775797 | 0.061775845 | 0.199436723 | 0.439557034 | 0.477276348 | 0.385543492 | 0.043734405 | 0.286671028 | 0.859728979 | 0.023980628 | 0.866015475 | 0.06631541 | 0.01736555 | 0.039327318 | 0 | 0.051179952 | 0.031996688 | 0.026015755 | gc_cells |
| 221 | UCP2 | 0.335065301 | 0.428622649 | 0.257512677 | 0.28547525 | 0.4162656 | 0.827128177 | 0.292770562 | 0.424144094 | 0.305720101 | 0.331408615 | 0.710920237 | 0.521153764 | 1.022655389 | 0.52170062 | 1.43552103 | 0.232048853 | 0.142127152 | 0.230785815 | 0.95733951 | 0.22317746 | 0.74909712 | 0.859111847 | gc_cells |
| 222 | BCL7A | 0.007916964 | 0.007459453 | 0.014117987 | 0.01248721 | 0.012060403 | 0.007769399 | 0.017106589 | 0.016583113 | 0.008833352 | 0.01163783 | 0.0123714 | 0.005523172 | 0.685144333 | 0.167140247 | 1.01205378 | 0.025144678 | 0.017745201 | 0.055266235 | 0.525785069 | 0.018854865 | 0.094011726 | 0.071284154 | gc_cells |
| 223 | ANP32B | 0.542048391 | 0.579693177 | 0.233899892 | 0.732397985 | 0.818796911 | 0.864354821 | 0.76211437 | 0.680181412 | 0.730230638 | 0.646110776 | 1.008449628 | 0.695161697 | 2.176181761 | 0.829461677 | 1.932040732 | 0.644450252 | 0.553476335 | 0.715353489 | 0.831557943 | 0.159058624 | 0.652674131 | 0.725795596 | gc_cells |
| 224 | LCP1 | 1.116256662 | 1.252961818 | 0.366519044 | 0.829068207 | 0.890481966 | 0.824401214 | 0.782730341 | 0.987504222 | 1.089414594 | 1.246916155 | 0.793292863 | 1.145660297 | 1.613933385 | 0.87263732 | 1.80605688 | 0.351806241 | 0.119837043 | 0.700435134 | 1.159735187 | 0.56144571 | 1.728106795 | 1.379243852 | gc_cells |
| 225 | CD22 | 0.019324563 | 0.014771082 | 0.087259574 | 0.014234138 | 0.011665826 | 0.01687344 | 0.020501124 | 0.01385974 | 0.014782825 | 0.005756959 | 0.0180707 | 0.016831951 | 0.79257919 | 1.074284745 | 1.830868867 | 0.052204189 | 0.029065581 | 0.335058429 | 0 | 0.246545707 | 0.099665958 | 0.098599598 | gc_cells |
| 226 | HMCES | 0.204732467 | 0.141043259 | 0.110850008 | 0.13615652 | 0.173909164 | 0.228142056 | 0.113182639 | 0.209145299 | 0.152127153 | 0.202250384 | 0.254702788 | 0.200712565 | 0.741541004 | 0.084443107 | 1.009794235 | 0.318630466 | 0.184921457 | 0.136986986 | 0.097726163 | 0.044156858 | 0.062742506 | 0.059784036 | gc_cells |
| 227 | ARPC2 | 1.568722273 | 1.654433036 | 0.572077206 | 1.254458447 | 1.413143519 | 1.538687597 | 1.13752387 | 1.528124322 | 1.537141156 | 1.729587791 | 1.446639264 | 1.631826708 | 2.353849068 | 1.397804718 | 2.462628339 | 1.104967943 | 0.937808139 | 1.13474706 | 1.63858031 | 0.539277149 | 1.904962191 | 1.960960892 | gc_cells |
| 228 | SERPINA9 | 0.001486152 | 0.0015775 | 0.008279789 | 0.000409241 | 0 | 0.000662883 | 0 | 0.002491457 | 0.000841527 | 0.003942718 | 0.00137822 | 0.001416741 | 0.270598434 | 0.019411773 | 0.720206417 | 0.002780691 | 0.002488928 | 0.001385507 | 0 | 0 | 0.002216117 | 0 | gc_cells |
| 229 | CCDC144A | 0.078516218 | 0.099674237 | 0.167656999 | 0.07679698 | 0.063601899 | 0.120600548 | 0.127516173 | 0.06249284 | 0.09602078 | 0.101174469 | 0.109649879 | 0.088780933 | 0.367704624 | 0.031596559 | 0.897877798 | 0.466818337 | 0.262546512 | 0.13553729 | 0.114649731 | 0.058646403 | 0.057269001 | 0.051805374 | gc_cells |
| 230 | SERF2 | 1.855192247 | 2.044987033 | 0.994370609 | 1.715310281 | 1.861334012 | 1.951532677 | 1.6805394 | 1.883788563 | 1.949007294 | 2.044359665 | 1.980882384 | 2.110153442 | 2.879973034 | 1.883464027 | 3.114378522 | 2.193201252 | 2.320406325 | 2.092849732 | 2.245596984 | 1.54677998 | 2.360819817 | 2.660895881 | gc_cells |
| 231 | ACTR3 | 0.879227172 | 0.963749673 | 0.428326305 | 0.80129131 | 0.87929112 | 0.937276762 | 0.746677341 | 0.953950467 | 0.989292694 | 1.078159187 | 0.939371243 | 1.089576512 | 1.71433307 | 1.058642967 | 1.992480784 | 0.564358637 | 0.39722198 | 0.895804808 | 0.981057526 | 0.370010188 | 1.407866472 | 1.188544353 | gc_cells |
| 232 | ATP5L | 1.315973795 | 1.409875457 | 0.31339031 | 1.312962654 | 1.439399069 | 1.423717696 | 1.27505395 | 1.324039447 | 1.424271824 | 1.396925231 | 1.478234301 | 1.436394318 | 2.283915228 | 1.24003585 | 2.342154111 | 1.411259869 | 1.431207595 | 1.111151051 | 1.673501059 | 0.284373523 | 1.449554045 | 1.552523151 | gc_cells |
| 233 | ARPC5 | 0.610869971 | 0.853566985 | 0.170798122 | 0.606734833 | 0.541248232 | 0.591411959 | 0.664266718 | 0.661493654 | 0.823350431 | 0.816051244 | 0.664431695 | 0.67398667 | 1.654700049 | 0.615668957 | 1.549230415 | 0.43005316 | 0.346124475 | 0.704472659 | 1.209841125 | 0.370435764 | 1.447217984 | 1.495449064 | gc_cells |
| 234 | CD53 | 0.763327929 | 0.937483687 | 0.245968988 | 0.697763893 | 0.667215476 | 0.713052453 | 0.645889421 | 0.773652049 | 0.874519114 | 0.797426797 | 0.615877524 | 0.874238271 | 1.182093924 | 1.093980289 | 1.867405657 | 0.473366557 | 0.36145193 | 0.522402172 | 1.028138629 | 0.251319269 | 1.099725325 | 0.972061445 | gc_cells |
| 235 | PARP1 | 0.249408599 | 0.293444893 | 0.087111767 | 0.182214578 | 0.193493102 | 0.296048765 | 0.216031509 | 0.186497815 | 0.215737635 | 0.22203651 | 0.332583768 | 0.256191759 | 1.097032984 | 0.419097352 | 1.339052962 | 0.371223609 | 0.383017907 | 0.222788735 | 0.219746278 | 0.069473653 | 0.169770583 | 0.203597068 | gc_cells |
| 236 | SLC25A5 | 0.882757082 | 0.679902635 | 0.33635471 | 0.772994091 | 0.922363408 | 0.843592747 | 0.44455635 | 0.956863747 | 0.810261014 | 1.037442523 | 0.851270843 | 0.964615848 | 2.013169745 | 0.95100928 | 1.935030742 | 0.692445429 | 0.555535312 | 0.81834206 | 1.212871761 | 0.215179466 | 1.190934924 | 1.529309071 | gc_cells |
| 237 | AC023590.1 | 0.002675726 | 0.006605558 | 0.002467309 | 0.01223924 | 0.009353624 | 0.013974926 | 0.01170887 | 0.006572999 | 0.004034991 | 0.009226586 | 0.014893689 | 0.011329564 | 0.4056464 | 0.024090033 | 0.766482799 | 0.018554708 | 0.013902447 | 0.008469332 | 0.381421873 | 0.014294148 | 0.01619376 | 0.015552673 | gc_cells |
| 238 | CCDC88A | 0.034938542 | 0.03369753 | 0.052289355 | 0.042467779 | 0.034969184 | 0.013943317 | 0.043016133 | 0.042166999 | 0.048372926 | 0.061868679 | 0.035448924 | 0.020045063 | 0.548834081 | 0.075129958 | 0.874265732 | 0.301715121 | 0.296603778 | 0.441936192 | 1.25959536 | 0.615060939 | 0.79041651 | 0.535926713 | gc_cells |
| 239 | NEAT1 | 2.267684764 | 2.028040041 | 2.776616565 | 1.811022807 | 2.053195611 | 2.406711292 | 1.772920149 | 1.939125965 | 1.625081457 | 2.386305649 | 2.053740047 | 2.247217807 | 1.710308525 | 1.348824886 | 1.547582146 | 3.215109047 | 1.920620285 | 2.774773412 | 2.592278494 | 3.991135182 | 3.910999257 | 2.089229463 | plasma_NFKBIA_hi |
| 240 | ANKRD28 | 0.281952225 | 0.1833791 | 0.778255136 | 0.579540244 | 0.802158223 | 0.234940867 | 0.217002626 | 1.146573528 | 0.534085159 | 0.594017398 | 0.31310885 | 1.23294675 | 0.49229874 | 0.142186477 | 0.166284213 | 1.511305763 | 0.732201897 | 0.969537 | 0.56129457 | 0.324876873 | 0.592117065 | 0.203699739 | plasma_NFKBIA_hi |
| 241 | HSPB1 | 0.511515974 | 0.457913725 | 0.910790065 | 0.799263238 | 0.373355504 | 0.581820384 | 0.578882437 | 0.550816974 | 0.727440237 | 0.474458105 | 0.647751031 | 0.50265212 | 0.492624774 | 0.757056566 | 0.491853506 | 2.016285224 | 1.058867095 | 0.888701646 | 0.8618268 | 0.390428638 | 0.764598595 | 0.738967708 | plasma_NFKBIA_hi |
| 242 | JUN | 3.210620118 | 2.736681661 | 4.302499796 | 3.800899281 | 2.506487616 | 2.607232398 | 2.431497049 | 3.215766293 | 3.480233535 | 3.009208044 | 2.578841315 | 2.970867114 | 1.830709361 | 2.666706319 | 1.613076926 | 4.226402803 | 2.516936694 | 3.557998338 | 2.640974281 | 3.103966741 | 2.068274328 | 1.638237473 | plasma_NFKBIA_hi |
| 243 | HSPA6 | 0.55770074 | 0.396194535 | 0.560660385 | 0.656707922 | 0.195483494 | 0.128058737 | 0.282263724 | 0.470566966 | 0.540899574 | 0.512554597 | 0.245959554 | 0.304879874 | 0.239665661 | 0.260551773 | 0.195067004 | 1.083779986 | 0.40825381 | 0.295191604 | 0.330814367 | 0.758763208 | 0.822836076 | 0.37952549 | plasma_NFKBIA_hi |
| 244 | AHNAK | 0.990770475 | 0.974020232 | 1.107959007 | 0.999102685 | 1.426118778 | 1.10092489 | 0.678952402 | 0.853076663 | 0.604053443 | 0.70432143 | 1.011157179 | 0.893664047 | 0.370853507 | 0.616153539 | 0.118225563 | 1.497272547 | 0.40460352 | 1.238833174 | 0.683912428 | 0.643155098 | 1.019394556 | 0.463597046 | plasma_NFKBIA_hi |
| 245 | BTG2 | 1.846415803 | 1.125677652 | 2.480543772 | 1.918467779 | 1.367427403 | 1.524718662 | 0.932487064 | 1.658483524 | 1.559770914 | 1.58738669 | 1.420393823 | 1.569422441 | 1.173610113 | 1.359423486 | 1.245901508 | 2.67127793 | 1.7322525 | 1.670958995 | 2.391450758 | 1.313689078 | 0.978172256 | 0.683776469 | plasma_NFKBIA_hi |
| 246 | ZFAND2A | 0.666175684 | 0.323783868 | 0.646477187 | 0.802147664 | 0.321012447 | 0.447430947 | 0.240059699 | 0.530928569 | 0.658860489 | 0.53809091 | 0.384700831 | 0.50181893 | 0.306700818 | 0.519425743 | 0.303638229 | 1.186453168 | 0.223786356 | 0.457836824 | 0.675840032 | 0.525138304 | 0.737322857 | 0.394857299 | plasma_NFKBIA_hi |
| 247 | H1FX | 0.650773652 | 0.518332616 | 0.797521317 | 0.714866381 | 0.723950038 | 0.653924339 | 0.664233139 | 0.757824608 | 0.689104845 | 0.684675852 | 0.784581434 | 0.726377836 | 0.880066522 | 0.277911542 | 0.364936755 | 1.332634454 | 0.783540932 | 0.616771465 | 1.087714502 | 0.573616685 | 0.545440132 | 0.417930633 | plasma_NFKBIA_hi |
| 248 | LINC00152 | 0.683502825 | 0.626114633 | 0.475260789 | 0.417107063 | 0.431435107 | 1.143130186 | 0.292224763 | 0.685769366 | 0.640265442 | 0.410831724 | 0.505173425 | 0.72253085 | 0.577625579 | 0.056407832 | 0.11230582 | 0.823408756 | 0.455942978 | 0.182190439 | 0.164723404 | 0.465101487 | 0.662980832 | 0.456000573 | plasma_NFKBIA_hi |
| 249 | IER2 | 1.752269843 | 1.579959876 | 2.197816958 | 1.760882011 | 1.218347315 | 1.287609868 | 1.254719005 | 1.344435016 | 1.623961498 | 1.930786872 | 1.14694659 | 1.336747287 | 1.171873552 | 1.771004057 | 1.427578282 | 2.82840532 | 1.597568122 | 2.204991831 | 0.814945433 | 2.577632461 | 1.371330315 | 1.00120048 | plasma_NFKBIA_hi |
| 250 | PRDM1 | 0.379094246 | 0.323145408 | 0.432129558 | 0.276074213 | 0.393839088 | 0.860615021 | 0.209364149 | 0.363530876 | 0.280602875 | 0.251291386 | 0.319417923 | 0.771474108 | 0.358665232 | 0.033766275 | 0.039235806 | 1.05428049 | 0.728409452 | 0.105965262 | 0.26330614 | 0.430318934 | 0.83330496 | 0.451723852 | plasma_NFKBIA_hi |
| 251 | SQSTM1 | 1.245894443 | 0.898034022 | 1.912682473 | 1.360969371 | 1.40225799 | 1.58901786 | 0.738255793 | 1.403899307 | 1.230070667 | 1.630066871 | 1.02556266 | 1.604843538 | 0.946851341 | 1.290729038 | 0.948879881 | 2.254403654 | 1.110148073 | 1.854951114 | 1.103477931 | 1.339431424 | 1.695811847 | 1.254498122 | plasma_NFKBIA_hi |
| 252 | DNAJB4 | 0.513301105 | 0.29808803 | 1.059073425 | 0.680721503 | 0.272338357 | 0.599231275 | 0.34019856 | 0.613293864 | 0.652395954 | 0.467954559 | 0.4609862 | 0.446871953 | 0.227341609 | 0.35571317 | 0.176661709 | 1.068407624 | 0.334334767 | 0.549792843 | 0.609483924 | 0.418129827 | 0.236380054 | 0.164139469 | plasma_NFKBIA_hi |
| 253 | RGCC | 1.399536719 | 0.390805143 | 0.632953893 | 0.912899593 | 1.749934054 | 1.454239028 | 0.336407965 | 1.531377112 | 0.478019457 | 1.180075044 | 1.330452187 | 1.145461904 | 0.609456441 | 0.1790277 | 0.227903203 | 0.991182583 | 0.438272632 | 0.428928107 | 0.334411332 | 0.205322998 | 0.93467845 | 0.564017114 | plasma_NFKBIA_hi |
| 254 | CITED2 | 0.738178673 | 0.465205713 | 0.874721983 | 0.856267105 | 0.624429021 | 0.440365781 | 0.383377361 | 0.835255847 | 0.845303134 | 0.624002116 | 0.637779678 | 0.812417262 | 0.461530493 | 0.230746736 | 0.342442398 | 1.175607076 | 0.710078067 | 0.473838901 | 0.28394783 | 0.339649358 | 0.366583407 | 0.357654623 | plasma_NFKBIA_hi |
| 255 | ATF3 | 0.847101081 | 0.173007506 | 0.658322811 | 0.618839053 | 0.620905483 | 0.500322661 | 0.070434511 | 1.400622644 | 0.597320295 | 0.89341342 | 0.28092929 | 1.182176361 | 0.402663061 | 0.252253047 | 0.110090542 | 0.852712992 | 0.167204063 | 0.819096722 | 1.199391292 | 0.976694654 | 1.920160232 | 0.988020371 | plasma_NFKBIA_hi |
| 256 | NFKBIA | 1.843296829 | 1.046815425 | 2.058193625 | 2.2793264 | 2.310715351 | 1.868289582 | 0.805761698 | 2.105506002 | 1.650881376 | 2.069552783 | 1.572601452 | 2.210251185 | 1.215496228 | 1.55333662 | 1.200709756 | 2.435747115 | 0.850311105 | 2.238860615 | 1.081737215 | 2.622676959 | 2.934199634 | 1.532301423 | plasma_NFKBIA_hi |
| 257 | TMEM107 | 0.17454347 | 0.212337711 | 0.494688766 | 0.207970795 | 0.124643643 | 0.105041266 | 0.16978791 | 0.186174473 | 0.298688198 | 0.296031313 | 0.142750824 | 0.244983221 | 0.314887815 | 0.155593815 | 0.173999168 | 0.902444761 | 0.449585614 | 0.31184426 | 0.880108671 | 0.583590282 | 0.240597538 | 0.189979797 | plasma_NFKBIA_hi |
| 258 | TNFAIP3 | 2.402549002 | 0.883038042 | 1.999956135 | 2.464295368 | 3.210698035 | 2.372752719 | 0.728186087 | 3.191052556 | 1.579347853 | 2.190765421 | 1.608349951 | 3.046887773 | 0.960872345 | 0.707173047 | 0.462834919 | 1.479801856 | 0.393017514 | 1.488975754 | 0.492909749 | 1.656383563 | 2.260808268 | 1.192867419 | plasma_NFKBIA_hi |
| 259 | UBC | 3.137200817 | 2.441247961 | 3.053280592 | 3.008343851 | 2.994444888 | 3.359659088 | 1.893011277 | 3.306684838 | 2.678676846 | 2.833847072 | 2.920954096 | 3.268384658 | 2.453510016 | 2.510911596 | 2.279941573 | 3.542161089 | 2.329717923 | 2.494495291 | 3.353452081 | 2.101658875 | 3.040766527 | 2.452469354 | plasma_NFKBIA_hi |
| 260 | TRIB1 | 0.051945596 | 0.025782977 | 0.093049582 | 0.031603306 | 0.031321058 | 0.309828378 | 0.021273558 | 0.022980626 | 0.02635524 | 0.046309078 | 0.246657304 | 0.064434629 | 0.205105419 | 0.05546679 | 0.12820838 | 0.794335871 | 0.370651264 | 0.091028458 | 0.6613122 | 0.258952974 | 0.500248493 | 0.252424191 | plasma_NFKBIA_hi |
| 261 | HERPUD1 | 0.757475416 | 0.571810768 | 0.699898702 | 0.491778599 | 0.548111651 | 0.781358939 | 0.328827946 | 0.540852781 | 0.490824267 | 0.757537332 | 0.551518756 | 0.590224602 | 1.226716305 | 1.435277961 | 1.667311424 | 2.82173628 | 2.537342564 | 0.656443017 | 2.312976556 | 1.290502099 | 1.170557119 | 0.89922078 | plasma_NFKBIA_lo |
| 262 | CPEB4 | 0.247029294 | 0.226350883 | 0.491967569 | 0.196173075 | 0.174558874 | 0.323033644 | 0.203095647 | 0.209010865 | 0.201419432 | 0.234643809 | 0.249024617 | 0.290102168 | 0.282664249 | 0.22004503 | 0.24695662 | 1.092969379 | 0.835672174 | 0.364006142 | 0.319506889 | 0.610720628 | 0.509914545 | 0.299131008 | plasma_NFKBIA_lo |
| 263 | FOSB | 2.325624866 | 1.460761163 | 1.72824248 | 2.397169064 | 2.159665809 | 1.799407416 | 1.170985008 | 2.652089719 | 2.102375577 | 2.488320249 | 1.661544489 | 2.149426206 | 1.22959967 | 1.941705715 | 1.156130769 | 2.920064623 | 1.578944536 | 2.972851627 | 2.652177877 | 2.255263365 | 2.476761386 | 1.370053848 | plasma_NFKBIA_lo |
| 264 | FOS | 3.358462788 | 2.31570215 | 3.593877487 | 2.786335982 | 2.447508266 | 1.970302822 | 2.464161244 | 2.597320955 | 3.277394811 | 3.249128929 | 2.794297925 | 2.652403957 | 1.739945499 | 3.503831518 | 1.825526944 | 3.539104603 | 1.816818301 | 3.610457028 | 3.553592972 | 2.258442809 | 2.851749978 | 2.274056968 | plasma_NFKBIA_lo |
| 265 | IGLL5 | 0.368420828 | 0.336344862 | 0.532550936 | 0.28485234 | 0.283369024 | 0.381724965 | 0.336269707 | 0.223229363 | 0.200358905 | 0.301990784 | 0.378078418 | 0.289071118 | 1.266035831 | 1.011233469 | 1.343468535 | 3.416340921 | 4.37172425 | 0.30017058 | 0.515297866 | 0.887086176 | 0.429588586 | 0.435716548 | plasma_NFKBIA_lo |
| 266 | IGJ | 0.283307398 | 0.258531483 | 0.437042438 | 0.245241614 | 0.233347745 | 0.272986817 | 0.267784748 | 0.232643716 | 0.217549134 | 0.244015978 | 0.28191594 | 0.26521481 | 0.960610967 | 0.446099494 | 0.859018388 | 4.171499085 | 4.542021029 | 0.219074229 | 2.20626307 | 0.553877126 | 0.325417277 | 0.264374423 | plasma_NFKBIA_lo |
| 267 | SSR4 | 0.998646046 | 1.137544348 | 0.275012279 | 0.928833709 | 1.018125289 | 0.913926984 | 0.88052002 | 0.946393286 | 1.085948074 | 1.012336502 | 0.996151888 | 1.035356152 | 0.961407208 | 0.786193659 | 0.774072109 | 3.464748493 | 4.096737384 | 1.138397056 | 1.957093538 | 0.422223766 | 1.154939602 | 1.150452485 | plasma_NFKBIA_lo |
| 268 | MZB1 | 0.106844854 | 0.085080312 | 0.157553614 | 0.074931644 | 0.056650118 | 0.194046659 | 0.069953826 | 0.105231073 | 0.129276799 | 0.172455603 | 0.087643116 | 0.108853363 | 0.809092533 | 0.243871417 | 0.570108662 | 2.771788489 | 3.357389718 | 0.036148954 | 1.548120744 | 0.156823219 | 0.105968147 | 0.110723836 | plasma_NFKBIA_lo |
| 269 | DERL3 | 0.01844112 | 0.023191934 | 0.039967505 | 0.018534734 | 0.020119867 | 0.028392722 | 0.019487445 | 0.018542865 | 0.020648395 | 0.031826272 | 0.025814276 | 0.023224058 | 0.249260224 | 0.108676247 | 0.090409846 | 1.788495489 | 2.403090177 | 0.032490357 | 0.748077861 | 0.070996721 | 0.04715376 | 0.034171139 | plasma_NFKBIA_lo |
| 270 | FKBP11 | 0.313189521 | 0.324119299 | 0.148379575 | 0.459698686 | 0.480564447 | 0.132630334 | 0.3580709 | 0.548841269 | 0.463097619 | 0.391572179 | 0.202645308 | 0.688882253 | 0.360589259 | 0.11042566 | 0.083185982 | 1.843907277 | 2.788866102 | 0.231323071 | 1.155184465 | 0.053088146 | 0.089124489 | 0.050129202 | plasma_NFKBIA_lo |
| 271 | RP11-731F5.2 | 0.008059958 | 0.00341704 | 0.009852303 | 0.005444201 | 0.004073888 | 0.006604977 | 0.003489572 | 0.003235713 | 0 | 0.01202359 | 0.000357584 | 0.000468152 | 0.039415762 | 0.014888203 | 0.015258003 | 0.576909975 | 0.847177355 | 0.010051254 | 0.00856978 | 0.005755362 | 0.01064495 | 0.009611123 | plasma_NFKBIA_lo |
| 272 | XBP1 | 0.536335626 | 0.482730992 | 0.458136044 | 0.505037432 | 0.626671217 | 0.319676511 | 0.325960925 | 0.540604155 | 0.408152881 | 0.726097586 | 0.437861747 | 0.554592231 | 0.506078177 | 0.244331375 | 0.295060453 | 2.585396807 | 2.878056029 | 0.611101608 | 1.472527523 | 1.191504786 | 0.878263843 | 0.434291329 | plasma_NFKBIA_lo |
| 273 | PRDX4 | 0.055212795 | 0.074677457 | 0.019144524 | 0.064949235 | 0.061690673 | 0.089577377 | 0.093225076 | 0.05073983 | 0.07457693 | 0.075486362 | 0.086001812 | 0.050128347 | 0.1993357 | 0.023880936 | 0.076882019 | 1.165935468 | 1.685709641 | 0.181883231 | 0.033729756 | 0.126454314 | 0.204545578 | [illegible] | plasma_NFKBIA_lo |
| 274 | SEC11C | 0.344443511 | 0.405107685 | 0.10659633 | 0.309230772 | 0.385913892 | 0.496067533 | 0.341657527 | 0.350759171 | 0.367262527 | 0.357451795 | 0.518802865 | 0.388436837 | 0.765294471 | 0.277823074 | 0.490007371 | 1.951751108 | 2.458453783 | 0.302954665 | 0.765752514 | 0.078267813 | 0.328339953 | 0.394994257 | plasma_NFKBIA_lo |
| 275 | FKBP2 | 0.339651119 | 0.464551242 | 0.151153731 | 0.379224599 | 0.392185486 | 0.271685806 | 0.349986063 | 0.462613995 | 0.499588836 | 0.486322251 | 0.336451617 | 0.42853042 | 0.517096776 | 0.291321752 | 0.376908816 | 1.970791166 | 2.407560401 | 0.816982083 | 1.184218549 | 0.288331372 | 0.620761744 | 0.704078177 | plasma_NFKBIA_lo |
| 276 | HSP90B1 | 0.90714148 | 0.831954234 | 0.677937893 | 0.840853701 | 1.062781875 | 0.795149304 | 0.737692257 | 1.052345784 | 0.873602995 | 1.069716808 | 0.804551441 | 1.14194577 | 1.00135693 | 0.806994224 | 0.511477437 | 2.741961078 | 3.089142366 | 1.278815866 | 2.385768031 | 0.988650996 | 1.454493375 | 1.43428727 | plasma_NFKBIA_lo |
| 277 | SDF2L1 | 0.300823325 | 0.377420269 | 0.08624543 | 0.230714149 | 0.229555098 | 0.315507153 | 0.18123938 | 0.280071137 | 0.338249389 | 0.399255094 | 0.251707139 | 0.351525728 | 0.515521021 | 0.130922789 | 0.18837886 | 1.364161559 | 1.671000922 | 0.322821005 | 0.403798663 | 0.175671564 | 0.55959027 | 0.548636943 | plasma_NFKBIA_lo |
| 278 | SSR3 | 0.3612608 | 0.381074586 | 0.115866234 | 0.383856078 | 0.476431453 | 0.40309513 | 0.330252266 | 0.397575827 | 0.405857312 | 0.37826182 | 0.417145699 | 0.49773279 | 0.676297664 | 0.268734599 | 0.374360049 | 1.573506603 | 1.842955404 | 0.502457811 | 1.040219692 | 0.156683986 | 0.672157107 | 0.671349247 | plasma_NFKBIA_lo |
| 279 | CD63 | 0.611955903 | 0.78094357 | 0.113593362 | 0.412475323 | 0.375983613 | 0.411163384 | 0.349303669 | 0.638161759 | 0.821060594 | 1.342376603 | 0.280794995 | 0.559350553 | 0.406262444 | 0.280898298 | 0.133245358 | 1.239678206 | 1.660736915 | 0.225454575 | 0.740872495 | 0.395262933 | 1.860459835 | 2.247355251 | plasma_NFKBIA_lo |
| 280 | RRBP1 | 0.210533126 | 0.192160149 | 0.220423794 | 0.136492737 | 0.147300681 | 0.164164733 | 0.138751933 | 0.155521528 | 0.158222177 | 0.321700411 | 0.1728581 | 0.192472428 | 0.380279353 | 0.096935413 | 0.312715954 | 1.651124472 | 1.761545287 | 0.641403423 | 1.063431526 | 0.614205684 | 0.492407657 | 0.467085129 | plasma_NFKBIA_lo |
| 281 | TMEM59 | 0.61309826 | 0.772255868 | 0.138254425 | 0.552551546 | 0.578522898 | 0.688047288 | 0.715684181 | 0.639803222 | 0.711348124 | 0.62544878 | 0.672520201 | 0.694330235 | 0.630293128 | 0.457017336 | 0.523587253 | 1.274341417 | 1.730035297 | 0.766170748 | 0.852746275 | 0.07366291 | 0.572680798 | 0.715073408 | plasma_NFKBIA_lo |
| 282 | TNFRSF17 | 0.009043077 | 0.005430261 | 0.008012772 | 0.007018823 | 0.005743657 | 0.007691934 | 0.006909447 | 0.007643259 | 0.007212964 | 0.008393231 | 0.008350264 | 0.006637133 | 0.267629914 | 0.083668714 | 0.27693455 | 1.163322637 | 1.429514275 | 0.010828952 | 0.120119941 | 0.008666896 | 0.018433607 | 0.017655004 | plasma_NFKBIA_lo |
| 283 | MANF | 0.317827605 | 0.331398607 | 0.095947425 | 0.248502629 | 0.300996809 | 0.245563194 | 0.188410825 | 0.321868356 | 0.315669783 | 0.350044159 | 0.227749371 | 0.35829672 | 0.417999354 | 0.153747129 | 0.216864821 | 1.113148971 | 1.425690178 | 0.381678271 | 0.798423113 | 0.17723348 | 0.429821082 | 0.385177115 | plasma_NFKBIA_lo |
| 284 | SPCS3 | 0.485191385 | 0.612040039 | 0.266034052 | 0.529471823 | 0.56065732 | 0.48338174 | 0.547912465 | 0.601229016 | 0.682037099 | 0.599021171 | 0.539243256 | 0.710602436 | 0.726352053 | 0.364473323 | 0.443791484 | 1.438316403 | 1.818998435 | 0.607972992 | 1.114086287 | 0.239054942 | 0.741868952 | 0.672659584 | plasma_NFKBIA_lo |

| | A | B | C | D | E | F | G | H | I | J | K | L | M | N | O | P | Q | R | S | T | U | V | W | X |
|---|---|---|---|---|---|---|---|---|---|---|---|---|---|---|---|---|---|---|---|---|---|---|---|---|
| 285 | SPCS1 | 0.573130437 | 0.661457559 | 0.140705269 | 0.594831539 | 0.589715424 | 0.726275747 | 0.586030681 | 0.648431295 | 0.733799429 | 0.659213657 | 0.845287657 | 0.690467368 | 0.880610375 | 0.544030697 | 0.720380357 | 1.741694292 | 2.089598508 | 0.780209386 | 1.704477128 | 0.091585715 | 0.562957091 | 0.677963312 | plasma_NFKBIA_lo |
| 286 | SPCS2 | 0.702426297 | 0.790135973 | 0.194246723 | 0.618446971 | 0.682743928 | 0.693140069 | 0.528893468 | 0.675646088 | 0.677803317 | 0.795708661 | 0.833495522 | 0.851914056 | 1.187548024 | 0.695289824 | 1.099497184 | 1.854511133 | 2.235898899 | 0.755958409 | 1.13150623 | 0.196272163 | 0.73603183 | 0.692029776 | plasma_NFKBIA_lo |
| 287 | ERLEC1 | 0.183913381 | 0.188360924 | 0.095863112 | 0.153180784 | 0.160072955 | 0.185316581 | 0.174686112 | 0.187038048 | 0.18623206 | 0.241111424 | 0.196898435 | 0.234875054 | 0.307977463 | 0.124172909 | 0.181528965 | 0.949587326 | 1.25015811 | 0.266520633 | 0.261312163 | 0.051922902 | 0.162819128 | 0.148480866 | plasma_NFKBIA_lo |
| 288 | VIMP | 0.347988902 | 0.371538474 | 0.183796329 | 0.286746106 | 0.322279703 | 0.259296667 | 0.27624252 | 0.368278552 | 0.373846956 | 0.470622082 | 0.274063636 | 0.340976669 | 0.393329391 | 0.173375163 | 0.211265722 | 1.38262749 | 1.498162895 | 0.510946731 | 1.362084162 | 0.192516627 | 0.525613987 | 0.468451844 | plasma_NFKBIA_lo |
| 289 | KRTCAP2 | 0.459885852 | 0.562451643 | 0.140674089 | 0.485125634 | 0.465873618 | 0.480078785 | 0.466191578 | 0.486810349 | 0.647846876 | 0.483550729 | 0.531008503 | 0.518017202 | 0.508996645 | 0.314812751 | 0.362071961 | 1.160450531 | 1.49739141 | 0.570937015 | 0.664182156 | 0.075816496 | 0.428207685 | 0.455294674 | plasma_NFKBIA_lo |
| 290 | SELM | 0.176110726 | 0.166839761 | 0.062894521 | 0.211791484 | 0.201259422 | 0.254347899 | 0.20715839 | 0.152579968 | 0.196429897 | 0.10435537 | 0.257715806 | 0.170173324 | 0.063403449 | 0.113379848 | 0.039474745 | 0.791651711 | 1.06076628 | 0.329622021 | 0 | 0.024667689 | 0.07961872 | 0.124771983 | plasma_NFKBIA_lo |
| 291 | LMAN1 | 0.288086471 | 0.316686663 | 0.10191382 | 0.265482759 | 0.300117366 | 0.324476719 | 0.263868855 | 0.286719744 | 0.346198929 | 0.320640702 | 0.300984338 | 0.338206907 | 0.416832955 | 0.211688173 | 0.185055743 | 1.094576479 | 1.368872084 | 0.376448262 | 0.613479582 | 0.041078831 | 0.217825886 | 0.321127464 | plasma_NFKBIA_lo |
| 292 | SEL1L | 0.142296896 | 0.1573247 | 0.103877335 | 0.112342456 | 0.121319513 | 0.158541958 | 0.132291554 | 0.135253533 | 0.150056414 | 0.179148053 | 0.153147766 | 0.176913057 | 0.197523102 | 0.114183736 | 0.145467558 | 0.857907459 | 1.135272984 | 0.212442553 | 0.277216728 | 0.213305212 | 0.247814159 | 0.183979647 | plasma_NFKBIA_lo |
| 293 | PPIB | 0.926803041 | 1.131629852 | 0.194144848 | 0.89628432 | 0.918461507 | 0.801524022 | 0.793121879 | 0.898509154 | 1.099809253 | 1.023856175 | 0.91774088 | 1.041663753 | 1.063474033 | 0.618550258 | 0.868260304 | 1.772517779 | 2.041652502 | 1.113223724 | 1.904203824 | 0.150985417 | 1.204955121 | 1.348274781 | plasma_NFKBIA_lo |
| 294 | NUCB2 | 0.457311881 | 0.541977591 | 0.104792139 | 0.270630307 | 0.235857241 | 0.108599091 | 0.336547314 | 0.437285663 | 0.519375383 | 0.391088978 | 0.362715815 | 0.256759411 | 0.426698517 | 0.044071961 | 0.057493279 | 0.841125036 | 1.073328605 | 0.364200107 | 1.067777911 | 0.045558157 | 0.169211233 | 0.231589257 | plasma_NFKBIA_lo |
| 295 | RABAC1 | 0.522947267 | 0.658441644 | 0.107600319 | 0.465470829 | 0.521708738 | 0.472390264 | 0.414999935 | 0.620648142 | 0.692915505 | 0.596996817 | 0.353396975 | 0.566775042 | 0.67378636 | 0.418635461 | 0.773800347 | 1.293487904 | 1.595854953 | 0.675901446 | 0.72167091 | 0.080043064 | 0.422833607 | 0.493005946 | plasma_NFKBIA_lo |
| 296 | CRELD2 | 0.203698017 | 0.211070252 | 0.095645072 | 0.146837525 | 0.155063576 | 0.175243927 | 0.137082298 | 0.176342331 | 0.238374573 | 0.240689464 | 0.14490308 | 0.218585946 | 0.32562094 | 0.11945203 | 0.174029369 | 0.893314985 | 1.097308376 | 0.227448254 | 0.272380403 | 0.119442857 | 0.272184691 | 0.23434518 | plasma_NFKBIA_lo |
| 297 | PDIA6 | 0.361942694 | 0.468813528 | 0.096856134 | 0.305683767 | 0.345617817 | 0.421602301 | 0.309927068 | 0.37621287 | 0.461489709 | 0.514304206 | 0.42786427 | 0.505829928 | 0.891041888 | 0.356750784 | 0.682855689 | 1.13078579 | 1.469792966 | 0.497859591 | 0.880181946 | 0.099004712 | 0.708335436 | 0.826068696 | plasma_NFKBIA_lo |
| 298 | C19orf10 | 0.264567796 | 0.282956381 | 0.046260249 | 0.255565686 | 0.292158012 | 0.231425695 | 0.208072649 | 0.254338594 | 0.274090233 | 0.361049204 | 0.240366255 | 0.32359442 | 0.438710464 | 0.202947174 | 0.295088845 | 0.973386268 | 1.173112595 | 0.354931894 | 0.84908 | 0.073720647 | 0.589185837 | 0.625255805 | plasma_NFKBIA_lo |
| 299 | PDIA4 | 0.11868793 | 0.155183154 | 0.065249661 | 0.090296593 | 0.092700937 | 0.121447644 | 0.109547482 | 0.107512465 | 0.117557094 | 0.180192564 | 0.116451908 | 0.135595063 | 0.378045884 | 0.077037953 | 0.364139088 | 0.95638657 | 1.147200564 | 0.202568556 | 0.603672282 | 0.22406468 | 0.289233146 | 0.397555314 | plasma_NFKBIA_lo |
| 300 | HSPA5 | 1.158744569 | 0.784179834 | 0.881732952 | 0.907225156 | 0.969526488 | 0.73395138 | 0.502672265 | 1.088331774 | 0.884676905 | 1.21350926 | 0.717892762 | 1.07849945 | 0.870199458 | 0.551343394 | 0.743646941 | 1.814994127 | 1.79582193 | 1.059548231 | 1.413865248 | 0.965381289 | 1.250885571 | 0.950658494 | plasma_NFKBIA_lo |
| 301 | RPN2 | 0.225536751 | 0.331351632 | 0.091991123 | 0.222430291 | 0.239873528 | 0.281151151 | 0.236850374 | 0.219341424 | 0.308472284 | 0.292471803 | 0.275049101 | 0.293341627 | 0.407843348 | 0.189589256 | 0.299644172 | 0.775959037 | 1.079987159 | 0.355948563 | 0.717862993 | 0.07092397 | 0.435670427 | 0.576751226 | plasma_NFKBIA_lo |
| 302 | TPSAB1 | 0.000972628 | 0 | 0.002043145 | 0.003460272 | 0.001023191 | 0.00090971 | 0.005332859 | 0.002568378 | 0.003255415 | 0 | 0.001700176 | 0.002016665 | 0.019198822 | 0.001827012 | 0.001013662 | 0.0021546 | 0.002488537 | 2.08317521 | 0 | 0.010674498 | 0.004542548 | 0.011310392 | mast_cells |
| 303 | CPA3 | 0.000980116 | 0 | 0.001782786 | 0.003605314 | 0 | 0.001966527 | 0.002541668 | 0.001388299 | 0.000821395 | 0.001612488 | 0.001221278 | 0.001183161 | 0.013115774 | 0.000720825 | 0 | 0.001084144 | 0.000161586 | 1.727066821 | 0 | 0.005655473 | 0.003417459 | 0.01239523 | mast_cells |
| 304 | GATA2 | 0.000926783 | 0.003422434 | 0.005017312 | 0.004068005 | 0.002073284 | 0.001068061 | 0.000608347 | 0.000519958 | 0.002909131 | 0.003438779 | 0.000716114 | 0.001874649 | 0.010727179 | 0.000814352 | 0 | 0.002658329 | 0.002689241 | 1.239176642 | 0 | 0.007105647 | 0.009875208 | 0.00425725 | mast_cells |
| 305 | HPGDS | 0.000899884 | 0 | 0.004595611 | 0.003837604 | 0.00296574 | 0.003526725 | 0.001295537 | 0 | 0.001659971 | 0 | 0.009746272 | 0.000702865 | 0.010696893 | 0.000444379 | 0.012206265 | 0.00107317 | 0.000146563 | 1.139047298 | 0 | 0 | 0.001966865 | 0.024375608 | mast_cells |
| 306 | KIT | 0.001282375 | 0.000742504 | 0.007877432 | 0.019801378 | 0.024801686 | 0.002446757 | 0.005715135 | 0.002903822 | 0.00433819 | 0.025106872 | 0.002312298 | 0.117636343 | 0.017445556 | 0.001037263 | 0.002882262 | 0.007179389 | 0.002433293 | 1.184125035 | 0.029427759 | 0.001176812 | 0.007713636 | 0.006566434 | mast_cells |
| 307 | VWA5A | 0.008176453 | 0.009881998 | 0.013547067 | 0.008206349 | 0.002860301 | 0.024247329 | 0.008229126 | 0.004654398 | 0.007756552 | 0.01755162 | 0.016075611 | 0.008288954 | 0.016273981 | 0.004021796 | 0.001863137 | 0.023990185 | 0.031530956 | 0.855547475 | 0.019588853 | 0.006476725 | 0.011458766 | 0.015985283 | mast_cells |
| 308 | HDC | 0.000461345 | 0.004351702 | 0.005892187 | 0.00229598 | 0.001844453 | 0.000999439 | 0.001599263 | 0.001431305 | 0.000714022 | 0.004319672 | 0.001997277 | 0.000878466 | 0.00757743 | 0.000725156 | 0.000592512 | 0.000835943 | 0.001742462 | 0.892628277 | 0 | 0 | 0.001709711 | 0.002908641 | mast_cells |
| 309 | CLU | 0.030027646 | 0.030577931 | 0.015158782 | 0.021473767 | 0.012195285 | 0.002898077 | 0.018092525 | 0.029804325 | 0.042161697 | 0.032112127 | 0.015884177 | 0.008340091 | 0.01126252 | 0.000457058 | 0.01081559 | 0.006426486 | 0.000786965 | 0.985589765 | 0.007018521 | 0.015545957 | 0.009948088 | 0.005561548 | mast_cells |
| 310 | LTC4S | 0.006075054 | 0.007978499 | 0.001889472 | 0.018491472 | 0.012947234 | 0.002796634 | 0.014063653 | 0.014130843 | 0.013629009 | 0.018296138 | 0.008571531 | 0.009745887 | 0.019961819 | 0.005602356 | 0.005623536 | 0.099504394 | 0.165678979 | 1.215383702 | 0.021341906 | 0.005930155 | 0.020841421 | 0.03101694 | mast_cells |
| 311 | IL1RL1 | 0.000961169 | 0.000677573 | 0.003334173 | 0.002558431 | 0.003352446 | 0.020005808 | 0.001743978 | 0.001418886 | 0.001490744 | 0.002135812 | 0.002313403 | 0.008522343 | 0.01347149 | 0.000551172 | 0 | 0.002920368 | 0.000269126 | 0.79486764 | 0 | 0.007913882 | 0.000946863 | 0.005209359 | mast_cells |
| 312 | SLC18A2 | 0.000407402 | 0 | 0.001331931 | 0.002329709 | 0.001665951 | 0.001807685 | 0.001934599 | 0.000432689 | 0.001289976 | 0 | 0.002072852 | 0.002031281 | 0.005522841 | 0.001005962 | 0 | 0.00068203 | 0 | 0.744590217 | 0.009420133 | 0 | 0.000167587 | 0 | mast_cells |
| 313 | MS4A2 | 0 | 0 | 0 | 0.000923984 | 0.000363291 | 0.000240577 | 0.002652262 | 0 | 0.00188465 | 0 | 0.000310619 | 0.000526289 | 0.00320774 | 0.000103395 | 0 | 0.000230357 | 0 | 0.788943168 | 0 | 0 | 0.000203784 | 0 | mast_cells |
| 314 | CTSG | 0 | 0.000776934 | 0 | 0.001138813 | 0 | 0 | 0.001896715 | 0 | 0.001800714 | 0 | 0.000200172 | 0.001219213 | 0.01218749 | 8.85E-05 | 0 | 0.000436497 | 0.000787465 | 0.99507068 | 0.00779332 | 0.003354622 | 0.001006351 | 0.005209359 | mast_cells |
| 315 | HPGD | 0.028573791 | 0.023417039 | 0.020174615 | 0.031610015 | 0.042267669 | 0.160608642 | 0.021916604 | 0.037620637 | 0.029325989 | 0.060697653 | 0.008426432 | 0.14144682 | 0.090614068 | 0.00285483 | 0.004987469 | 0.052908808 | 0.031485362 | 0.604309553 | 0 | 0.007931197 | 0.013566593 | 0.008845719 | mast_cells |
| 316 | CD9 | 0.02724513 | 0.011279775 | 0.010527854 | 0.04011589 | 0.023846523 | 0.013626597 | 0.030887609 | 0.177820044 | 0.228244043 | 0.114705725 | 0.012328228 | 0.04219525 | 0.127930316 | 0.020347788 | 0.11025459 | 0.080938567 | 0.101737217 | 1.204795878 | 0.361279392 | 0.03700048 | 0.197044159 | 0.44009804 | mast_cells |
| 317 | CD69 | 3.124821488 | 1.748673389 | 2.411466955 | 3.085595411 | 2.948524444 | 1.966765982 | 1.159139198 | 3.520311692 | 2.501003148 | 2.897344837 | 2.128867515 | 3.110006161 | 1.151327438 | 2.742089877 | 0.870841382 | 1.239955771 | 0.324718312 | 1.805049731 | 0.636213737 | 0.509243996 | 0.52945327 | 0.344008692 | mast_cells |
| 318 | KLRB1 | 0.601721127 | 0.512640093 | 0.494465185 | 1.92151749 | 1.990128964 | 1.278552465 | 1.400714149 | 1.598093905 | 1.486566605 | 1.660757303 | 1.989555854 | 2.800629206 | 0.465564824 | 0.050230579 | 0.05273308 | 0.494102053 | 0.066157416 | 0.709269418 | 0.035597069 | 0.064232721 | 0.110007601 | 0.095693587 | mast_cells |
| 319 | IGFBP7 | 0.019233412 | 0.043915597 | 0.009074262 | 0.002619568 | 0.00184722 | 0.002606338 | 0.004743969 | 0.001824818 | 0.003989371 | 0.098001953 | 0.003122659 | 0.002091574 | 0.013098548 | 0.001631757 | 0.003244377 | 0.005105874 | 0.001990398 | 0.633907619 | 0.033096383 | 0.057403302 | 0.108891451 | 0.038909526 | mast_cells |
| 320 | CLIC3 | 0.170058397 | 0.254285698 | 0.02902829 | 0.055522291 | 0.069731272 | 0.027590636 | 0.053687536 | 0.182355238 | 0.23698312 | 0.59128367 | 0.011062484 | 0.137516103 | 0.040814687 | 0.04163444 | 0.007404804 | 0.040531805 | 0.03337153 | 0.042345603 | 1.546863616 | 0.00179842 | 0.0185153 | 0.021466414 | pdcs |
| 321 | SCT | 0 | 0 | 0 | 0.000442935 | 0 | 0 | 0 | 0.000674218 | 0.000796576 | 0.002110482 | 0 | 0 | 0.0007187 | 0.000599211 | 0.000901013 | 0.000757322 | 0.000360815 | 0.019385774 | 1.265236596 | 0 | 0.005152321 | 0.001431857 | pdcs |
| 322 | SMPD3 | 0.017216154 | 0.018374579 | 0.002616243 | 0.004880128 | 0.003861426 | 0.02868842 | 0.011101398 | 0.019294729 | 0.022896333 | 0.034914169 | 0.038190474 | 0.005933921 | 0.012935119 | 0.005862534 | 0.004954868 | 0.027315481 | 0.031587545 | 0.003933318 | 1.253524455 | 0.003271172 | 0.010070724 | 0 | pdcs |
| 323 | LILRA4 | 0.000386366 | 0 | 0.001784246 | 0.000181937 | 0 | 0 | 0 | 0.000565826 | 0 | 0 | 0 | 0.000233798 | 0.001894652 | 0.010621167 | 0.00367256 | 0.001014342 | 0.000793065 | 0.003184183 | 1.260350279 | 0.0022558 | 0.01563618 | 0.008407176 | pdcs |
| 324 | GZMB | 0.798936162 | 0.885286936 | 0.04678006 | 0.023655516 | 0.032841324 | 0.05027632 | 0.016825547 | 0.306070315 | 0.191183445 | 1.189253705 | 0.029995903 | 0.387404491 | 0.21100377 | 0.007749313 | 0.013109023 | 0.069884402 | 0.025642766 | 0.018545587 | 3.340824722 | 0.013838932 | 0.062185815 | 0.034023671 | pdcs |
| 325 | TSPAN13 | 0.005407762 | 0.002742955 | 0.019542865 | 0.004766192 | 0.005074981 | 0.021841245 | 0.001277051 | 0.00355098 | 0.004142759 | 0.025868742 | 0.009026848 | 0.009542072 | 0.196942175 | 0.354122136 | 0.368001914 | 0.127737339 | 0.156873895 | 0.054157413 | 1.738130353 | 0.010170429 | 0.0426498 | 0.042777822 | pdcs |
| 326 | EGLN3 | 0.002570445 | 0.000498813 | 0.027948364 | 0.018115339 | 0.02182042 | 0.008892353 | 0.014620717 | 0.006518318 | 0.004694222 | 0.007766149 | 0.029803012 | 0.034067225 | 0.007510105 | 0.005702545 | 0.009175724 | 0.003618206 | 0.000719098 | 0.042528395 | 1.110833291 | 0.032539981 | 0.0174122 | 0.024687686 | pdcs |
| 327 | C12orf75 | 0.414623306 | 0.56123923 | 0.020814317 | 0.122042091 | 0.208024356 | 0.132825531 | 0.078548403 | 0.226494192 | 0.214670789 | 0.262396619 | 0.060639122 | 0.261171708 | 0.888302173 | 0.148276682 | 0.475733478 | 0.097117343 | 0.046507153 | 0.080216083 | 1.686039 | 0.019748967 | 0.077462505 | 0.050109445 | pdcs |
| 328 | SERPINF1 | 0.000233602 | 0.001824283 | 0.001103605 | 0.002159054 | 0.001930482 | 0.000813281 | 0.003316521 | 0.003520349 | 0.009101391 | 0 | 0.000552137 | 0.002897876 | 0.004373002 | 0.005825347 | 0.011173415 | 0.004363488 | 0.004740504 | 0.186535224 | 1.310537804 | 0.005543727 | 0.052197334 | 0.220547827 | pdcs |
| 329 | PLD4 | 0.000838424 | 0 | 0.00577374 | 0.000485604 | 0.000396602 | 0.000822508 | 0.000821137 | 0.000478535 | 0.001203112 | 0 | 0 | 0.000427369 | 0.018816566 | 0.08252475 | 0.080878809 | 0.00644305 | 0.008355185 | 0.003722696 | 1.351385325 | 0.01503004 | 0.149224182 | 0.229456452 | pdcs |
| 330 | BCL11A | 0.004384159 | 0.007162552 | 0.026170367 | 0.003468514 | 0.002540014 | 0.006438438 | 0.005785295 | 0.004821559 | 0.002736851 | 0.002311142 | 0.005062199 | 0.0054234 | 0.3404153 | 0.610353809 | 0.853982446 | 0.015960926 | 0.013804659 | 0.039256758 | 1.329015057 | 0.056884665 | 0.176719398 | 0.112952496 | pdcs |
| 331 | LINC00996 | 0.02186605 | 0.03108302 | 0.01724146 | 0.018660603 | 0.010408695 | 0.003675985 | 0.010665551 | 0.023881112 | 0.027896098 | 0.354030686 | 0.007106173 | 0.032725026 | 0.013398588 | 0.068897316 | 0.004798522 | 0.058069945 | 0.064136458 | 0.177747281 | 1.252274727 | 0.06910741 | 0.055818312 | 0.273142876 | pdcs |
| 332 | TCF4 | 0.010942239 | 0.014072183 | 0.045914826 | 0.01144475 | 0.00645506 | 0.025059651 | 0.01325133 | 0.007983938 | 0.010616891 | 0.052849601 | 0.019291477 | 0.011259308 | 0.552422872 | 0.762860893 | 0.860279337 | 0.566053413 | 0.485887954 | 0.40766126 | 2.563655132 | 0.133879679 | 0.229579432 | 0.221351396 | pdcs |
| 333 | ITM2C | 0.387897411 | 0.391216556 | 0.056057225 | 0.116133761 | 0.106484358 | 0.088728945 | 0.104288175 | 0.422942326 | 0.477005936 | 0.413227572 | 0.083956174 | 0.243854697 | 0.214903593 | 0.140215526 | 0.021134414 | 0.985538302 | 1.292472067 | 0.604724959 | 1.70084007 | 0.038172309 | 0.087280723 | 0.198546477 | pdcs |
| 334 | PTGDS | 0.008209364 | 0.03810652 | 0.006106687 | 0.001789951 | 0.006232645 | 0.001592873 | 0.003390809 | 0.001340483 | 0.000756169 | 0.016998406 | 0.001687477 | 0.001837582 | 0.005677583 | 0.001112002 | 0.002290116 | 0.00172534 | 0.001495852 | 0.159450667 | 1.088487505 | 0.009478 | 0.01151974 | 0.095365007 | pdcs |
| 335 | UGCG | 0.161533158 | 0.118617054 | 0.11905358 | 0.157116686 | 0.159911069 | 0.346613197 | 0.150185371 | 0.115491738 | 0.128170066 | 0.14472284 | 0.301006951 | 0.150026513 | 0.38745393 | 0.377124759 | 0.62645012 | 0.162741564 | 0.106203917 | 0.243915124 | 1.835549979 | 0.059389385 | 0.311720339 | 0.188172132 | pdcs |
| 336 | IL3RA | 0.001677981 | 0.003977193 | 0.003195462 | 0.006485956 | 0.00289291 | 0.005635483 | 0.00566559 | 0.000853713 | 0.004722215 | 0.003859363 | 0.002693531 | 0.003838863 | 0.003299898 | 0.010326265 | 0.008500282 | 0.007363179 | 0.008274963 | 0.119618083 | 1.596864844 | 0.161255246 | 0.298991668 | 0.141908371 | pdcs |
| 337 | ALOX5AP | 0.442840492 | 0.383725007 | 0.063633163 | 0.39696278 | 0.395258364 | 0.263371873 | 0.352250967 | 0.592580072 | 0.7045309 | 0.531004365 | 0.598664614 | 0.571226309 | 0.477179687 | 0.300567028 | 0.894279684 | 0.143886522 | 0.035049457 | 0.760656656 | 1.996019988 | 0.047768701 | 0.218409116 | 0.182635618 | pdcs |
| 338 | CCDC50 | 0.031524528 | 0.039704743 | 0.065302281 | 0.058021995 | 0.056482782 | 0.154382444 | 0.073302208 | 0.067414043 | 0.057437829 | 0.167396792 | 0.239349805 | 0.084164802 | 0.284612408 | 0.586959267 | 0.221675878 | 0.360356483 | 0.370596546 | 0.159284624 | 1.922555871 | 0.075279006 | 0.29308878 | 0.280293224 | pdcs |
| 339 | IRF4 | 0.29501949 | 0.110224944 | 0.354427576 | 0.158008405 | 0.220354661 | 0.397998876 | 0.080073692 | 0.266251036 | 0.131379742 | 0.211674099 | 0.269929201 | 0.433349896 | 0.285050919 | 0.464551457 | 0.155945033 | 0.753629717 | 0.73372119 | 0.104128142 | 1.695146565 | 0.11092118 | 0.291251067 | 0.072995594 | pdcs |
| 340 | PLAC8 | 0.211983776 | 0.425909724 | 0.098608376 | 0.470257697 | 0.420009154 | 0.027140968 | 0.426553011 | 0.193020275 | 0.31471107 | 0.528921392 | 0.132835429 | 0.154208874 | 0.061980217 | 0.756829892 | 0.054163142 | 0.073287568 | 0.039466952 | 0.137275009 | 2.438964988 | 0.133694478 | 0.721212318 | 0.456287946 | pdcs |
| 341 | PPP1R14B | 0.304625158 | 0.32071122 | 0.048044304 | 0.307118766 | 0.33029498 | 0.145513624 | 0.242586192 | 0.369912579 | 0.448626961 | 0.44434962 | 0.213560295 | 0.322857412 | 0.652178012 | 0.475094767 | 0.519020292 | 0.345774989 | 0.307854401 | 0.368953643 | 2.40073795 | 0.059540087 | 0.501980045 | 0.517481231 | pdcs |
| 342 | IRF7 | 0.17589797 | 0.172530144 | 0.169904356 | 0.137334724 | 0.147248705 | 0.188026916 | 0.111396622 | 0.168860642 | 0.157864777 | 0.205679471 | 0.149744473 | 0.200299665 | 0.158455222 | 0.183182157 | 0.268072355 | 0.286092586 | 0.257193454 | 0.13636416 | 2.159579497 | 0.571683833 | 0.429816312 | 0.161111846 | pdcs |
| 343 | SEC61B | 0.869633672 | 0.925642007 | 0.293153358 | 0.82572808 | 0.973559661 | 0.872620957 | 0.697976054 | 0.922462835 | 0.954254432 | 1.025093975 | 0.872374725 | 1.064232025 | 1.415225261 | 0.610564367 | 1.22923627 | 1.997016356 | 2.222099188 | 1.103748215 | 2.805592798 | 0.21462644 | 1.329936613 | 1.155472194 | pdcs |
| 344 | DUSP5 | 0.63762527 | 0.219166378 | 0.477983679 | 0.472786233 | 0.749226174 | 0.403300269 | 0.087343001 | 0.643105632 | 0.270685482 | 0.498756905 | 0.458142494 | 0.995684387 | 0.319334859 | 0.284896891 | 0.190678139 | 0.937247594 | 0.299482599 | 0.216506154 | 2.157129869 | 0.533078286 | 0.582990434 | 0.189509627 | pdcs |
| 345 | RAB11FIP1 | 0.37323736 | 0.235351387 | 0.646153973 | 0.493673551 | 0.643469937 | 1.02379642 | 0.225023373 | 0.594250199 | 0.387157579 | 0.783187106 | 0.974122715 | 0.85857997 | 0.889570734 | 0.942012424 | 1.102366112 | 0.580413615 | 0.142131094 | 0.38244821 | 2.34899632 | 0.509655108 | 0.948031344 | 0.44300055 | pdcs |
| 346 | IRF8 | 0.113689532 | 0.044009136 | 0.077090548 | 0.04362911 | 0.045612693 | 0.032164305 | 0.026933189 | 0.119642128 | 0.070672804 | 0.472806489 | 0.040230946 | 0.067281582 | 0.768818864 | 0.814531558 | 1.512329 | 0.070847861 | 0.027378684 | 0.077411306 | 2.241781516 | 0.692878869 | 0.821458296 | 0.623240479 | pdcs |
| 347 | LDLRAD4 | 0.209095421 | 0.126225144 | 0.417953005 | 0.33285001 | 0.344745673 | 0.430966762 | 0.231635662 | 0.945771267 | 0.498603113 | 0.596109521 | 0.394889778 | 0.570560413 | 0.262230919 | 0.334676635 | 0.182920495 | 0.149539693 | 0.022617834 | 0.243506112 | 1.670996278 | 0.301392143 | 0.386825903 | 0.107403204 | pdcs |
| 348 | LTB | 0.744378016 | 0.878877217 | 0.772604044 | 2.07436238 | 2.188194398 | 2.173513811 | 2.047322817 | 1.144157907 | 1.579337964 | 0.629888746 | 1.892663652 | 2.154613865 | 1.475949381 | 2.103917964 | 2.710895204 | 0.462672623 | 0.100548419 | 0.749828473 | 1.760454361 | 0.166009178 | 0.381877431 | 0.436250967 | pdcs |
| 349 | CHML | 0.044862362 | 0.041124455 | 0.049549794 | 0.035744163 | 0.023174938 | 0.028614102 | 0.035746808 | 0.042584758 | 0.053680931 | 0.045890254 | 0.020351445 | 0.028781515 | 0.052366624 | 0.154217617 | 0.046490613 | 0.068948099 | 0.041856243 | 0.066591482 | 1.184294631 | 0.041554333 | 0.312433126 | 0.144128697 | pdcs |
| 350 | CXCR4 | 1.908729262 | 0.917220735 | 0.854485692 | 1.3001875 | 1.765067937 | 0.96203251 | 0.582084129 | 1.401309649 | 0.777636477 | 1.252052627 | 1.217606613 | 1.04346562 | 1.058100396 | 2.021505286 | 0.843239483 | 0.582094539 | 0.16707254 | 0.647022727 | 2.783040666 | 0.513035308 | 1.369051849 | 0.619729731 | pdcs |
| 351 | RGS1 | 1.510014898 | 0.77468985 | 1.651347015 | 1.444606609 | 1.467206563 | 2.505842969 | 0.551271089 | 1.749827483 | 1.08857103 | 1.571681508 | 1.406864697 | 2.052486676 | 1.266615722 | 0.694353746 | 1.27333071 | 2.253584734 | 0.801204831 | 1.798908547 | 3.155938484 | 0.848405141 | 1.288677631 | 1.099100664 | pdcs |
| 352 | 7SK.2 | 0.41307766 | 0.637378957 | 1.792163904 | 0.495019652 | 0.335190829 | 0.498805267 | 0.740339105 | 0.388116259 | 0.558208072 | 0.59346796 | 0.500547073 | 0.461836349 | 0.797274264 | 0.756776662 | 1.237460931 | 0.780773936 | 0.681681222 | 0.592732967 | 0.592280596 | 3.231276566 | 0.502389701 | 0.369306673 | SOX4/MAFA |
| 353 | METTL12 | 0.199247919 | 0.218593187 | 0.682364699 | 0.189271901 | 0.157158523 | 0.137306672 | 0.233841583 | 0.128633341 | 0.195053794 | 0.203596374 | 0.167177484 | 0.150398008 | 0.209219842 | 0.153238931 | 0.19493353 | 0.202857909 | 0.150704335 | 0.146290319 | 0.120125136 | 1.825489452 | 0.109536443 | 0.071367889 | SOX4/MAFA |
| 354 | HIST1H2AL | 0.036335946 | 0.057766981 | 0.21765095 | 0.027725948 | 0.014929218 | 0.038254236 | 0.042138574 | 0.014471763 | 0.030042293 | 0.032987013 | 0.049631052 | 0.02239937 | 0.396083017 | 0.045011612 | 0.078151319 | 0.12658604 | 0.092434052 | 0.03236347 | 0.063598109 | 1.148560429 | 0.031333739 | 0.02536297 | SOX4/MAFA |
| 355 | RP11-231D20.2 | 0.011335679 | 0.028142134 | 0.148345109 | 0.023779629 | 0.002689383 | 0.012896753 | 0.028408385 | 0.003466462 | 0.023431801 | 0.022116423 | 0.008079604 | 0.008942994 | 0.011705996 | 0.009988582 | 0.015345619 | 0.010486089 | 0.00661392 | 0.007030962 | 0.00818725 | 0.850653638 | 0.019389899 | 0.005350185 | SOX4/MAFA |
| 356 | HEXIM1 | 0.371241424 | 0.306089296 | 1.153907235 | 0.291807629 | 0.311296416 | 0.422452169 | 0.252660137 | 0.287729962 | 0.338340075 | 0.417063735 | 0.293990482 | 0.369561468 | 0.20340598 | 0.191642213 | 0.165829877 | 0.363030093 | 0.221780074 | 0.424260155 | 0.370880032 | 1.976469967 | 0.245728376 | 0.181478368 | SOX4/MAFA |
| 357 | MAFA | 0.011299853 | 0.009020617 | 0.060045688 | 0.011389725 | 0.016918271 | 0.010431521 | 0.006301718 | 0.015927729 | 0.012734636 | 0.008232973 | 0.007717414 | 0.013470266 | 0.030338538 | 0.013244911 | 0.078048483 | 0.025178677 | 0.009814546 | 0.013772692 | 0.016459468 | 0.807703988 | 0.031836887 | 0.011744354 | SOX4/MAFA |
| 358 | HIST1H3A | 0.050865568 | 0.079454561 | 0.333128262 | 0.045355337 | 0.022231443 | 0.039749958 | 0.088702531 | 0.02203204 | 0.055565545 | 0.05649574 | 0.046281794 | 0.033622407 | 0.05043028 | 0.02844935 | 0.033454792 | 0.091409918 | 0.077778827 | 0.041142318 | 0.068736965 | 0.793042681 | 0.025236958 | 0.016547172 | SOX4/MAFA |
| 359 | IER5L | 0.435844468 | 0.247913913 | 0.648990781 | 0.279643746 | 0.146093533 | 0.286198388 | 0.154969757 | 0.342345606 | 0.378566181 | 0.525372829 | 0.239528699 | 0.323106715 | 0.180356602 | 0.169468384 | 0.152768779 | 0.194745174 | 0.080070686 | 0.325813475 | 0.057659762 | 1.654578028 | 0.270547569 | 0.277287439 | SOX4/MAFA |
| 360 | MED29 | 0.265532033 | 0.258557731 | 0.489464969 | 0.198032981 | 0.261139032 | 0.254828686 | 0.203298877 | 0.321527988 | 0.268139656 | 0.272269195 | 0.216621271 | 0.237669481 | 0.23328899 | 0.17640372 | 0.262340996 | 0.470517142 | 0.291580719 | 0.205726189 | 0.382411733 | 1.704614556 | 0.479825154 | 0.265542471 | SOX4/MAFA |
| 361 | PPP1R10 | 0.88956005 | 0.911930654 | 2.043648174 | 0.964801581 | 0.68321205 | 0.902921328 | 0.889267728 | 0.794226119 | 0.970357672 | 0.961704874 | 0.732780166 | 0.844677671 | 0.593310295 | 0.674632896 | 0.669908473 | 1.235555128 | 0.74592359 | 0.92045604 | 1.152407209 | 1.946558737 | 0.627146752 | 0.365971659 | SOX4/MAFA |
| 362 | RP11-796E2.4 | 0.224108519 | 0.309249463 | 0.631288768 | 0.310597731 | 0.182459957 | 0.30868371 | 0.3604191 | 0.15294819 | 0.273505206 | 0.267443148 | 0.260535944 | 0.213270663 | 0.303946733 | 0.339595154 | 0.410712206 | 0.200566461 | 0.129149691 | 0.134903746 | 0.016783641 | 1.220586514 | 0.208185849 | 0.122074039 | SOX4/MAFA |
| 363 | PLK2 | 0.034346225 | 0.023092887 | 0.2891787 | 0.116466562 | 0.070103536 | 0.042668815 | 0.062043305 | 0.047137467 | 0.063898898 | 0.155496232 | 0.097298395 | 0.067735099 | 0.056391441 | 0.086997641 | 0.09659715 | 0.09885011 | 0.061032367 | 0.242994789 | 0.075927613 | 1.166096458 | 0.125171132 | 0.075119019 | SOX4/MAFA |
| 364 | HIST1H2AC | 0.106071604 | 0.207191286 | 0.691409196 | 0.124866015 | 0.086561645 | 0.159218497 | 0.32596211 | 0.09069007 | 0.192611884 | 0.194289856 | 0.191814751 | 0.1567179 | 0.408332781 | 0.248195692 | 0.193611749 | 0.812523995 | 0.620435065 | 0.239801728 | 0.419196413 | 1.215631238 | 0.12253549 | 0.113015298 | SOX4/MAFA |
| 365 | RP5-821D11.7 | 0.106516699 | 0.169861121 | 0.497273749 | 0.156822608 | 0.076574942 | 0.147495715 | 0.162899161 | 0.09321332 | 0.221552321 | 0.12706827 | 0.148481378 | 0.131803064 | 0.190354219 | 0.226468243 | 0.247208767 | 0.281993244 | 0.151675806 | 0.061036658 | 0.140752137 | 0.923946998 | 0.105538228 | 0.085158713 | SOX4/MAFA |
| 366 | ILF3-AS1 | 0.266287705 | 0.291011897 | 0.572965342 | 0.351292198 | 0.197335832 | 0.244914456 | 0.318141723 | 0.198878889 | 0.324516155 | 0.265396666 | 0.27520026 | 0.192484806 | 0.28097806 | 0.388380297 | 0.326930264 | 0.302746707 | 0.19443896 | 0.191811435 | 0.337669305 | 1.468865879 | 0.359759597 | 0.314067997 | SOX4/MAFA |
| 367 | CTD-2562J15.6 | 0.101399454 | 0.122943338 | 0.216854645 | 0.109086491 | 0.067723743 | 0.035520849 | 0.086994149 | 0.021879662 | 0.059211122 | 0.094244756 | 0.039309326 | 0.03151651 | 0.030722111 | 0.121791016 | 0.030789023 | 0.173729373 | 0.07829788 | 0.087469155 | 0.038420872 | 0.742982016 | 0.058397458 | 0.036352633 | SOX4/MAFA |
| 368 | RNF213 | 1.323968892 | 1.620108307 | 1.366968333 | 1.014947913 | 0.954925018 | 1.385974593 | 1.199862929 | 1.053609841 | 1.188322738 | 1.177937662 | 0.974831636 | 1.3508573 | 0.582667074 | 0.558239198 | 0.311229249 | 0.73737875 | 0.487212728 | 0.547821037 | 0.930705295 | 1.964244057 | 0.963522227 | 0.741620055 | SOX4/MAFA |
| 369 | MAFB | 0.014988865 | 0.013493321 | 0.02489308 | 0.008989799 | 0.003458267 | 0.008437417 | 0.002547114 | 0.00498849 | 0.004384526 | 0.008400427 | 0.009287718 | 0.00727489 | 0.045814816 | 0.004616135 | 0.029290407 | 0.015804119 | 0.009090495 | 0.059545239 | 0.006489826 | 1.86138327 | 0.893818328 | 0.884799824 | SOX4/MAFA |
| 370 | AMD1 | 1.273475996 | 0.745942625 | 1.329550433 | 0.895103681 | 1.026460904 | 1.034873802 | 0.553132844 | 1.118166912 | 0.818103556 | 1.010396063 | 0.78205531 | 1.038248307 | 0.699319344 | 0.641286577 | 0.782854125 | 0.639974133 | 0.352068979 | 0.766573007 | 0.48031169 | 1.788453635 | 0.751780385 | 0.572789037 | SOX4/MAFA |
| 371 | RP11-727F15.9 | 0.051892262 | 0.091611549 | 0.200502495 | 0.066634902 | 0.027429679 | 0.047886775 | 0.083231793 | 0.036646474 | 0.108168455 | 0.063713905 | 0.055267058 | 0.036696963 | 0.080190923 | 0.07686606 | 0.068401403 | 0.388037832 | 0.239920185 | 0.081680798 | 0.023607733 | 0.690627647 | 0.059340296 | 0.036412416 | SOX4/MAFA |
| 372 | CTA-29F11.1 | 0.276657443 | 0.247485821 | 0.543241477 | 0.324305914 | 0.238478205 | 0.177333699 | 0.205887194 | 0.18973007 | 0.263180265 | 0.221577945 | 0.196249834 | 0.228616892 | 0.218045353 | 0.13026165 | 0.174373853 | 0.24045861 | 0.144819611 | 0.206113061 | 0.21548462 | 1.305542816 | 0.324442873 | 0.165896213 | SOX4/MAFA |
| 373 | SOX4 | 0.029017621 | 0.027632548 | 0.145686003 | 0.031561345 | 0.017645165 | 0.16264686 | 0.04593494 | 0.028954625 | 0.041577977 | 0.126667503 | 0.056441868 | 0.065553906 | 0.242900558 | 0.163601694 | 0.326829 | 0.180361713 | 0.085979703 | 0.722644988 | 2.221179209 | 1.726083503 | 0.371424604 | 0.225572673 | SOX4/MAFA |
| 374 | RPPH1 | 0.087370373 | 0.084596101 | 0.374278183 | 0.062177182 | 0.049200847 | 0.046025572 | 0.057268569 | 0.058869239 | 0.071859493 | 0.08816512 | 0.05829214 | 0.053663552 | 0.101600486 | 0.071848746 | 0.076414364 | 0.232539684 | 0.099974093 | 0.071376117 | 0.491633836 | 1.013041294 | 0.171285726 | 0.085413392 | SOX4/MAFA |
| 375 | HIST1H2AG | 0.064593934 | 0.122993537 | 0.468275989 | 0.079179953 | 0.043048488 | 0.054140225 | 0.142224397 | 0.048670718 | 0.130270948 | 0.071421412 | 0.071146549 | 0.048287432 | 0.420084444 | 0.060606503 | 0.060493149 | 0.4275468 | 0.360070801 | 0.053594603 | 0.057205857 | 0.5410243 | 0.022333191 | 0.032945415 | SOX4/MAFA |
| 376 | LINC00936 | 0.255450981 | 0.284239877 | 0.708029555 | 0.420641313 | 0.268408965 | 0.246029229 | 0.337573784 | 0.250668776 | 0.365934117 | 0.237262355 | 0.353192979 | 0.256656341 | 0.343113249 | 0.393189925 | 0.497258632 | 0.569487603 | 0.261841453 | 0.213883791 | 0.215076096 | 1.751869444 | 0.965576499 | 0.450114535 | SOX4/MAFA |
| 377 | HES1 | 0.013490321 | 0.009856947 | 0.115611462 | 0.036160612 | 0.019190505 | 0.116173943 | 0.017766564 | 0.016294531 | 0.017866694 | 0.032446571 | 0.073246269 | 0.028491995 | 0.060817753 | 0.362651967 | 0.158000102 | 0.044268214 | 0.019066854 | 0.638805574 | 0.085036198 | 1.758801915 | 0.614892743 | 0.614964135 | SOX4/MAFA |
| 378 | WDR74 | 0.155159829 | 0.209079886 | 0.629957644 | 0.201584 | 0.173077557 | 0.238126688 | 0.260019016 | 0.191486411 | 0.210286022 | 0.305892125 | 0.24464817 | 0.183417713 | 0.250307525 | 0.274461326 | 0.286748815 | 0.588690134 | 0.348824279 | 0.337906841 | 0.57574243 | 1.312993755 | 0.314362734 | 0.227357204 | SOX4/MAFA |
| 379 | RNU12 | 0.073898972 | 0.100259434 | 0.519356589 | 0.082823194 | 0.038360996 | 0.060350348 | 0.099747578 | 0.046704839 | 0.087522557 | 0.099275726 | 0.061484554 | 0.079216415 | 0.102145785 | 0.052379205 | 0.081738576 | 0.500031084 | 0.301914955 | 0.125226379 | 0.351690744 | 0.825933233 | 0.095903772 | 0.059377415 | SOX4/MAFA |
| 380 | EXT1 | 0.024185024 | 0.043125264 | 0.210758327 | 0.045113182 | 0.0279857 | 0.067033751 | 0.067964865 | 0.032799971 | 0.051746152 | 0.065242196 | 0.053326043 | 0.0455353 | 0.108571679 | 0.02387746 | 0.173325036 | 0.102765611 | 0.076926889 | 0.148231931 | 0.071388915 | 0.748235404 | 0.103345048 | 0.065263959 | SOX4/MAFA |
| 381 | GLS | 1.07141225 | 0.828908396 | 1.407141945 | 0.880306409 | 0.932193527 | 1.113425803 | 0.725210665 | 1.146785778 | 0.881696372 | 0.975227701 | 0.903989294 | 1.082495254 | 0.807975222 | 0.685543923 | 0.92525734 | 1.00851781 | 0.650926555 | 0.67376337 | 0.696491245 | 1.574044853 | 0.814771031 | 0.413206009 | SOX4/MAFA |
| 382 | GADD45G | 0.075773491 | 0.076722477 | 0.550075826 | 0.296671247 | 0.184554509 | 0.511083259 | 0.227857158 | 0.111852473 | 0.181377655 | 0.223161196 | 0.610170533 | 0.475295307 | 0.195556574 | 0.070948454 | 0.152852576 | 0.162563282 | 0.045901533 | 0.271594413 | 0.060561485 | 0.916111361 | 0.175993404 | 0.253387285 | SOX4/MAFA |
| 383 | RP1-313I6.12 | 0.16408294 | 0.101602255 | 0.294802446 | 0.286256023 | 0.298633736 | 0.119234574 | 0.121219433 | 0.308547171 | 0.196263315 | 0.21448183 | 0.139519246 | 0.183893272 | 0.134544046 | 0.427446352 | 0.179831536 | 0.313802764 | 0.090926945 | 0.223209859 | 0.567720253 | 0.915729369 | 0.15905324 | 0.067058191 | SOX4/MAFA |
| 384 | YME1L1 | 0.749866219 | 0.675585029 | 1.593808674 | 0.859769242 | 0.733613449 | 0.815937529 | 0.658744249 | 0.765406508 | 0.823899502 | 0.773416177 | 0.80022284 | 0.759382181 | 0.673532966 | 0.643611386 | 0.653229024 | 0.730567663 | 0.490088884 | 0.6428802 | 0.880456049 | 1.56103684 | 0.675794161 | 0.39250079 | SOX4/MAFA |
| 385 | INTS6 | 0.595385708 | 0.450760425 | 0.969091481 | 0.591498499 | 0.549346392 | 0.458165292 | 0.335142271 | 0.571256071 | 0.58754604 | 0.592099848 | 0.378110543 | 0.627108895 | 0.370896539 | 0.415561714 | 0.417324341 | 0.666011049 | 0.317519519 | 0.661162353 | 0.592476059 | 1.308641928 | 0.402022883 | 0.199751413 | SOX4/MAFA |
| 386 | SNORD3A | 0.032284265 | 0.059532334 | 0.313997439 | 0.050858678 | 0.01386993 | 0.032622926 | 0.078566893 | 0.031439149 | 0.069312334 | 0.099725641 | 0.028028305 | 0.057902116 | 0.089698054 | 0.04396139 | 0.081187078 | 0.290236221 | 0.139127426 | 0.089913632 | 0.365054493 | 0.66843731 | 0.098369904 | 0.057630019 | SOX4/MAFA |
| 387 | RN7SL1 | 0.231502094 | 0.299336222 | 0.56684377 | 0.269556479 | 0.206932036 | 0.213870022 | 0.290504087 | 0.219477673 | 0.27639206 | 0.275962202 | 0.222165166 | 0.292629114 | 0.189866968 | 0.161800647 | 0.112748357 | 0.394453759 | 0.402045898 | 0.329488465 | 0.348930073 | 1.164011792 | 0.395456247 | 0.147509523 | SOX4/MAFA |
| 388 | HIST2H2AA3 | 0.179647154 | 0.183943941 | 0.450600375 | 0.182720683 | 0.13102519 | 0.092296988 | 0.157319369 | 0.155654493 | 0.218703317 | 0.245847473 | 0.102838379 | 0.185418073 | 0.284272035 | 0.147681307 | 0.117261758 | 0.535498962 | 0.267992261 | 0.133603195 | 0.291553303 | 1.067998809 | 0.335671705 | 0.227191181 | SOX4/MAFA |
| 389 | C6orf120 | 0.082673752 | 0.118141078 | 0.192308939 | 0.109685425 | 0.086835238 | 0.088149787 | 0.136556631 | 0.084047323 | 0.152594847 | 0.095504444 | 0.10368217 | 0.095794689 | 0.1108176 | 0.090582683 | 0.145040925 | 0.153041555 | 0.118728064 | 0.10201214 | 0.070105245 | 0.738863024 | 0.11876212 | 0.121111767 | SOX4/MAFA |
| 390 | USP36 | 0.429138907 | 0.199368143 | 0.337135321 | 0.331187967 | 0.605921617 | 0.375257587 | 0.195721426 | 0.50585485 | 0.254151298 | 0.335580143 | 0.47388415 | 0.535583558 | 0.230578294 | 0.487565156 | 0.296917349 | 0.260946373 | 0.087029773 | 0.197716898 | 0.884990605 | 1.448123887 | 0.698573534 | 0.369972581 | SOX4/MAFA |
| 391 | CXCL10 | 0.00411408 | 0.002762922 | 0.016956349 | 0.007395945 | 0.004675795 | 0.006847638 | 0.009158262 | 0.002854688 | 0.003220484 | 0.00235871 | 0.005461341 | 0.004801768 | 0.030417898 | 0.00953004 | 0.017574592 | 0.121872658 | 0.044039603 | 0.021353215 | 0.009416666 | 0.990803569 | 0.451139616 | 0.230044678 | SOX4/MAFA |
| 392 | FAM117A | 0.224630993 | 0.194873605 | 0.34642024 | 0.181332421 | 0.181194944 | 0.146712711 | 0.157687142 | 0.152685555 | 0.20475446 | 0.180355223 | 0.158249814 | 0.221766103 | 0.128065912 | 0.124995545 | 0.133889234 | 0.276004897 | 0.18385706 | 0.127942007 | 0.10748503 | 0.620475879 | 0.091258561 | 0.07221663 | SOX4/MAFA |
| 393 | RHOB | 0.308110944 | 0.261651809 | 0.414346981 | 0.271811785 | 0.150140213 | 0.146902816 | 0.154315324 | 0.243624291 | 0.271529056 | 0.366214302 | 0.183688452 | 0.195646109 | 0.149008699 | 0.387575766 | 0.154633882 | 0.73118819 | 0.395937721 | 0.466080568 | 0.217648005 | 1.167670537 | 0.441803431 | 0.337900017 | SOX4/MAFA |
| 394 | TLE4 | 0.564599383 | 0.557044086 | 1.160935526 | 0.664321049 | 0.615423528 | 0.462311349 | 0.586251814 | 0.525327808 | 0.576160994 | 0.648838303 | 0.380035295 | 0.522674114 | 0.416511333 | 0.335341214 | 0.437309281 | 0.522120147 | 0.361492689 | 0.64144176 | 0.284032322 | 1.132510203 | 0.4110518 | 0.299479211 | SOX4/MAFA |
| 395 | HNRNPAB | 0.643172112 | 0.412395027 | 0.643414626 | 0.471713983 | 0.583923898 | 0.629043182 | 0.39867545 | 0.649053973 | 0.538768506 | 0.767994704 | 0.519384398 | 0.73119111 | 1.172705777 | 0.441330045 | 0.636498026 | 0.555558277 | 0.365290318 | 0.592368383 | 0.796905154 | 1.282580198 | 0.634993278 | 0.583208884 | SOX4/MAFA |
| 396 | IL1B | 0.020459998 | 0.019181699 | 0.050219367 | 0.017489691 | 0.020721778 | 0.03031838 | 0.021862806 | 0.017313868 | 0.018573927 | 0.015401905 | 0.019699143 | 0.019030387 | 0.022787846 | 0.018935363 | 0.015054993 | 0.028675712 | 0.022377798 | 0.094522725 | 0.074305487 | 2.045773641 | 3.154396035 | 1.113826064 | IL1B/LYZ |
| 397 | SOD2 | 0.389805184 | 0.323237025 | 0.529659364 | 0.478956843 | 0.631875991 | 0.51969649 | 0.340033971 | 0.323684689 | 0.35928753 | 0.425637636 | 0.419758114 | 0.406366989 | 0.489181067 | 0.52800626 | 0.431230024 | 0.717014126 | 0.411130906 | 0.574000807 | 0.47258514 | 2.387126985 | 3.424386892 | 2.04496043 | IL1B/LYZ |
| 398 | IL8 | 0.042889459 | 0.034859493 | 0.067785119 | 0.037595441 | 0.044318364 | 0.079771231 | 0.034141627 | 0.036247485 | 0.031349528 | 0.042638532 | 0.045449437 | 0.057376791 | 0.048242748 | 0.029814858 | 0.025544112 | 0.063908339 | 0.054468299 | 0.211496018 | 0.079778569 | 1.361200989 | 3.085683646 | 1.543460831 | IL1B/LYZ |
| 399 | G0S2 | 0.015665845 | 0.015982711 | 0.084946792 | 0.029516307 | 0.032859135 | 0.029788501 | 0.025408135 | 0.014099494 | 0.015219987 | 0.024569551 | 0.091958883 | 0.025480887 | 0.046345264 | 0.048406425 | 0.07226986 | 0.25920962 | 0.14027817 | 0.141487419 | 0 | 0.734891705 | 2.246695734 | 0.735302619 | IL1B/LYZ |
| 400 | EREG | 0.006824307 | 0.003539622 | 0.006686682 | 0.005434745 | 0.009476007 | 0.00844886 | 0.004622192 | 0.004868487 | 0.002000515 | 0.015863385 | 0.008530782 | 0.004384493 | 0.008008085 | 0.003809201 | 0.005999919 | 0.009851723 | 0.010129693 | 0.065449799 | 0.007065339 | 0.344748516 | 1.425912893 | 0.450232765 | IL1B/LYZ |
| 401 | IL1RN | 0.003409729 | 0.006149491 | 0.008213733 | 0.004600942 | 0.003234854 | 0.011588177 | 0.004366789 | 0.002306486 | 0.002333261 | 0.006280279 | 0.004544108 | 0.004740596 | 0.007327884 | 0.003415983 | 0.010367433 | 0.012519487 | 0.009292895 | 0.029465476 | 0.009777078 | 0.776617276 | 1.347372309 | 0.38580462 | IL1B/LYZ |
| 402 | S100A9 | 0.004462695 | 0.00390454 | 0.009979693 | 0.005691378 | 0.013357417 | 0.016082892 | 0.006807968 | 0.003844874 | 0.002484085 | 0.003300243 | 0.008480352 | 0.006878588 | 0.013657237 | 0.003838901 | 0.001259557 | 0.012358062 | 0.011543203 | 0.053642348 | 0.009141119 | 1.671248179 | 2.166477487 | 0.859884942 | IL1B/LYZ |
| 403 | CXCL3 | 0.006857595 | 0.005116306 | 0.023445166 | 0.005624185 | 0.007794 | 0.008301923 | 0.00686771 | 0.002630055 | 0.003886019 | 0.014087594 | 0.006169151 | 0.005904954 | 0.014752053 | 0.00488536 | 0.00365146 | 0.017460261 | 0.010003157 | 0.105153549 | 0.028181385 | 0.230608797 | 1.028225733 | 0.500970701 | IL1B/LYZ |
| 404 | PPIF | 0.155847323 | 0.114559306 | 0.18318752 | 0.178859355 | 0.21103934 | 0.232453058 | 0.0788565 | 0.176364394 | 0.169980966 | 0.162152384 | 0.203905355 | 0.318636885 | 0.268251721 | 0.171002349 | 0.137302472 | 0.220561734 | 0.122003781 | 0.176146734 | 0.095623613 | 0.465718363 | 1.61833445 | 0.59350849 | IL1B/LYZ |
| 405 | FTH1 | 2.415538541 | 2.090004223 | 1.531198098 | 2.609802406 | 3.598655804 | 2.902384698 | 2.002761321 | 2.82121001 | 2.28541028 | 2.561187898 | 2.757381756 | 2.785458011 | 2.590587404 | 2.829997078 | 2.449829531 | 2.109903271 | 1.86763585 | 3.587340231 | 3.474609855 | 3.206972887 | 5.095193948 | 4.49410914 | IL1B/LYZ |
| 406 | SERPINB9 | 0.571349867 | 0.315657359 | 0.272848978 | 0.32162047 | 0.530221442 | 0.565627234 | 0.190786052 | 0.904317193 | 0.361602078 | 0.472126311 | 0.304587665 | 0.608168383 | 0.404713596 | 0.393052243 | 0.301174854 | 0.298213671 | 0.078458119 | 0.260659651 | 0.50488273 | 0.444304909 | 1.599936766 | 0.514602302 | IL1B/LYZ |
| 407 | LYZ | 0.021209058 | 0.025059224 | 0.041030079 | 0.022480029 | 0.027569906 | 0.031596882 | 0.030492703 | 0.032593953 | 0.023785858 | 0.031416991 | 0.023187481 | 0.026624783 | 0.065024441 | 0.018426938 | 0.022402099 | 0.030000177 | 0.026811818 | 0.109819723 | 0.206589437 | 1.163232819 | 3.603665046 | 3.907100542 | IL1B/LYZ |
| 408 | TNFRSF1B | 0.479217607 | 0.3170432 | 0.147267487 | 0.207661298 | 0.440806688 | 0.990756271 | 0.172972695 | 0.443116147 | 0.204858874 | 0.406434192 | 0.476228658 | 0.544618757 | 0.234803099 | 0.113579254 | 0.032447774 | 0.135580802 | 0.026723163 | 0.091385599 | 0.313895085 | 0.398637013 | 1.194624267 | 0.537533568 | IL1B/LYZ |
| 409 | PLEK | 0.446225987 | 0.713153218 | 0.056918608 | 0.031760386 | 0.029756584 | 0.10857343 | 0.043667175 | 0.07025249 | 0.069787142 | 0.357177479 | 0.027669393 | 0.105809159 | 0.441796271 | 0.349030942 | 0.861254066 | 0.20321007 | 0.131020995 | 0.177815568 | 1.299402527 | 1.013466849 | 2.001518733 | 1.117921893 | IL1B/LYZ |
| 410 | LITAF | 0.616050926 | 0.509639338 | 0.070149088 | 0.180355398 | 0.302699189 | 0.284506042 | 0.191422719 | 0.272324934 | 0.162032727 | 0.605391047 | 0.211787269 | 0.324535668 | 0.24168092 | 0.468361944 | 0.208532064 | 0.280869879 | 0.174861393 | 0.506971538 | 0.679873729 | 0.466193519 | 1.793989647 | 1.225778712 | IL1B/LYZ |
| 411 | GPR183 | 0.759357888 | 0.347169304 | 0.718755348 | 1.136586055 | 1.837790997 | 1.361362408 | 0.54933456 | 1.083212643 | 0.532578351 | 0.639805825 | 1.11873154 | 1.066745361 | 0.332530138 | 0.930448368 | 0.192384713 | 0.570368285 | 0.262040657 | 0.426803841 | 2.931887632 | 0.406543598 | 2.142869767 | 1.138329028 | IL1B/LYZ |
| 412 | NINJ1 | 0.115568074 | 0.067246611 | 0.056289836 | 0.09334562 | 0.318661068 | 0.29987087 | 0.061700716 | 0.202067622 | 0.067056168 | 0.225244417 | 0.196840718 | 0.318176553 | 0.152227413 | 0.039223217 | 0.044985915 | 0.118387981 | 0.045409745 | 0.243466296 | 0.195452461 | 0.298293988 | 1.216327746 | 0.669826203 | IL1B/LYZ |
| 413 | PLAUR | 0.03023641 | 0.019562596 | 0.046443631 | 0.090511371 | 0.156037844 | 0.030782828 | 0.028014705 | 0.050707222 | 0.039828182 | 0.034666754 | 0.0376327 | 0.085174079 | 0.024843287 | 0.012164341 | 0.007951624 | 0.0527167 | 0.034970472 | 0.304202299 | 0.734451458 | 1.067921425 | 1.918568397 | 0.811057809 | IL1B/LYZ |
| 414 | OLR1 | 0.003471396 | 0.001457411 | 0.003476146 | 0.004888753 | 0.003729247 | 0.005547594 | 0.003032972 | 0.003740633 | 0.003916827 | 0.003110856 | 0.003776224 | 0.002798742 | 0.005224197 | 0.002666085 | 0.006831726 | 0.006161795 | 0.004607248 | 0.015128607 | 0.016498997 | 0.496867262 | 1.017928534 | 0.289712491 | IL1B/LYZ |
| 415 | IER3 | 0.274887664 | 0.253253157 | 0.549662934 | 0.345970248 | 0.147628867 | 0.104303278 | 0.221149779 | 0.202812479 | 0.440378517 | 0.437245178 | 0.110333804 | 0.275869229 | 0.165952901 | 0.068290991 | 0.060062045 | 0.495211621 | 0.193302264 | 0.628606263 | 0.106794396 | 1.42330276 | 2.330432731 | 1.150709521 | IL1B/LYZ |
| 416 | FCN1 | 0.00105626 | 0 | 0.005166119 | 0.00073888 | 0.000205062 | 0.002918997 | 0.001506884 | 0.000467331 | 0.00123777 | 0.001364406 | 0.000725851 | 0.000737178 | 0.000563611 | 0.00033243 | 0.004435642 | 0.002251608 | 0.000771469 | 0.013606655 | 0 | 0.317572651 | 0.840808799 | 0.150516237 | IL1B/LYZ |
| 417 | C15orf48 | 0.015009383 | 0.015843357 | 0.040578296 | 0.018223434 | 0.008655397 | 0.012678444 | 0.018890347 | 0.011606033 | 0.031084341 | 0.021355269 | 0.007449595 | 0.016429539 | 0.020154441 | 0.009040589 | 0.011517199 | 0.027138125 | 0.014769724 | 0.104651773 | 0.049545447 | 1.185833666 | 1.813041469 | 0.809533317 | IL1B/LYZ |
| 418 | C5AR1 | 0.002365476 | 0.0010691 | 0.00552625 | 0.001589333 | 0.00143263 | 0.003882895 | 0.002427457 | 0.001102225 | 0.001827618 | 0.004663739 | 0.001889991 | 0.00107662 | 0.003503994 | 0.001727747 | 0.007338328 | 0.002088766 | 0.002735133 | 0.005770146 | 0.008455383 | 0.153643541 | 0.831170581 | 0.533558408 | IL1B/LYZ |
| 419 | TIMP1 | 0.073229159 | 0.086825455 | 0.051713746 | 0.250203227 | 0.310599154 | 0.163541694 | 0.187721343 | 0.135192703 | 0.157383084 | 0.144921147 | 0.17954007 | 0.166099426 | 0.101274701 | 0.043691623 | 0.023520952 | 0.442942727 | 0.500728154 | 1.298757838 | 0.117405951 | 0.856699048 | 2.375994702 | 1.394569167 | IL1B/LYZ |
| 420 | CXCL2 | 0.012686048 | 0.010307627 | 0.026542658 | 0.011040712 | 0.007287194 | 0.011529134 | 0.01475435 | 0.005406449 | 0.007409284 | 0.026708227 | 0.010719666 | 0.007693005 | 0.016029552 | 0.007707476 | 0.005829085 | 0.016263551 | 0.013033244 | 0.164398038 | 0.103674831 | 0.877740255 | 1.643557614 | 0.890718565 | IL1B/LYZ |
| 421 | S100A8 | 0.000559623 | 0.002327931 | 0.004707753 | 0.004377637 | 0.003520898 | 0.007593458 | 0.000466307 | 0.001765965 | 0.002933539 | 0.002721529 | 0.002695386 | 0.003331278 | 0.001482448 | 0.001820537 | 0.001965084 | 0.00553128 | 0.002628864 | 0.030668549 | 0 | 1.319458601 | 1.465721958 | 0.376712693 | IL1B/LYZ |
| 422 | INHBA | 0.001881443 | 0.001438696 | 0.006665309 | 0.00062134 | 0.003662756 | 0.002771982 | 0.00227454 | 0 | 0.001580135 | 0.006875614 | 0.002872196 | 0.000803635 | 0.003801977 | 0.003505543 | 0.002314885 | 0.005905642 | 0.004576893 | 0.040154028 | 0.027366815 | 0.332231748 | 0.661965051 | 0.188125732 | IL1B/LYZ |
| 423 | C1QC | 0.002327398 | 0.001272769 | 0.002524205 | 0.000189841 | 0.00103632 | 0.001260169 | 0.00184914 | 0.00285375 | 0 | 0 | 0.000518843 | 0.002066954 | 0.002205044 | 0.001065497 | 0.000987148 | 0.001462397 | 0.002596226 | 0.005145101 | 0 | 0.100136244 | 0.257610415 | 2.209997034 | APOE/C1QC |
| 424 | C1QB | 0.001631959 | 0.000684389 | 0.006780844 | 0.001610872 | 0.001734837 | 0.003643447 | 0.002461809 | 0.003468228 | 0 | 0.003118444 | 0.00151288 | 0.003089389 | 0.003985839 | 0.002321097 | 0.001998104 | 0.003886914 | 0.001873551 | 0.012320712 | 0 | 0.139592553 | 0.334480485 | 2.433308442 | APOE/C1QC |
| 425 | C1QA | 0.004009219 | 0.003151036 | 0.004316375 | 0.001665136 | 0.002456273 | 0.005584373 | 0.00177437 | 0.006269226 | 0 | 0.002904108 | 0.003010369 | 0.003879201 | 0.006730828 | 0.00290352 | 0.000753392 | 0.004317403 | 0.003966549 | 0.017436477 | 0.042221668 | 0.206257182 | 0.432943463 | 2.737516492 | APOE/C1QC |
| 426 | SLC40A1 | 0.001773234 | 0.003133908 | 0.021811417 | 0.04451314 | 0.031358853 | 0.004108911 | 0.06777099 | 0.011323605 | 0.024804092 | 0.007469478 | 0.029915145 | 0.003422459 | 0.004866645 | 0.001884059 | 0.003072596 | 0.008779381 | 0.008580262 | 0.065348398 | 0.045369806 | 0.057501538 | 0.087165495 | 0.833982791 | APOE/C1QC |

| | A | B | C | D | E | F | G | H | I | J | K | L | M | N | O | P | Q | R | S | T | U | V | W | X |
|---|---|---|---|---|---|---|---|---|---|---|---|---|---|---|---|---|---|---|---|---|---|---|---|---|
| 427 | SEPP1 | 0.004387066 | 0.001741242 | 0.005285427 | 0.003053066 | 0.001792782 | 0.006422776 | 0.002774341 | 0.001570255 | 0.003792055 | 0.002240421 | 0.00369472 | 0.003932675 | 0.001102142 | 0.001945957 | 0.003066591 | 0.003109386 | 0.004899426 | 0.160404976 | 0 | 0.035140455 | 0.02369412 | 1.154293289 | APOE/C1QC |
| 428 | FUCA1 | 0.037183958 | 0.041062953 | 0.016960486 | 0.028936283 | 0.023854018 | 0.030909447 | 0.032115706 | 0.031619736 | 0.036723507 | 0.070224724 | 0.033255888 | 0.03900821 | 0.102607085 | 0.026773622 | 0.145175817 | 0.05172548 | 0.044387137 | 0.068737285 | 0.01514313 | 0.034287567 | 0.073198741 | 0.726480785 | APOE/C1QC |
| 429 | DNASE1L3 | 0.001996473 | 0.002412217 | 0.003470365 | 0.002099673 | 0.002424765 | 0.003731293 | 0.002162487 | 0.000562447 | 0.002842853 | 0 | 0.002364659 | 0.0007973 | 0.021229153 | 0.098900059 | 0.024466639 | 0.00857747 | 0.003902305 | 0.013735526 | 1.346638907 | 0.042779376 | 0.185772147 | 1.097281957 | APOE/C1QC |
| 430 | ACP5 | 0.100923861 | 0.113603275 | 0.041758028 | 0.051581216 | 0.044180702 | 0.265745044 | 0.048640585 | 0.084989756 | 0.11040044 | 0.219658471 | 0.111412206 | 0.131672684 | 0.142036624 | 0.305149872 | 0.193667767 | 0.102352567 | 0.076347572 | 0.224747155 | 0.143743972 | 0.087573903 | 0.381880711 | 1.220801591 | APOE/C1QC |
| 431 | APOE | 0.005904859 | 0.005931031 | 0.010588963 | 0.004673689 | 0.002455235 | 0.002591418 | 0.003261695 | 0.001303237 | 0.004659284 | 0.004144318 | 0.003798603 | 0.003674888 | 0.003936452 | 0.003368727 | 0.002952773 | 0.009505588 | 0.006183883 | 0.349450038 | 0.007982577 | 0.049840766 | 0.05794316 | 0.956053084 | APOE/C1QC |
| 432 | CTSC | 0.800916169 | 0.999231652 | 0.273391194 | 0.412608736 | 0.361483731 | 1.214484137 | 0.373316126 | 0.55502698 | 0.599709047 | 0.800491956 | 0.549751399 | 0.807525054 | 0.460056104 | 0.151286255 | 0.13060418 | 0.352056673 | 0.249973628 | 0.546653662 | 1.477863457 | 0.261276875 | 0.829692769 | 1.80178262 | APOE/C1QC |
| 433 | APOC1 | 0.000763679 | 0 | 0.001103605 | 0.000508023 | 0.000817117 | 0.002043388 | 0.000705506 | 0.000500979 | 0.00861815 | 0.001002691 | 0.001506094 | 0.001889671 | 0.008081839 | 0.001398046 | 0.003118017 | 0.004979002 | 0.000727081 | 0.085067437 | 0.006368161 | 0.057551931 | 0.128455093 | 0.770944133 | APOE/C1QC |
| 434 | TMSB4X | 4.493308776 | 4.670610695 | 2.604134806 | 4.411251139 | 4.472254851 | 4.232175443 | 4.296717189 | 4.5886434 | 4.728020661 | 4.561205568 | 4.481138776 | 4.544479013 | 4.492648208 | 4.204609947 | 4.660201653 | 2.711589764 | 2.0673587 | 3.789479667 | 3.511786671 | 2.346808499 | 4.215032285 | 5.051304769 | APOE/C1QC |
| 435 | FTL | 2.319982026 | 2.329138844 | 1.201242626 | 2.425447724 | 2.351704924 | 2.575706894 | 2.199302922 | 2.502580816 | 2.620674 | 2.44809224 | 2.219266712 | 2.392612029 | 2.107269435 | 2.790701591 | 2.115036798 | 3.411028845 | 3.395202889 | 3.217863908 | 3.010146892 | 2.313543999 | 4.4988226 | 5.487505917 | APOE/C1QC |
| 436 | C1orf54 | 0.007644028 | 0.002852118 | 0.002076205 | 0.009911485 | 0.010106231 | 0.006479681 | 0.011410754 | 0.010172383 | 0.010894327 | 0.010932325 | 0.004396956 | 0.010179349 | 0.013486702 | 0.007471183 | 0.005037105 | 0.007842606 | 0.005024135 | 0.087155461 | 0.044130072 | 0.017903861 | 0.172851121 | 0.731396227 | APOE/C1QC |
| 437 | NPC2 | 0.102889083 | 0.10776438 | 0.035318321 | 0.162789387 | 0.157414883 | 0.192875059 | 0.131102167 | 0.121815112 | 0.137606075 | 0.152717355 | 0.158450118 | 0.119245321 | 0.209864443 | 0.311591853 | 0.171908594 | 0.68181343 | 0.881263152 | 0.608286623 | 1.705027916 | 0.321704812 | 1.444403228 | 2.133188203 | APOE/C1QC |
| 438 | HLA-DPA1 | 0.655079104 | 0.706341173 | 0.156606915 | 0.100284564 | 0.105717555 | 0.244090655 | 0.129303497 | 0.233199752 | 0.279909688 | 0.313166296 | 0.153164298 | 0.282444376 | 1.44328239 | 2.496801943 | 2.548154247 | 0.437255428 | 0.410414328 | 0.338572041 | 2.440629506 | 0.540171875 | 2.844908177 | 3.546372834 | APOE/C1QC |
| 439 | AIF1 | 0.074829682 | 0.094922056 | 0.028586736 | 0.10685144 | 0.06797363 | 0.016049776 | 0.13676873 | 0.202654816 | 0.337064456 | 0.044233682 | 0.018095545 | 0.020272179 | 0.125860454 | 0.035900097 | 0.118019286 | 0.023564083 | 0.009562489 | 0.119410792 | 0.514979745 | 1.127990248 | 2.217784356 | 2.856553942 | APOE/C1QC |
| 440 | LIPA | 0.057020661 | 0.083077363 | 0.013535992 | 0.057426524 | 0.046607387 | 0.044806932 | 0.063864733 | 0.0574461 | 0.086543463 | 0.070855421 | 0.056626882 | 0.055359177 | 0.050920887 | 0.080023024 | 0.095560284 | 0.042496697 | 0.041016636 | 0.111684674 | 0.169618461 | 0.059414496 | 0.315404941 | 1.009031733 | APOE/C1QC |
| 441 | VAMP8 | 0.566350448 | 0.737762684 | 0.127268326 | 0.573016061 | 0.56268913 | 0.733156321 | 0.547803574 | 0.583548938 | 0.773421049 | 0.726307807 | 0.613231386 | 0.68378352 | 0.95796381 | 0.488135063 | 0.784868813 | 0.471410818 | 0.439223441 | 0.797913392 | 1.712181769 | 0.124682699 | 1.04360251 | 1.766856667 | APOE/C1QC |
| 442 | CST3 | 0.01726064 | 0.029876285 | 0.020108491 | 0.012923209 | 0.012859155 | 0.016334593 | 0.018622489 | 0.014613931 | 0.012600506 | 0.05892258 | 0.022563832 | 0.012489873 | 0.052339214 | 0.014230864 | 0.016825276 | 0.063744086 | 0.08114439 | 1.030147973 | 1.816486442 | 0.628732394 | 2.494141451 | 3.163847492 | APOE/C1QC |
| 443 | FCGRT | 0.047396612 | 0.070699221 | 0.016605248 | 0.042296355 | 0.046934032 | 0.034692567 | 0.049710269 | 0.034796963 | 0.05049581 | 0.037498342 | 0.030571654 | 0.039434489 | 0.034351128 | 0.092903593 | 0.041034602 | 0.18697975 | 0.250171316 | 0.271542637 | 0.676607543 | 0.322830474 | 0.914973547 | 1.580197895 | APOE/C1QC |
| 444 | TMEM176B | 0.002272617 | 0 | 0 | 0.001942378 | 0 | 0.003491127 | 0.003877426 | 0.000518712 | 0.000582762 | 0 | 0.000975183 | 0.001176323 | 0.002629981 | 0.001818815 | 0.004840237 | 0.002717491 | 0.001401295 | 0.538139098 | 0.009955846 | 0.082089628 | 0.503357992 | 1.230130142 | APOE/C1QC |
| 445 | HLA-DMA | 0.213732282 | 0.265961454 | 0.080587076 | 0.03244137 | 0.032190916 | 0.142881493 | 0.043735724 | 0.084719136 | 0.071476709 | 0.156103161 | 0.07156329 | 0.07424653 | 0.810274848 | 1.119763039 | 1.647618172 | 0.285891244 | 0.330644256 | 0.156971736 | 1.327915723 | 0.173366414 | 1.252997408 | 1.691150716 | APOE/C1QC |
| 446 | CD68 | 0.011023064 | 0.013719703 | 0.013905755 | 0.007970654 | 0.009106716 | 0.012061477 | 0.005952879 | 0.013921119 | 0.011045175 | 0.015614179 | 0.005086415 | 0.010022425 | 0.015298099 | 0.029631879 | 0.033459768 | 0.025190552 | 0.030443415 | 0.240827685 | 0.858277254 | 0.409696416 | 1.093127326 | 1.623272008 | APOE/C1QC |
| 447 | PLA2G7 | 0 | 0.000554209 | 0.003881625 | 0.000271608 | 0.000205062 | 0.001648154 | 0.00045855 | 0.001261766 | 0.001610116 | 0 | 0.000560791 | 0.000313605 | 0.001557402 | 0.000279894 | 0.000268441 | 0.002928186 | 0.000540736 | 0.00597222 | 0 | 0.165824904 | 0.519108686 | 1.027986748 | APOE/C1QC |
| 448 | CPVL | 0.00113236 | 0 | 0.003670808 | 0.001893652 | 0.001231926 | 0.003327009 | 0.001954304 | 0.000794973 | 0.000554484 | 0.003287483 | 0.001171236 | 0.000927781 | 0.020021365 | 0.002760633 | 0.003886841 | 0.003815175 | 0.00512299 | 0.012585912 | 0.144420107 | 0.144786238 | 0.921538153 | 1.381613969 | APOE/C1QC |
| 449 | GPNMB | 0 | 0 | 0 | 0.000837166 | 0.000524349 | 0.000491737 | 0.001249805 | 0 | 0 | 0.001474859 | 0.0002703 | 0.000894208 | 0.002858562 | 0.000598442 | 0.000662583 | 0.006317627 | 0.005176922 | 0.042607667 | 0 | 0.01968854 | 0.157196794 | 0.551529324 | APOE/C1QC |
| 450 | FCGR3A | 0.113243587 | 0.306019558 | 0.021688011 | 0.005438737 | 0.00230719 | 0.002341027 | 0.013563475 | 0.014212739 | 0.042895815 | 0.33656956 | 0.003101702 | 0.00729029 | 0.024126491 | 0.001201983 | 0.002489451 | 0.007161425 | 0.00132893 | 0.005596372 | 0 | 0.061019427 | 0.261177948 | 0.652843041 | APOE/C1QC |
| 451 | CTSD | 0.358184324 | 0.46160013 | 0.098564658 | 0.212318325 | 0.166727917 | 0.305600471 | 0.214232781 | 0.301738676 | 0.373875912 | 0.706923103 | 0.277354282 | 0.379409691 | 0.234900784 | 0.09438227 | 0.091409346 | 0.353419937 | 0.358054807 | 1.196514702 | 0.438852669 | 0.398303693 | 0.786533758 | 1.714786948 | APOE/C1QC |
| 452 | PRDX1 | 0.54879721 | 0.548501177 | 0.265978052 | 0.501464443 | 0.593621822 | 0.734501733 | 0.40749008 | 0.537877174 | 0.485373791 | 0.480129384 | 0.628343964 | 0.576876816 | 1.398183407 | 0.725254901 | 1.41361579 | 1.049591195 | 1.111973403 | 1.084480464 | 0.763097528 | 0.558067967 | 0.966227122 | 1.622283692 | APOE/C1QC |
| 453 | FABP5 | 0.619585605 | 0.432100605 | 0.153233865 | 0.101181175 | 0.166092065 | 0.213807538 | 0.104047626 | 0.268947684 | 0.179454403 | 0.362386761 | 0.553734402 | 0.490827245 | 1.032170022 | 0.186147525 | 0.529892762 | 0.197751634 | 0.077612944 | 0.321700604 | 0.408330339 | 0.12894109 | 0.602328534 | 0.990518777 | APOE/C1QC |
| 454 | MMP9 | 0.010968349 | 0.002358021 | 0.005963861 | 0.003291104 | 0.009546536 | 0.009331583 | 0 | 0.003133429 | 0.003901255 | 0.005080127 | 0.006402938 | 0.008555422 | 0.003172515 | 0.005206475 | 0.001355954 | 0.005614045 | 0.00325767 | 0.025244565 | 0 | 0.087099494 | 0.351986147 | 0.551627612 | APOE/C1QC |
