## Supplementary Table 2 for "Transcriptional Atlas of Ileal-Anal Pouch Immune Cells from Ulcerative Colitis Patients"

| Variable | **Normal pouch n=5** | **Pouchitis n=10** | **Ulcerative colitis n=13** |
|---|---|---|---|
| Sex<br>Female | <br>2 (40%) | <br>4 (40%) | <br>7 (54%) |
| Age at IBD diagnosis (median, IQR) | 36 (30-38) | 28 (13-37) | 24 (19-27) |
| Maximum extent at IPAA or biopsy<br>Proctitis (E1)<br>Left-sided colitis (E2)<br>Extensive colitis (E3) | <br>0<br>0<br>5 (100%) | <br>0<br>1 (10%)<br>9 (90%) | <br>2 (15%)<br>3 (23%)<br>7 (54%) |
| Age at IPAA (median, IQR) | 42 (39-45) | 34 (16-40) | - |
| Indication for IPAA<br>Medically refractory disease<br>Colorectal neoplasia | <br>5 (100%)<br>0 | <br>9 (90%)<br>1 (10%) | - |
| Previous medication exposures<br>Systemic steroid<br>Aminosalicylate<br>Immunomodulator<br>Anti-tumor necrosis factor<br>Anti-integrin<br>Anti- IL12/23<br>Janus kinase inhibitor<br>Antibiotic | <br>3 (60%)<br>5 (100%)<br>2 (40%)<br>4 (80%)<br>2 (40%)<br>0<br>0<br>5 (100%) | <br>9 (90%)<br>10 (100%)<br>7 (70%)<br>8 (80%)<br>1 (10%)<br>2 (20%)<br>1 (10%)<br>10 (100%) | <br>11 (85%)<br>13 (100%)<br>2 (15%)<br>9 (69%)<br>8 (62%)<br>0<br>0<br>7 (54%) |
| Medication exposure at biopsy<br>Systemic steroid<br>Aminosalicylate<br>Immunomodulator<br>Anti-tumor necrosis factor<br>Anti-integrin<br>Anti- IL12/23<br>Janus kinase inhibitor<br>Antibiotic | <br>0<br>0<br>0<br>0<br>0<br>0<br>0<br>0 | <br>3 (30%)<br>1 (10%)<br>1 (10%)<br>1 (10%)<br>0<br>1 (10%)<br>1 (10%)<br>4 (40%) | <br>4 (31%)<br>5 (39%)<br>0<br>3 (23%)<br>5 (39%)<br>0<br>1 (8%)<br>0 |
| Mayo score (median, IQR)<br>Partial<br>Endoscopic | - | - | 10 (10-10)<br>7 (7-7)<br>3 (3-3) |
| PDAI (median, IQR)<br>Clinical<br>Endoscopic<br>Histologic | 1 (1-2)<br>1 (1-2)<br>0<br>1 (1-1) | 6 (3-10)<br>2 (0-4)<br>3 (2-4)<br>1 (1-2) | - |

Abbreviations: Inflammatory bowel disease (IBD); Ileal pouch-anal anastomosis (IPAA); Pouchitis disease activity index (PDAI).
